# Diallel analysis reveals *Mx1*-dependent and *Mx1*-independent effects on response to influenza A virus in mice

**DOI:** 10.1101/214205

**Authors:** Paul L. Maurizio, Martin T. Ferris, Gregory R. Keele, Darla R. Miller, Ginger D. Shaw, Alan C. Whitmore, Ande West, Clayton R. Morrison, Kelsey E. Noll, Kenneth S. Plante, Adam S. Cockrell, David W. Threadgill, Fernando Pardo-Manuel de Villena, Ralph S. Baric, Mark T. Heise, William Valdar

## Abstract

Influenza A virus (IAV) is a respiratory pathogen that causes substantial morbidity and mortality during both seasonal and pandemic outbreaks. Infection outcomes in unexposed populations are affected by host genetics, but this host genetic architecture is not well understood. Here we obtain a broad view of how heritable factors affect a mouse model of response to IAV infection using an 8×8 diallel of the eight inbred founder strains of the Collaborative Cross (CC). Expanding on a prior statistical framework for modeling treatment response in diallels, we explore how a range of heritable effects modify acute host response to IAV through 4 days post-infection. Heritable effects in aggregate explained about 57% of the variance in IAV-induced weight loss. Much of this was attributable to a pattern of additive effects that became more prominent through day 4 post-infection and was consistent with previous reports of anti-influenza myxovirus resistance 1 (*Mx1*) polymorphisms segregating between these strains; the additive effects largely recapitulated haplotype effects observed at the *Mx1* locus in a previous study of the incipient CC (pre-CC), and are also replicated here in a CC recombinant intercross (CC-RIX) population. Genetic dominance of protective *Mx1* haplotypes was observed to differ by subspecies origin: relative to the *domesticus* null *Mx1* allele, *musculus* acts dominantly whereas *castaneus* acts additively. After controlling for *Mx1*, heritable effects, though less distinct, accounted for about 34% of the phenotypic variance. Implications for future mapping studies are discussed.

## 2. ARTICLE SUMMARY

Seasonal and pandemic influenza viruses comprise an important public health burden, but the architecture of host genetic resistance to viruses is poorly understood. We conducted an influenza challenge in a diallel cross of eight inbred mouse strains. Roughly 60% of variation in disease at 4 days post-infection was explained by genetic and non-genetic diallel effects, with about 34% of variation independent of the host resistance factor, *Mx1*. The dominance of *Mx1* resistance was highly dependent on subspecies *Mx1* allelic origin, and similar *Mx1* effect patterns were found in the related Collaborative Cross strains, motivating integrated infection studies in these populations.

## 3. INTRODUCTION

Pathogenic response to viral infection varies dramatically between individuals infected with the same viral strain and dose, and much of this variation is heritable. The impact of host genetics is evident both on the primary exposure to a virus during early life (Strunk *et al.* 2013) and upon infection with newly emerging viral strains – the latter, where prior immune exposure to a variant viral strain is not cross-protective, being especially common for quickly evolving RNA viruses such as influenza A (IAV) (Morens *et al.* 2010). Pathogenesis induced by IAV, whether contracted during early childhood or later in life, is thus likely to have a significant heritable component. A greater understanding of this heritability should improve our ability not only to identify populations at risk of enhanced morbidity and mortality during an emerging pandemic but also to identify successful options for treatment.

The past several years have seen significant progress identifying and characterizing host genes that modulate susceptibility to IAV infection, via knockout mouse studies, in vitro screens, and studies of primary immunodeficiencies and allelic variants in humans (To *et al.* 2015). In humans, screening for inborn errors identified a major role for interferon regulatory factor 7 (*Irf7*) in modulating the the severity of primary IAV infection (Ciancanelli *et al.* 2015), and allelic variation in *Ifitm3*, which was identified in a high-throughput siRNA screen, was associated with differential severity of IAV infection outcomes during the 2009 H1N1 pandemic (Everitt *et al.* 2012).

Most of our insights into genes modulating host IAV resistance, however, have come from studies on mice. These include studies using knockout mice, which have identified host genetic factors critical to antiviral responses, including *Tlr3* (Hidaka *et al.* 2006) and *Isg15* (Lenschow *et al.* 2007), and studies that examine differences between laboratory inbred strains. Inbred strain studies were the first to identify the *Myxovirus resistance* (*Mx*) family of proteins as important for host antiviral response (Staeheli *et al.* 1988), and inbred studies have continued to demonstrate the relevance of genetic background to multiple aspects of IAV pathogenesis (Srivastava *et al.* 2009; Alberts *et al.* 2010; Leist *et al.* 2016; Samet and Tompkins 2017).

Yet despite the identification of clear phenotypic differences between inbred strains, there have been relatively few attempts to dissect the genetic basis of those differences using traditional quantitative trait locus (QTL) mapping approaches such as the use of F2s or backcrosses (although see Boivin *et al.* 2012). This may be in part because traditional QTL mapping approaches tend to rely on outbred animals — and when it comes to studying viral pathogenesis, outbreds are in many respects problematic. One important limitation is phenotyping. Studying the response to an infection is equivalent to studying the causal effect of an applied treatment: its strict definition relies on a comparison between otherwise identical individuals subject to infection vs. control. But such like-for-like comparisons are biologically and technically challenging to make in an outbred population, where every individual is genetically distinct, and this has undesirable consequences for downstream interpretation: namely, that when genetic determinants of severe IAV pathogenesis are confounded with those influencing baseline phenotypes, the roles of any detected QTL are ambiguous. A related disadvantage of outbreds from the perspective of genetics is the inability to obtain biological replicates. This makes it harder to distinguish which aspects of pathology are stable consequences of genetics versus products of stochastic variability. This is particularly important, since it also makes it almost impossible to follow-up on genuinely extreme responders for additional mechanistic and genetic analysis. Translating strain differences in IAV pathogenesis to meaningful QTL studies ideally requires an experimental paradigm that combines population-level genetic diversity with individual-level replicability.

An exciting opportunity is therefore presented by replicable genetic reference populations, in particular, those based on panels of recombinant inbred (RI) strains. Across a panel of RIs, genetic variation is often more or less evenly distributed, providing a basis for QTL mapping; within a RI strain, individuals are genetically identical, providing a basis for replication. The combination allows infection response to be rigorously defined and genomic regions affecting that response to be mapped. It also permits the creation of sophisticated experiments that target a wider range of heritable mechanisms: crossing RIs with each other to form RI intercrosses (RIXs), or crossing them with outside strains, produces replicable systems capable of distinguishing, for example additive, dominance, and parent-of-origin effects, among others (Threadgill *et al.* 2002; Williams and Williams 2017).

Recombinant inbred (RI) genetic reference panels range from inbred lines derived from crosses between two mouse strains to more complex multi-parental crosses. The BxD RI panel, derived from two founder strains, has been used to study the impact of genetic variation on susceptibility to IAV infection and map QTL associated with these effects. Boon *et al.* (2009) studied H5N1 infection in females from 66 BXD strains, and Nedelko *et al.* (2012) studied H1N1 infection in 53 BXD strains, with both studies identifying QTL associated with susceptibility to infection. The Collaborative Cross (CC) RI panel is a multiparental population (MPP) descended from eight inbred founder strains (Threadgill *et al.* 2002; Churchill *et al.* 2004), with these founders including representatives from the three major domesticated house mouse subspecies (Yang *et al.* 2011). As such, the CC captures considerably more genetic diversity, and thanks to its breeding structure this diversity is also more uniformly distributed across the genome, with as many as eight distinct haplotypes segregating at any given locus within the population (Collaborative Cross Consortium 2012; Srivastava *et al.* 2017). The eight CC founder strains have distinct pathogenesis profiles in response to influenza virus (Leist *et al.* 2016), suggesting that the CC RI panel is capable of a broader phenotypic range than would be observed in less complex populations. Indeed, studies using an incompletely inbred, ancestor population of the CC (the pre-CC), demonstrated high levels of phenotypic variation across the population and successfully mapped several QTL associated with variation in susceptibility to IAV infection (Ferris *et al.* 2013; Bottomly *et al.* 2012). The CC therefore represents a promising resource for studying how genetically diverse populations respond to IAV infection.

Determining an optimal strategy for how the CC should be used to study the genetic architecture of IAV pathogenesis is nonetheless complicated because (1) the space of possible experimental designs is vast, and (2) information about what types of heritable effects are likely to be present is extremely limited. Regarding (1), with more than 70 CC strains currently available, including all reciprocal F1 hybrids (so called CC-RIXs), there are more than 4900 potential replicable configurations; since only a subset of these configurations can be explored within any realistic experiment, any chosen experimental design necessarily targets some types of heritable effects to the exclusion of others. Regarding (2), to date, most in vivo studies of IAV pathogenesis have been confined to candidate genes or additive interactions at single loci; studies investigating the broader question of what types of heritability are at play during IAV infection are largely absent.

In order to rationally design studies of heritable effects in complex populations such as the CC it is therefore helpful to have advance knowledge of which types of heritable effects might be present. One source of such information is phenotype data collected on the multiparental founders and their F1 hybrid offspring, a combination that can be more formally described as an (inbred) diallel. Diallels have long history in quantitative genetics, having been used originally in plant breeding studies to judge the relative merits of different strain combinations and subsequently for gaining insight into the heritable architecture of a broad range of phenotypes (*e.g.*, refs in Christie and Shattuck 1992; Lenarcic *et al.* 2012; Okoro and Mbajiorgu 2017), including host-pathogen interactions in, *e.g.*, crickets (Rantala and Roff 2006), flies (Wayne *et al.* 2011), and insects (Pereira *et al.* 2015) (see Methods and Discussion for connections to other diallel literature).

Here we use a diallel of the CC founders and their reciprocal F1 hybrids (hereafter, a CC founder diallel) to give an overall predictive picture of the range and relative influence of different types heritable effects on IAV pathogenesis likely to be present in CC founder-derived MPPs, a group that includes not only replicable MPPs such as the CC and the CC-RIX but also irreplicable ones such as the Diversity Outbred population (Churchill *et al.* 2012). We take advantage of the diallel design’s replicability to measure IAV-induced pathogenesis in a precise way, as the response to an applied treatment defined in terms of post-infection weight-loss differences (deltas) between matched sets of mock and infected individuals. Adapting a recently developed statistical framework for analyzing treatment-response diallels (Crowley *et al.* 2014), we use those deltas to model how pathogenic response to IAV is modulated by parentage, sex and their interaction, framed in terms of additive genetics, dominance, epistasis, parent-of-origin, and sex-specific versions thereof.

After observing that, following IAV infection, diallel individuals show a broad, continuous distribution of day 4 post-infection weight loss, we find, through statistical modeling, that the IAV-induced weight loss includes substantial contributions of host additive, epistatic, and sex-specific effects, with much of the heritable variation closely tracking the genotype state implied by the three distinct functional alleles of the previously identified susceptibility locus *Mx1*. Confirming previous findings, the functional CAST/EiJ *Mx1* allele, in contrast with functional NZO/HlLtJ and PWK/PhJ *Mx1* alleles, provides intermediate levels of protection against H1N1 influenza strains. Unexpectedly, and confirmed through additional modeling, we found that different classes of functional *Mx1* alleles exhibit distinct functional patterns, additive or dominant, when combined with null *Mx1* alleles. Further, illustrating our general rationale, we show that the major strain-specific *Mx1* effect patterns are consistent across two CC founder-derived MPPs: the pre-CC, as determined from reanalysis of a previously published dataset (Ferris *et al.* 2013), and a previously unpublished 117-line CC-RIX, in which we conduct a limited analysis focused on the *Mx1* locus.

**Figure 1.**
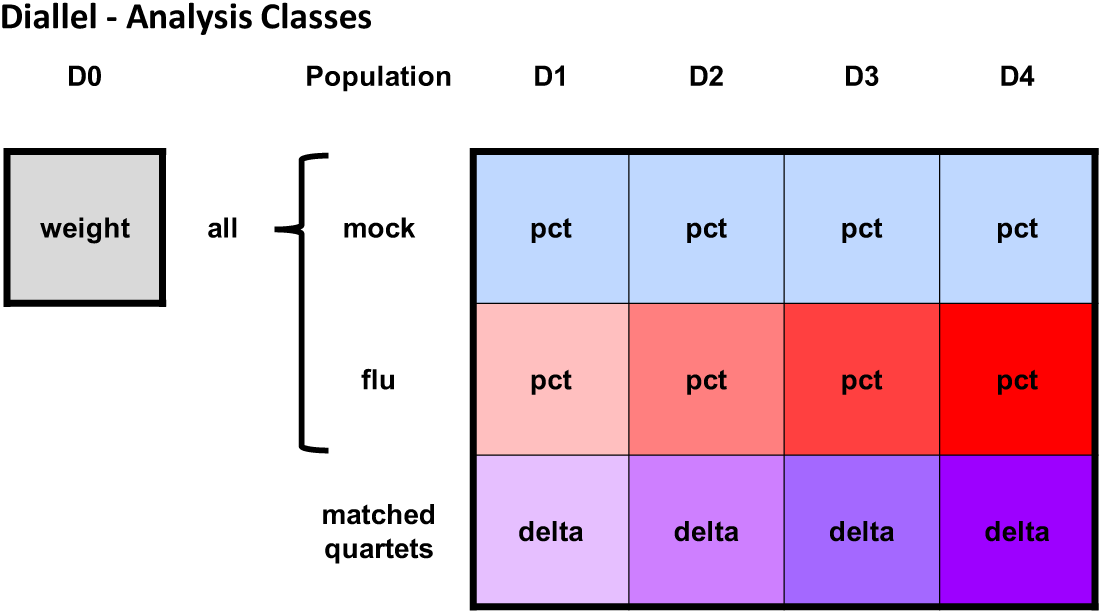
Phenotype and treatment response classes for analysis of influenza A virus infection in the diallel. Each filled square represents a weight or weight change phenotype that is modeled independently. The gray square represents the starting body weight in all animals, prior to treatment, at day 0 (D0) (analyzed with Model 1 in Table 2). Light blue squares represent animals that were mock-treated and red squares represent animals infected with IAV, with daily weights for each taken from D1 through D4 post-infection (and these analyzed with Model 2 in Table 2). Purple squares represent infection response, the primary quantity of interest, estimated using match quartets of 1 mock to 3 infected mice (analyzed with Models 3 and 4 in Table 2). Labels within each square indicate phenotypes analyzed, where weight = pre-infection body weight, pct = post-infection percent change in starting D0 weight (post), and delta = infection response, as described in the **Statistical Methods** section. The coloring increases in saturation from D1 through D4 for the influenza and matched quartet groups to indicate an overall increasing amount of post-infection weight loss over time.

## 4. EXPERIMENTAL MATERIALS AND METHODS

### A. CC founder diallel mice

The inbred and F1 mice used within this study were bred in-house at the University of North Carolina at Chapel Hill (UNC-CH). This colony was directly descended from the subset of animals used to generate the initial CC funnels (Collaborative Cross Consortium 2012), and included mice from the following eight strains at the Jackson Laboratory: A/J (“AJ”, #000646); C57BL/6J (“B6”, #000664); 129S1/SvImJ (“129”, #002448) NOD/ShiLtJ (“NOD”, #001976); NZO/HlLtJ (“NZO”, #002105); CAST/EiJ (“CAST”, #000928); PWK/PhJ (“PWK”, #003715); and WSB/EiJ (“WSB”, #001145). Mice from the UNC-CH colony were then used to generate all 62 possible inbred and (reciprocal) F1 combinations between these 8 strains, excluding NZO×CAST and NZO×PWK matings which are non-productive (Chesler *et al.* 2008) (Figure 2A). This yielded a total of 124 distinct combinations of sex and parentage (hereafter, described as “diallel categories”). Lung tissues were collected from a subset of each of the founder inbred strains in this study, at D2 and D4 post-infection, and were used for a separate comparative RNA-seq analysis by Xiong *et al.* (2014).

### B. Mouse infections in the diallel

Mice were weaned at approximately 21 days old and housed 4 per cage, within each diallel category, under standard conditions (12 h light/dark; food and water ad libitum Of the 4 mice in a cage, 1 was randomly assigned to mock and 3 to influenza infection, as there is no evidence that mice can transmit influenza virus. Each cage was then assigned to a harvest timepoint – Day 2 post-infection (D2 p.i., n=533 mice), or D4 p.i. (n=510 mice).

At 8-12 weeks of age, based on their assignments, mice were anesthetized with isoflurane and inoculated intranasally with 500 plaque-forming units (PFU) of mouse-adapted influenza A virus (H1N1 A/Puerto Rico/8/1934; short name PR8) or with the diluent, phosphobuffered saline (PBS) alone as a mock control. For each inbred line and F1 cross, about 6 mice (range: 5-9) of each sex were infected with IAV PR8, and about two mice (range: 2-3) of each sex were mock-infected, giving a total of 1,043 mice across 54 experimental batches. Treatment assignment was random: same-sex siblings from the same cage (and therefore batch) were randomly assigned at weaning to mock or infected groups prior to being moved to new cages. Body mass was recorded daily. All animal experiments were carried out in compliance with the Guide for the Care and Use of Laboratory Animals (Institute of Laboratory Animal Resources, National Research Council, 1996, https://www.ncbi.nlm.nih.gov/books/NBK232589/). Animal protocols were approved by the Institutional Animal Care and Use Committee of UNC-CH.

**Figure 2.**
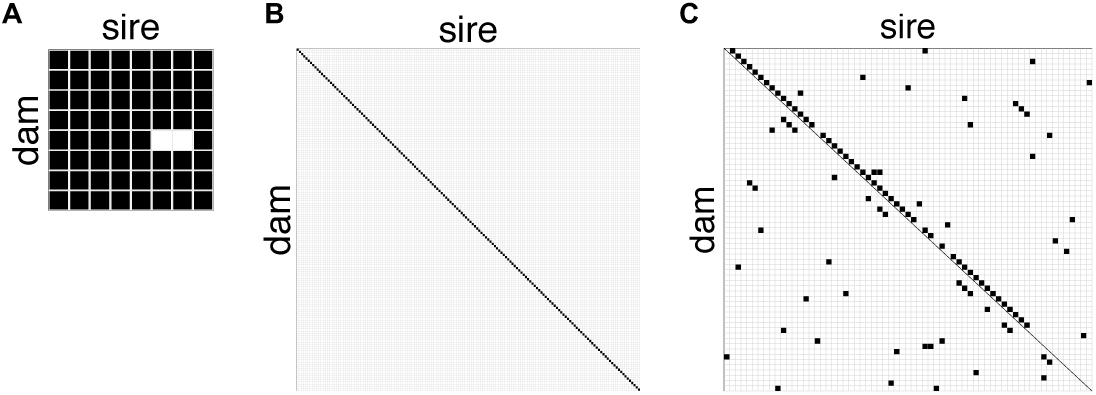
Diagram of breeding strategy for diallel, pre-CC, and CCRIX. (A) The diallel cross produces inbred (n=8) and F1 (n=54 lines) genotypes from an 8×8 cross of inbred strains. (B) The pre-CC is comprised of incompletely inbred (n=155 lines) genotypes from 155 inbreeding funnels. (C) The CC-RIX produces F1 hybrid lines (n=105 lines) from a sparse, round robin-like cross of 65 inbred CC strains.

### C. Mouse infections in the pre-CC and CC-RIX

In order to verify strain-specific haplotype effects measured in the diallel for host resistance locus, *Mx1*, we sought out CC-related IAV infection data sets for which we could isolate *Mx1* locus-specific effects.

#### C.1. Existing data from pre-CC study

In the QTL mapping study of host response to IAV infection of Ferris *et al.* (2013), 155 female pre-CC mice from as many pre-CC lines were infected with IAV (PR8) at 8-12 weeks of age and assayed for post-infection weight loss via daily weights, with phenotypes collected including starting weight (D0) and weight at D4 p.i. (Figure 2B). This study did not include mock-infected mice.

#### C.2. CC-RIX study

As part of a larger as yet unpublished study of genetic factors affecting host response to IAV, 1,402 female mice were bred from 105 F1 crosses of CC strains (*i.e.*, 105 CC-RIX lines) as described in Supplemental Material (Figure 2C, Figure S2). These mice were infected at 8-12 weeks of age with 5,000 PFU IAV (A/California/04/2009; short name CA04), a human 2009 pandemic H1N1 isolate (Itoh *et al.* 2009), and phenotypes were collected, including starting weight (D0) and weight at D7 p.i. CC-RIX were bred under similar conditions to diallel mice. This experiment, whose broader analysis is still ongoing, included both flu-infected and mock-treated mice. However, since the design did not match these to the same exacting degree as the diallel, with mock controls missing entirely for some batch/line combinations, in the current study we consider data from the infected mice only. CC animals used to generate CC-RIX lines were purchased from the Systems Genetics Core at UNC-Chapel Hill; information about CC strains available for distribution is found at http://csbio.unc.edu/CCstatus/index.py?run=AvailableLines (Morgan and Welsh 2015).

## 5. STATISTICAL MODELS AND METHODS

Our statistical analysis of heritable effects in the diallel (hereafter, diallel effects) relies heavily on the BayesDiallel model and approach described by Lenarcic *et al.* (2012) and Crowley *et al.* (2014). BayesDiallel was originally proposed in Lenarcic *et al.* (2012) for diallel analysis of routine, single-outcome phenotypes, describing how the mean value of those phenotypes was shifted by changes in parentage and sex. Although in some ways the method was built upon a canon of existing diallel literature (*e.g.*, refs in Christie and Shattuck 1992), including more recent work that used random effects (Zhu and Weir 1996; Tsaih *et al.* 2005) and Bayesian hierarchical modeling (Greenberg *et al.* 2010), in other ways it represents a new parameterization, and a generalization of many earlier methods (see Lenarcic *et al.* 2012 for explicit connections to those methods). In Crowley *et al.* (2014), we extended BayesDiallel to treatment response phenotypes, in particular, to when the modeled outcome is the phenotypic difference between placebo and treated matched pairs; the model in this case describes a causal effect modification, or, in a slight abuse of terminology, a gene-by-treatment (G×T) effect. Herein, that treatment response approach is extended further: to our more complex matching regime of quartets rather than pairs, and with a different imputation procedure to deal with quartets that are incomplete.

This section begins by reviewing the BayesDiallel model for single-outcome phenotypes. This is used not only to analyze our primary baseline phenotype, body weight at day 0 (D0 weight), but is also foundational for our subsequent analyses. Then we introduce our definition of infection response based on matched quartets, which gives rise to treatment responses defined for each of four time-points (D1, D2, D3, and D4 p.i.), and describe how they are modeled using BayesDiallel. The analysis is then modified further to estimate the impact of haplotype state at the susceptibility locus *Mx1*, and we describe how the interaction of haplotype pairs of this locus is examined by estimating relative degrees of haplotype additivity and dominance. Last we describe an illustrative comparative analysis of the effect of the *Mx1* locus on IAV response in pre-CC and CC-RIX mice.

### A. Diallel model for single outcome phenotypes

Diallel effects for single outcome phenotypes, that is, phenotypes measured as a single value per mouse, were modeled using the “fulls” model of BayesDiallel (Lenarcic *et al.* 2012; Crowley *et al.* 2014). BayesDiallel is a Bayesian linear mixed model that decomposes phenotypic variation into separate heritable components corresponding to additive genetics, dominance/inbred effects, parent-of-origin (“maternal”), epistasis, and all sex-specific versions thereof. It models the phenotype value *y_i_* of mouse *i* as

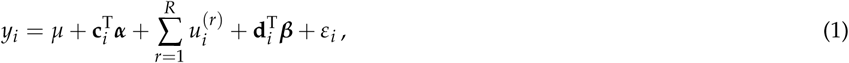

where *µ* is the intercept, and *ε_i_* is the residual error, normally distributed as *ε_i_ ~* N(0, *σ*^2^), with variance *σ*^2^. The 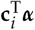 term represents the contribution of an arbitrary set of user-specified fixed effect covariates, with predictors encoded in vector **c**_*i*_ and fixed effects ***α***; the 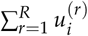 term represents the contribution of an arbitrary set of *R* user-defined random effect covariates, which for single outcome phenotypes in this study always includes an effect of experimental batch; and the 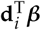 term represents the contribution of heritable components of the diallel, written as a linear combination of the diallel effects vector ***β*** and diallel category vector **d**_*i*_. Here **d**_*i*_ is shorthand for **d**_*{jks}*[*i*],_ where *{jks}*[*i*] denotes *i*’s diallel category, that is, its unique combination of mother strain *j*, father strain *k* and sex *s*. The diallel category vector **d**_*{jks}*_ is defined with the diallel effects *β* so as to give following linear combination:

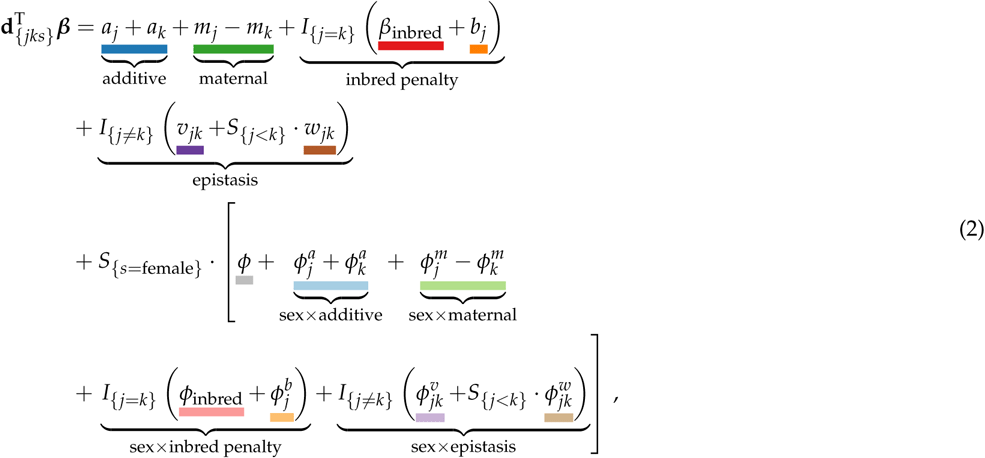

where *a_j_* is the additive effect of strain *j* (*e.g.*, the additive effect parameter *a*_AJ_ is the expected increase in phenotype on adding one haploid genome of strain AJ), *m_j_* is an additional increase in phenotype induced by strain *j* being the mother (parent-of-origin effect), indicator *I_{X}_* is 1 if *X* is true and 0 otherwise, *β*_inbred_ is the overall effect of being inbred, *b_j_* is the additional effect of being inbred for strain *j*, *v_jk_* is the additional effect of the combining strains *j* with *k* regardless of which is the mother (symmetric epistasis), indicator *S_{X}_* is 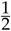 if *X* is true and 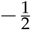 otherwise, *w_jk_* is a deviation from *v_jk_* induced by parent-of-origin (asymmetric epistasis); *φ* is the effect of being female rather than male, and 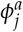 is the sex-specific deviation from additive effect *a_j_*, with other superscripted *φ* terms (*e.g.*, *φ^m^*, etc.) defined analogously. Each set of related variables, *e.g.*, the additive effects *a*_1_, …, *a_J_* for *J* parents, is modeled as a group via a constrained normal distribution, that is, *a*_1_, *a*_2_, …, *a_J_* ~ marginally 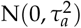, but subject to Σ_*j*_ a_*j*_ = 0, after Crowley *et al.* (2014). The variance of each group, *e.g.*, 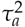 was modeled with a weak inverse gamma prior, 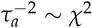, with this prior also used for the residual variance *σ*^2^. The prior for fixed effects, *e.g.*, *µ*, is set to a vague normal distribution, *µ ~* N(0, 10^3^). A summary of the diallel effects parameters is given in Table 1. Model fitting proceeded using Markov Chain Monte Carlo (MCMC) via Gibbs sampling (algorithm in Lenarcic *et al.* 2012), with results based on samples from 12.5 × 10^6^ MCMC iterations (5 chains of length 2500, after 500 iterations burnin). See also later section **Reporting BayesDiallel results.**

**Table 1.**
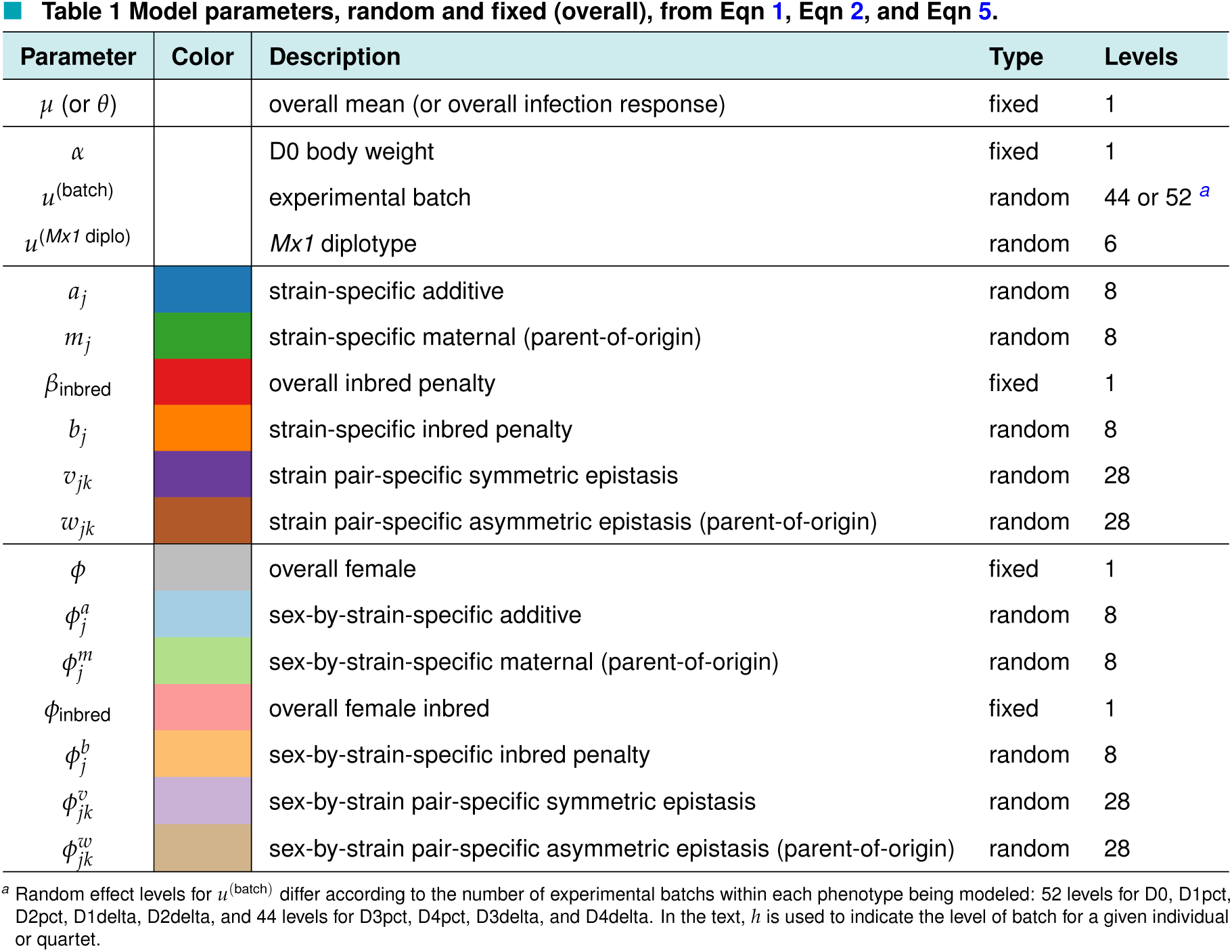
Model parameters, random and fixed (overall), from Eqn 1, Eqn 2, and Eqn 5.

| Parameter | Color | Description | Type | Levels |
| --- | --- | --- | --- | --- |
| $\mu$ (or $\theta$) | | overall mean (or overall infection response) | fixed | 1 |
| $\alpha$ | | D0 body weight | fixed | 1 |
| $u^{(\text{batch})}$ | | experimental batch | random | 44 or 52 <sup>a</sup> |
| $u^{(Mx1 \text{ diplo})}$ | | *Mx1* diplotype | random | 6 |
| $a_j$ | | strain-specific additive | random | 8 |
| $m_j$ | | strain-specific maternal (parent-of-origin) | random | 8 |
| $\beta_{\text{inbred}}$ | | overall inbred penalty | fixed | 1 |
| $b_j$ | | strain-specific inbred penalty | random | 8 |
| $v_{jk}$ | | strain pair-specific symmetric epistasis | random | 28 |
| $w_{jk}$ | | strain pair-specific asymmetric epistasis (parent-of-origin) | random | 28 |
| $\phi$ | | overall female | fixed | 1 |
| $\phi_j^a$ | | sex-by-strain-specific additive | random | 8 |
| $\phi_j^m$ | | sex-by-strain-specific maternal (parent-of-origin) | random | 8 |
| $\phi_{\text{inbred}}$ | | overall female inbred | fixed | 1 |
| $\phi_j^b$ | | sex-by-strain-specific inbred penalty | random | 8 |
| $\phi_{jk}^v$ | | sex-by-strain pair-specific symmetric epistasis | random | 28 |
| $\phi_{jk}^w$ | | sex-by-strain pair-specific asymmetric epistasis (parent-of-origin) | random | 28 |

<sup>a</sup> Random effect levels for $u^{(\text{batch})}$ differ according to the number of experimental batches within each phenotype being modeled: 52 levels for D0, D1pct, D2pct, D1delta, D2delta, and 44 levels for D3pct, D4pct, D3delta, and D4delta. In the text, $h$ is used to indicate the level of batch for a given individual or quartet.

### B. Modeling infection response as mock-corrected percent change in body weight post-infection

A standard measure used to assess pathogenesis in IAV-infected mice is weight loss. Weight loss correlates with several host and viral factors, including viral load, immune response phenotypes and lung histopathology (Ferris *et al.* 2013; Leist *et al.* 2016); as such, it provides a simple, non-invasive measure of infection pathology that can be assessed for a large number of mice. We measured the percentage change in body weight relative to D0

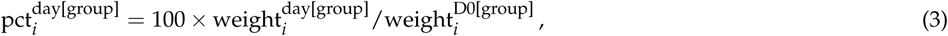

for mouse *i* on day *∈* {D1, D2, D3, D4} in group *∈* {flu, mock}, where, *e.g.*, 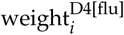 and 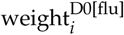 are the body weight for IAV-infected mouse *i* at D4 and at D0, respectively. These measures, which we describe as single outcome phenotypes, were analyzed using BayesDiallel as above (Table 2), but they were not the main focus of our study. Our main focus was a derived measure, IAV infection response, defined next.

In defining IAV infection response we note that from a causal inference perspective (described more fully in **Appendix A**), weight loss in an IAV-infected mouse 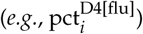 reflects two confounded processes: weight loss due to IAV-induced pathogenesis and weight loss due to other aspects of the experimental procedure. To obtain an unconfounded estimate of weight loss due to IAV-induced pathogenesis alone, we defined IAV infection response as the difference between weight loss in mice subject to infection by IAV and those subject to mock. Specifically, since in our experimental design we match one mock mouse to three infected — this reflecting our expectation that phenotypes from infected mice will be more variable and will thus need more replicates for comparable precision — infection response was defined in terms of “matched quartets”, *q* = 1, …, *Q*, where each matched quartet *q* comprised four mice of the same diallel category from the same experimental batch, with the first three mice, *q*[1], *q*[2] and *q*[3], being IAV-infected and the last mouse, *q*[4], receiving mock treatment. Infection response at a given day for quartet *q* was thus defined as a “delta”,

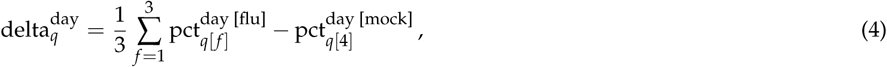

following the more general definition in Eq 11 in **Appendix A**.

Diallel effects on infection response were then modeled using BayesDiallel in manner analogous to the single outcome case in Eq 1, as

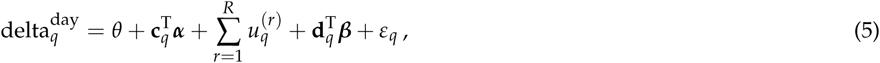

where now the unit of observation is the matched quartet *q*, rather than the individual *i*, and where, for example, **d**_*q*_ is shorthand for **d**_*{jk*,*s}*[*q*]_, the diallel category appropriate for *q*. The shift to modeling treatment response does, however, change how the parameters are interpreted. The intercept in the above formula, relabeled at *θ*, now acquires a special meaning, representing an overall causal effect due to infection, and the diallel effects in *β* now describe how that causal effect is modified by parentage, sex and their interaction; for example, the additive effect parameter *a*_AJ_ is the expected increase in infection response on adding one haploid genome of strain AJ. Regarding covariates, as for the single outcome phenotypes, this model included a random effect of batch, and, to reduce potential dependence between the deltas and baseline body weight, we also included a fixed effect covariate for the quartet mean D0 weight (ie, 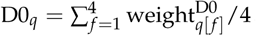) in **c**_*q*_ (Table 2).

**Table 2.**
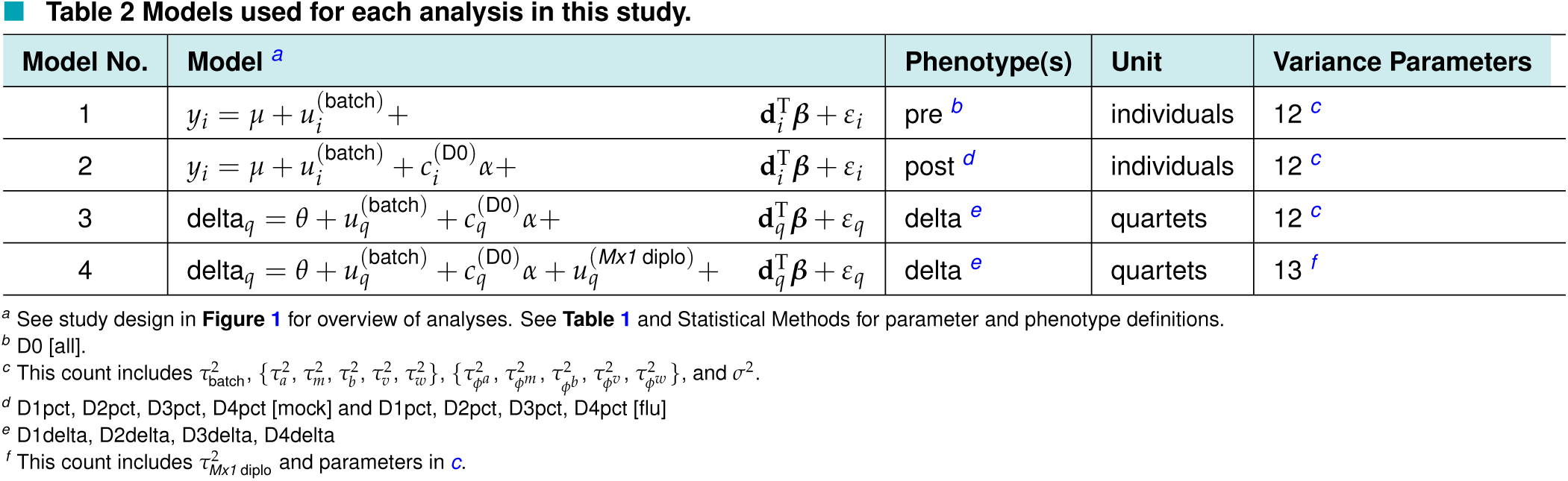
Models used for each analysis in this study.

| Model No. | Model <sup>a</sup> | Phenotype(s) | Unit | Variance Parameters |
| --- | --- | --- | --- | --- |
| 1 | $y_i = \mu + u_i^{(\text{batch})} + \mathbf{d}_i^T \boldsymbol{\beta} + \varepsilon_i$ | pre <sup>b</sup> | individuals | 12 <sup>c</sup> |
| 2 | $y_i = \mu + u_i^{(\text{batch})} + c_i^{(\text{D0})} \alpha + \mathbf{d}_i^T \boldsymbol{\beta} + \varepsilon_i$ | post <sup>d</sup> | individuals | 12 <sup>c</sup> |
| 3 | $\text{delta}_q = \theta + u_q^{(\text{batch})} + c_q^{(\text{D0})} \alpha + \mathbf{d}_q^T \boldsymbol{\beta} + \varepsilon_q$ | delta <sup>e</sup> | quartets | 12 <sup>c</sup> |
| 4 | $\text{delta}_q = \theta + u_q^{(\text{batch})} + c_q^{(\text{D0})} \alpha + u_q^{(\text{Mx1 diplo})} + \mathbf{d}_q^T \boldsymbol{\beta} + \varepsilon_q$ | delta <sup>e</sup> | quartets | 13 <sup>f</sup> |

<sup>a</sup> See study design in Figure 1 for overview of analyses. See Table 1 and Statistical Methods for parameter and phenotype definitions.
<sup>b</sup> D0 [all].
<sup>c</sup> This count includes $\tau_{\text{batch}}^2$ , $\{\tau_a^2, \tau_m^2, \tau_b^2, \tau_v^2, \tau_w^2\}$ , $\{\tau_{\phi a}^2, \tau_{\phi m}^2, \tau_{\phi b}^2, \tau_{\phi v}^2, \tau_{\phi w}^2\}$ , and $\sigma^2$ .
<sup>d</sup> D1pct, D2pct, D3pct, D4pct [mock] and D1pct, D2pct, D3pct, D4pct [flu]
<sup>e</sup> D1delta, D2delta, D3delta, D4delta
<sup>f</sup> This count includes $\tau_{\text{Mx1 diplo}}^2$ and parameters in c.

Although our experimental design stipulated even multiples of 4 mice per diallel category, practical constraints on animal breeding and availability meant that in some cases this number was 3 or 5, such that some quartets had either missing infecteds or surplus mocks. To ensure the definition of delta in Eq 4 remained consistent, and in particular that deltas from different quartets had comparable precision, the diallel analysis was performed on *M* = 1, 000 imputed versions of the data, with each imputed dataset being composed of exact quartets in which missing phenotypes had been filled using stochastic regression imputation and surplus mocks had been (randomly) deleted (details in **Appendix B**). On each imputed dataset we collected 125 MCMC samples from 12,500 total time steps (*i.e.*, by recording values at every 100th timestep); results were based on the aggregate of these samples from the *M* imputed datasets (*i.e.*, on 125,000 MCMC samples in total).

### C. Reporting BayesDiallel results: HPD, MIP, VarP and TreVarP

Point and interval estimates of individual diallel effects, *e.g.*, additive effect *a*_AJ_, are reported as posterior means and 95% highest posterior density (HPD) intervals. The overall contribution of a particular inheritance group is reported in two ways: as a Variance Projection (VarP), *e.g.*, VarP[a] for the contribution of additive effects to a phenotype or Treatment Response Variance Projections (TreVarPs), *e.g.*, TreVarP[a] for the contribution of additive effects to an infection response; and as a model inclusion probability (MIP), *e.g.*, MIP[a] for the probability of additive effects being included in the model.

The VarP is a heritability-like measure that predicts how much of the total phenotypic sum of squares would be explained by each component in a new, completely balanced diallel. Unlike traditional heritability, it is calculated based on the effects, ***β***, rather than the variance components, 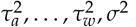, and as such benefits not only from greater interpretability but also from the stability and accuracy provided by hierarchical shrinkage (as detailed in Crowley *et al.* 2014). Since the VarP is a function of the posterior predictive distribution and calculated at each iteration of the MCMC chain, reported via Bayesian posterior summaries, specifically, the posterior mean and the *α*-level equiprobable central posterior quantile (posterior interval). The VarPs for infection response phenotypes are, following Crowley *et al.* (2014), given the special name of TreVarPs, to acknowledge their more delicate interpretation.

The MIP reflects a different type of inference: rather than being a function of the parameters estimated in the full, sexed BayesDiallel model of Eq 1 and Eq 2, it describes the results of model selection, that is, an assessment of which diallel categories could be excluded without a substantial loss in fit. As in Crowley *et al.* (2014), we use the exclusionary Gibbs group sampler of Lenarcic *et al.* (2012). Each diallel category is set to have a prior inclusion probability of 0.5, reflecting a prior opinion that inclusion and exclusion are equally likely. This prior is then updated by the phenotype data and the model selection procedure to give a (posterior) MIP. MIPs are interpreted following the conventions in Crowley *et al.* (2014): MIPs in the range (0.25,0.75) indicate that the data does not provide sufficient evidence to make an informed decision about exclusion or inclusion; MIPs within (0.05,0.25] or [0.75,0.95) represent positive evidence for exclusion or inclusion respectively; (0.01,0.05] or [0.95,0.99) represent strong evidence; and [0,0.01] or [0.99,1] represent strong to decisive evidence. These conventions are based on those proposed by Kass and Raftery (1995) for Bayes factors, which are connected to MIPs by the relation

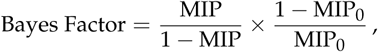

where MIP_0_ is the prior inclusion probability, and where the above simplifies to MIP/(1 *−* MIP) in our case of MIP_0_ = 0.5.

### D. Estimating *Mx1* effects in the diallel

The critical host resistance factor (*Mx1*) has been shown to drive IAV-resistance in the CC founder strains and has been mapped in the pre-CC (Ferris *et al.* 2013). *Mx1* was previously described as having three major, naturally occurring functional classes of resistance to influenza H1N1 arising from the subspecies *Mus musculus domesticus* (hereafter, *dom*; members include AJ, B6, 129, NOD and WSB), *M. m. castaneous* (*cast*; CAST) and *M. m. musculus* (*mus*; PWK and NZO), of which *dom* is considered to be null whereas *mus* and *cast* are protective. (Note that *dom Mx1* in the CC founder strains is comprised of two unique null alleles, and that the subspecific *Mx1* alleles observed in the CC may not be representative of the those segregating in the wild.) To estimate the contribution of *Mx1* haplotypes as discernible in the diallel, and thereby also estimate the extent of heritable effects that remain after *Mx1* is controlled for, we define the following haplotype combinations (diplotypes) as six levels of the random effect, *u*^(*Mx1* diplo)^: {*dom × dom*}, {*dom × cast*}, {*cast × cast*}, {*cast × mus*}, {*mus × dom*}, and {*mus × mus*}; we then repeat our diallel analysis with this effect included (Model 4 in Table 2).

#### D.1. Estimating a dominance index for Mx1 alleles

Dominance is typically defined in the context of bialleles. Since in this population, *Mx1* has a multiallelic series, we define dominance between allele pairs. Following Kacser and Burns (1981), which is built on the work of (Wright 1934), we define the “dominance index” for a wild-type (wt) against a mutant (mut) allele as

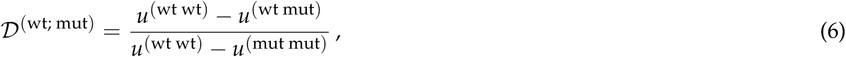

where values for 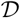 are close to −0.5 when the effect of the wild-type is overdominant to the mutant (the effect of the mutant is underrecessive), 0 when the effect of the wild-type is completely dominant to the mutant (the effect of the mutant is recessive), close to 0.5 when the effect of the wild-type is additive (not dominant, or incompletely dominant) to the mutant, close to 1 when the effect of the wild-type is recessive (the effect of the mutant is dominant), and close to 1.5 when the effect is underrecessive (the effect of the mutant is overdominant). Overdominance is given by values of 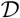 that are much less than 0 and underdominance by values that are much greater than 1. This definition is used to define dominance indices *u*^(*cast*; *dom*)^ and *u*^(*mus*; *dom*)^, describing the degree of dominance of the protective alleles *cast* and *mus* respectively against the null allelle *dom*. To assess the degree to which *cast* and *mus* differ in their relation to *dom*, we further define a “dominance difference index”,

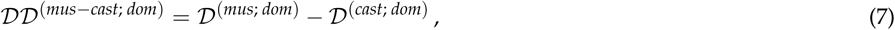

where negative values indicate that *mus* has more of a dominance-based relationship to *dom* than does *cast*, positive values indicate the converse, and zero indicates that the relationships of *cast* and *mus* to *dom* show dominance equally.

When the BayesDiallel model includes *Mx1* effects, the aforementioned dominance index and dominance difference index are both functionals of the posterior; posterior samples of these indices were therefore obtained by simply applying Eq 6 and Eq 7 to the sampled *Mx1* effects at each timestep of the MCMC chain.

The Kacser and Burns (1981) dominance index is a simple re-parameterization of the degree of dominance parameter, *a*_CR_, defined by Comstock and Robinson (1948) and used by Gardner and Lonnquist (1959). In the Comstock and Robinson model, the mean centered phenotypes are coded, translating our model above, as: *u*^(wt; wt)^ = *w*, *u*^(wt; mut)^ = *aw*, and *u*^(mut; mut)^ = *−w*. This gives the relation 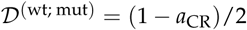 or equivalently, 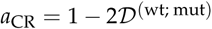.

### E. Estimating haplotype effects at the *Mx1* locus in the pre-CC and CC-RIX

The additive effect parameters estimated in the diallel do not precisely distinguish the effects at the *Mx1* locus because they are confounded with any potential effects genome-wide that follow the same pattern of strain classification. An unconfounded estimate of haplotype effects at *Mx1* requires a population in which the remainder of the genome is randomized, *e.g.*, by recombination. To this end, we make use of two related data sets on IAV-induced weight loss in two CC-derived MPPs: IAV (PR8) infection in the pre-CC and IAV (CA04) infection in a set of CC-RIX lines. These two studies, described in more detail below, were in other respects less rigorous than our diallel: the experimental measurement of the infection response was based on infected mice only with no mocks in the pre-CC, and although mocks were collected in the CC-RIX, their relative sparsity (200-300 mocks to >1400 infecteds) complicates analysis based on matching alternate treatment groups; the experimental batching was subject to a less exacting degree of randomization across genetically distinct categories; the available combinations of *Mx1* diplotypes are limited mostly to homozygotes in the pre-CC, and incompletely and unevenly sampled in the CC-RIX; the *Mx1* diplotype state for each line is known only probabilistically, having been inferred by hidden Markov models (HMMs) applied to genotyping data. Nonetheless, if effects at the *Mx1* locus were largely independent of those elsewhere in the genome, we might expect that *Mx1* effects in the pre-CC and CC-RIX would be broadly consistent with those in the diallel.

Estimation of haplotype effects at the *Mx1* locus was performed using the Diploffect model (Zhang *et al.* 2014), a Bayesian hierarchical model that estimates effects of diplotype substutions at a specified QTL when the diplotype states themselves are known only probabilistically. The effects estimated by Diploffect are analogous to those estimated by BayesDiallel: phenotype *y_i_* of mouse *i* is modeled as

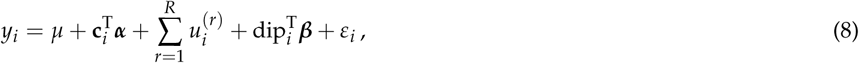

where dip_*i*_ is a vector representing the diplotype state of mouse *i* at the QTL and is shorthand for dip_*{jk}*[*i*]_, where *{jk}*[*i*] denotes *i*’s diplotype state composed of haplotypes from CC founder strains *j* and *k*, *β* are the corresponding effects, and all other variables are as in Eq 1. The diplotype vector dip_*{jk}*_ is defined with *β* so as to give the linear predictor

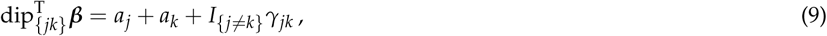

where *a_j_* and *a_k_* are additive (haplotype) effects modeled as 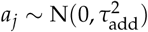, broadly equivalent to the additive effects in BayesDiallel’s Eq 2, and 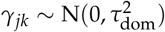 are dominance deviations, which are the converse to BayesDiallel’s inbred parameters. Dominance deviations are expected to be poorly informed when heterozygotes are sparsely represented, as in the CC-RIX and in particular the largely inbred pre-CC, but are nonetheless included to stabilize inference of additive effects. For numerical stability, phenotypes were first centered and scaled to unit variance, and variance parameters (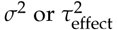, where effect is add, dom or *r ∈ R*) were given mildly informative priors of the form 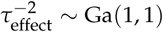. Estimation proceeded by importance sampling (the DF.IS and DF.IS.kinship methods in Zhang *et al.* 2014) using integrated nested Laplace approximations (INLA; Martins *et al.* 2013), with 100 importance samples taken, and parameter estimates for additive effects are reported as posterior means, posterior medians and HPD intervals.

#### E.1. Pre-CC study

In the study of Ferris *et al.* (2013), IAV infection response was measured on 155 mice from as many pre-CC lines as weight loss following infection with IAV (PR8 variant, as for the diallel). QTL mapping of D4 p.i. weight loss, equivalent to 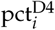 in the diallel study, identified a QTL, *HrI1*, containing the *Mx1* gene, with peak marker JAX00072951 (chr16:98,148,641; Mouse Diversity Array of Yang *et al.* 2009). We estimated haplotype effects at this peak marker using Diploffect (Zhang *et al.* 2014), applied to the phenotype and the original HMM probabilities of Ferris *et al.* (2013), with the model including a fixed effect covariate for D0 weight.

#### E.2. CC-RIX study

For the CC-RIX study of infection response to IAV (CA04 strain), we calculated weight loss values for all 1,402 infected mice at D7 p.i. (analogous to a 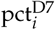 measure), and for all 105 CC-RIX lines obtained diplotype probabilities at marker UNC27478095 (16:97,591,482; MegaMUGA array, described in Morgan *et al.* 2016) from the Inbred Strain Variant database (ISVdb; Oreper *et al.* 2017). Haplotype effects were then estimated by Diploffect applied to debatched CC-RIX line means as follows. First, we fit linear mixed model (by REML using R package lme4 of Bates *et al.* 2015) to the individual-level phenotypes (n=1,402) with fixed effects of D0 weight and lab (2 levels), and random effects of mating (107 levels: 105 RIXs + 2 additional levels distinguishing minor breeding differences, when CC010 and CC042 strains were re-derived from breeder females into a new facility) and infection date (59 levels). The residuals of this model were then averaged over the *n_i_* mice of each CC-RIX line *i*, and used as the response *y_i_* in Eq 8 with precision-weighting *ε_i_ ~* N(0, *σ*^2^/*n_i_*) and a between-line polygenic random effect 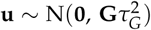, where 105 × 105 genetic relationship matrix **G** was calculated between all CC-RIX pairs based on the founder haplotype probabilities (dosages) at each locus, according to the method described in Gatti *et al.* (2014).

### F. Availability of Data and Software

Analyses were conducted in the statistical programming language R (R Core Team 2017). In addition to R packages cited above, we used the packages BayesDiallel (Lenarcic *et al.* 2012) and Diploffect.INLA (Zhang *et al.* 2014). The data, analysis software, and scripts are available on flu-diallel repository on GitHub, at https://github.com/mauriziopaul/flu-diallel. A static version is posted as a public, open access Zenodo repository, at http://dx.doi.org/10.5281/zenodo.293015. Phenotype data from the diallel and CC-RIX animals used in this study will be available on the Mouse Phenome Database (Grubb *et al.* 2014), at https://phenome.jax.org with persistent identifier RRID:SCR_003212.

File S1 contains an account of the supplemental files which can be used to reproduce our analysis. File S2 contains the software packages used for this analysis. File S3 contains the diallel data file, and File S4, S5, and S6 contain the data analysis files required for analyzing the diallel, pre-CC, and CC-RIX, respectively. After unzipping, the files FluDiData.csv, Flu-pre-CC-data.csv, and Flu-CC-RIX-data.csv contain raw phenotypes, cross (or line, strain), and mouse ID information from the three mouse populations used in this study. The script files MIMQ*.sh are used in bash to call R scripts to run the BayesDiallel analysis on diallel phenotypes. The script files main_analysis*.R are used with Diploffect to run Diploffect analysis on the pre-CC and CC-RIX phenotypes. Additional *.RData, *.pl, *.alleles, and *.csv files are uploaded which contain settings, genotypes, and founder haplotype probabilities used by the scripts.

## 6. RESULTS

**Figure 3.**
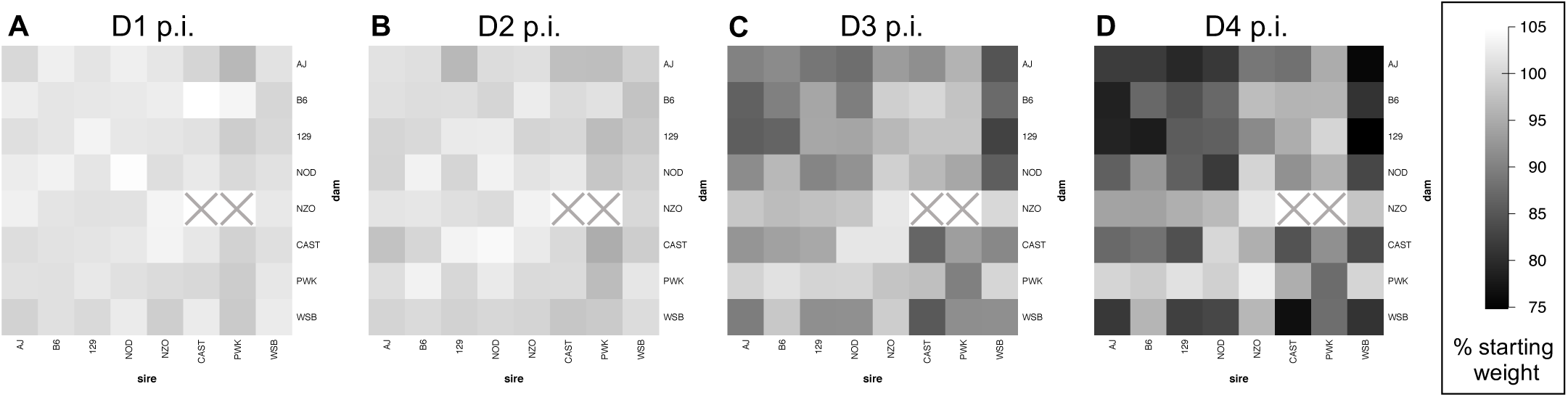
Influenza-induced weight loss in an 8×8 diallel cross of mice, through four days post-infection. Mean weight change, as % D0 weight, is shown at (A) day 1, (B) day 2, (C) day 3, and (D) day 4 post-infection with 500pfu IAV (PR8) in male and female inbreds and F1 hybrids of CC founder strains (n=774 for D1 and D2, n=382 for D3 and n=381 for D4). Results from mock-infected mice not shown. Squares with a grey "X" indicate matings that do not produce offspring.

Mice from the eight inbred founder strains of the CC were used to generate a near-complete 8 × 8 diallel. This study used offspring (n=1,043) of both sexes (519 females, 524 males) representing 62 of the 64 crosses (Figure S1), including all inbred combinations (n=129) and all F1 hybrids (n=914) except NZO×CAST and NZO×PWK. Within each diallel category, defined as the combination of sex and (reciprocal) parentage, and in each experimental batch, mice were randomly assigned at weaning to infection or mock groups in a ratio of 3:1; complete sets of 3 infected with 1 mock were described as matched quartets. Mice in the infected group were inoculated with IAV PR8 and in the mock group with PBS. For each mouse, body weight was measured prior to infection (D0 or baseline weight), and at days 1-4 post-infection (D1, D2, D3, D4). D0 weight is reported in grams whereas post-infection weight is hereafter reported as a percentage of D0 weight, *e.g.*, D4pct. Not all mice survived the protocol: one infected mouse died after D3 weights were taken and one mouse died from anesthesia on D0.

**Figure 4.**
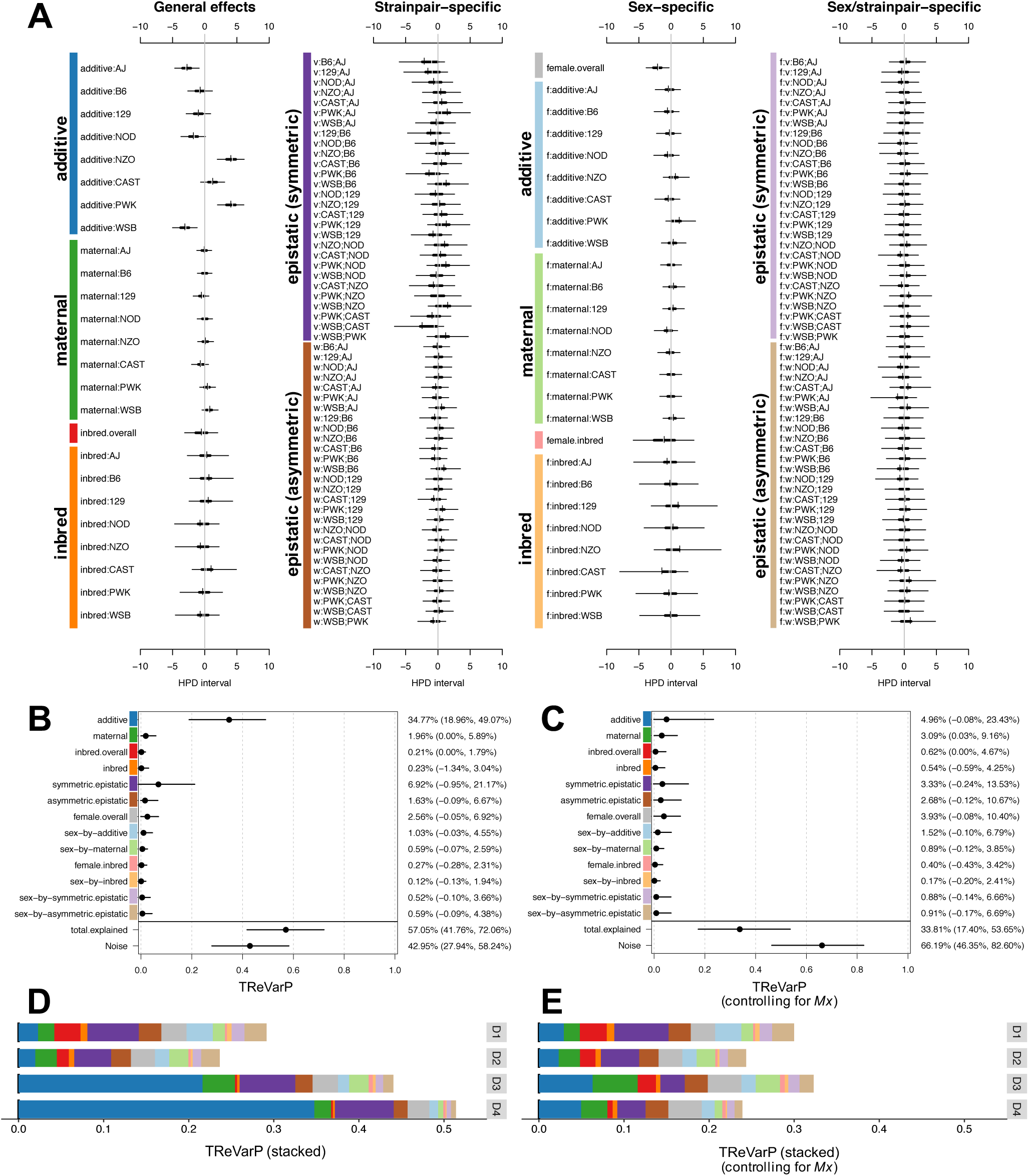
Diallel effects on host weight IAV-infection response, before and after accounting for *Mx1* haplotypes. A) Effect estimates for additive, maternal, inbred, and epistatic effects, including sex-specific effects, are presented as highest posterior density (HPD) intervals across 163 individual effects categories for IAV-induced weight change at D4 p.i. (phenotype D4delta). HPDs are given for each parameter, including 95% (thin line) and 50% (thick line) intervals, and median (white break) and mean (black vertical line). Parameters are labeled according to the methods. Symmetric epistatic, asymmetric epistatic, and sex-specific parameters are indicated by “v:”, “w:”, and “f:”, respectively. The overall treatment effect, *θ*, (not shown) is −8.85% (−9.92%, −7.78%). B-C) Treatment Response Variance projections (TReVarPs), a generalization of heritability for diallel effects classes, at D4 are shown for three fixed (overall) effects, five random effects classes and five corresponding sex-specific random effects classes (posterior median and 95% quantile-based confidence intervals) before (B) and (C) after accounting for diplotypes of the host influenza resistance allele, *Mx1*. D-E) TReVarPs before and after *Mx1* for all four post-infection timepoints.

### A. F1 hybrids of the CC founders show a wide range of phenotypic outcomes

The CC founders include five strains we have previously characterized as susceptible to IAV-induced pathology (AJ, B6, 129, NOD, and WSB), two strains as resistant (NZO and PWK), and one (CAST) that exhibits a distinct intermediate weight loss phenotype (Ferris *et al.* 2013). Results for the inbred founders measured in our diallel replicate those earlier findings, and the post-infection weight loss among the infected F1 hybrids spanned the range of phenotypes observed in the founders (Figure S3), consistent with the notion of IAV-induced weight loss being a complex trait with contributions from multiple loci.

### B. Diallel effects on baseline mouse weight strongly replicate previous CC founder diallel studies

The effects of parentage and sex on D0 weight were estimated using BayesDiallel. Described further in **Methods**, BayesDiallel decomposes the heritable effects observable in the diallel into 160 parameters (diallel effects) grouped into 13 distinct heritability classes. In sketch form, it models the average phenotype of mice of sex *s* bred from mother of strain *j* and father of strain *k* as

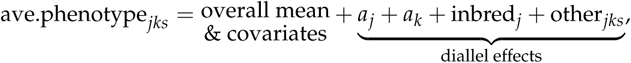

where covariates always includes experimental batch, *a_j_* and *a_k_* are the additive effects of the two parents, inbred_*j*_ is an additional effect included only when *j* = *k*), and other_*jks*_ models the effects of further nuances of sex and parentage as deviations from this base model (listed in **Methods** and Table 1).

Diallel effects estimated for D0 weight are reported in Figure S6A as 95% HPD intervals for each parameter, and two summary measures, VarPs and MIPs, for each of the 13 heritability classes are given in Figure S6B,C. Briefly, VarPs (Figure S6C) report the contribution of the effect group as the proportion of the total phenotypic variance, whereas MIPs (Figure S6B) assess the strength of support for whether an effect group should be included at all, with probabilities near 1 providing stronger support for inclusion, probabilities near 0 supporting exclusion, and probabilities near 0.5 reflecting a lack of information either way.

The pattern of effects for D0 weight was strikingly similar to that seen for baseline body weight in two previous diallels of the CC founders (Lenarcic *et al.* 2012; Crowley *et al.* 2014), despite those earlier studies being independent experiments with no particular attempt made to align experimental protocols, and included substantial additive effects, strain-specific parent-of-origin effects, signals of epistasis, and sex-specific versions thereof. For example, we largely replicated the pattern of inbred, additive, and maternal effects observed in both Lenarcic *et al.* (2012) and Crowley *et al.* (2014), and also found a higher-order sex-specific PWK×CAST symmetric epistatic effect in Lenarcic *et al.* (2012). We also observed some new epistatic and sex-specific epistatic effects largely due to increased power from a larger sample size.

### C. Diallel effects on IAV infection response

Infection response was defined as the percentage change in body weight induced by IAV infection, with more negative values indicating more severe pathogenesis. This was calculated at each timepoint, D1, D2, D3, and D4 p.i., as the difference between matched infected and mock mice, yielding a single infection response number (a “delta”, *e.g.*, D4delta) for each matched quartet (3 infected mice and 1 mock). The effects of parentage and sex on infection response were then analyzed for each timepoint separately using BayesDiallel as above, with an additional covariate of D0 weight (see **Methods** for details). Although results are provided in Supplemental Materials for all timepoints, we will focus on those for D4 p.i. since this showed the greatest difference between infected and mock.

#### C.1. IAV infection causes weight loss through day 4 post-infection, with greater susceptibility in females

IAV infection in the diallel induced an overall mean change in body weight (*i.e.*, overall infection response *θ* in **Eq** 5 and Table 1) of −0.13% (95% HPD interval: −0.48, 0.22; MIP=1) on D1 p.i., −0.83% (−1.33%,-0.32%; MIP=1) on D2 p.i., −5.60% (−6.47%, −4.73%; MIP=1) on D3, and −8.85% (−9.92%, −7.78%; MIP=1) on D4 (Table S3; see also progression in Figure 3). Consistent with previous mouse studies of sex effects on infection (Robinson *et al.* 2011; Lorenzo *et al.* 2011), females given the same dose of virus as male mice had increased weight loss: a negative effect of female sex was estimated at all four time points p.i., gradually increasing in magnitude from −0.89% (−1.45%, −0.36%) at D1 p.i. to −2.11% (−3.87%, −0.30%) at D4 p.i. (Figure 4), suggesting that enhanced susceptibility in females may occur at least as early as D1 p.i. Although all mice received the same dose of virus regardless of starting body weight, heavier mice experienced a transient increase in percent weight loss at D2 p.i. compared with lighter mice: the D0 weight effect (*α* in Eq 5) on the infection response at D2 p.i. was −0.31% (−0.52%, −0.09%), such that for every 10 grams of starting weight beyond 0 grams, an additional *≈* 3.1% weight was lost on D2; however, this effect disappeared by D3 p.i. No other significant effects of starting weight on IAV-induced weight loss were detected at other time points, indicating that heavier mice were infected at least as effectively as lighter mice, and that starting body weight does not in general confound our exploration of strain and cross-specific effects.

#### C.2. Diallel effects on infection response reflect mostly additive genetics, consistent with differences in Mx1 haplotype

Infection response in our diallel was strongly driven by additive effects. On D3 p.i., enhanced susceptibility to weight loss in infected animals was affected most by contributions from strain AJ, −2.17% (−3.72%, −0.61%), and enhanced resistance from contributions of NZO, 2.54% (0.72%, 4.27%), and PWK, 1.70% (0.12%, 3.23%), strains. On D4 p.i., enhanced suscepitibility was greatest from AJ, −2.77% (−4.66%, −0.86%) and WSB, −3.09% (−5.01%, −1.18%), with enhanced resistance greatest from NZO, 4.07% (1.95%, 6.12%), and PWK, 4.06% (1.97%, 6.08%) (Figure 4A). In terms of its additive effect, CAST was more resistant than the *Mx1*-null strains (AJ, B6, 129, NOD, and WSB) but about half as resistant as the *Mx1*-functional strains (NZO and PWK), consistent with it conferring intermediate protection in the heterozygote state.

To summarize these effects: for each dose of AJ or WSB genomes inherited from a parent, about 2%-3% of additional starting body weight is lost post-infection, indicating enhanced susceptibility compared with the overall mean weight loss; for each NZO and PWK genome inherited, about 4% more of starting body weight is retained post-infection, compared with the mean treatment effect, indicating enhanced resistance.

Diallel effects explained over half of the total variance of infection response at D4, with a treatment response VarP for all effect groups collectively of 57% (TReVarP[all]=0.571; 0.418, 0.721). The variance explained by additive effects only, which is related to the narrow-sense heritability, was estimated as 34.8% (TReVarP[a]=0.348; 0.190, 0.491), and also detected were potential additional contributions of epistasis (TReVarP[v]=0.069; −0.001, 0.212) and maternal effects (TReVarP[m]=0.020; 0.000, 0.059) (Figure 4B,D, Table S1).

#### C.3. Evidence for additive, inbred, epistatic and parent-of-origin effects mounts as disease progresses

The relevance of diallel effects to infection response became more marked with time (Table S3, Figure S7-S10). At D1 and D2 p.i., model inclusion probabilities gave strong support only to an overall infection response, with no evidence of this effect being modified by sex or parentage (Figure S7-S8). At D3 p.i., however, we found positive to strong evidence of additive (MIP[a]=0.978), inbred (MIP[b]=0.958), and asymmetric epistatic (MIP[w]=0.820; *i.e.*, parent-of-origin epistatic) effects (Figure S9). By D4 p.i., support for additive (MIP[a]=0.998) and inbred (MIP[b]=0.999) effects had become decisive (see **Methods** for MIP interpretation), and there was strong support for both symmetric epistatic (MIP[v]=0.960) and asymmetric epistatic (MIP[w]=0.966) effects (Figure S10, Table S3).

### D. Modeling effects consistent with *Mx1* haplotype

To help distinguish diallel effects that are consistent with the subspecies haplotype of the resistance factor *Mx1* (hereafter, *Mx1* effects), we incorporated the *Mx1* subtype explicitly into the model as a genotype covariate with three alleles, one for each subspecies branch: *dom* (AJ, B6, 129, NOD, WSB), *cast* (CAST), and *mus* (NZO, PWK).

#### D.1. *Mx1* effects are increasingly evident with disease progression; explain ~40% of the diallel effects at D4 p.i

In keeping with the increased support seen for diallel effects over time, evidence for a non-zero *Mx1* effect increases from positive evidence of exclusion on D1 (MIP=0.035) to no evidence for inclusion or exclusion on D2 (MIP=0.552), to decisive evidence for inclusion on D3 (MIP=1.000) and D4 (MIP=1.000) (Figure S11–S14); a comparable level of support for inclusion in the model was seen only for effects of overall treatment and batch. After controlling for *Mx1*, the variance explained by diallel effects at D4 was substantially reduced, from 57% to 33.8% (TReVarP[all|*Mx1*]=0.338; 0.174,0.537) (Figure 4C,E, Table S2). This was consistent with *Mx1* accounting for about 40% of the variance explained by the diallel, including most of the additive effects (mathematically the *Mx1* term models effects that compete with a subset of the additive and dominance diallel effects).

**Figure 5.**
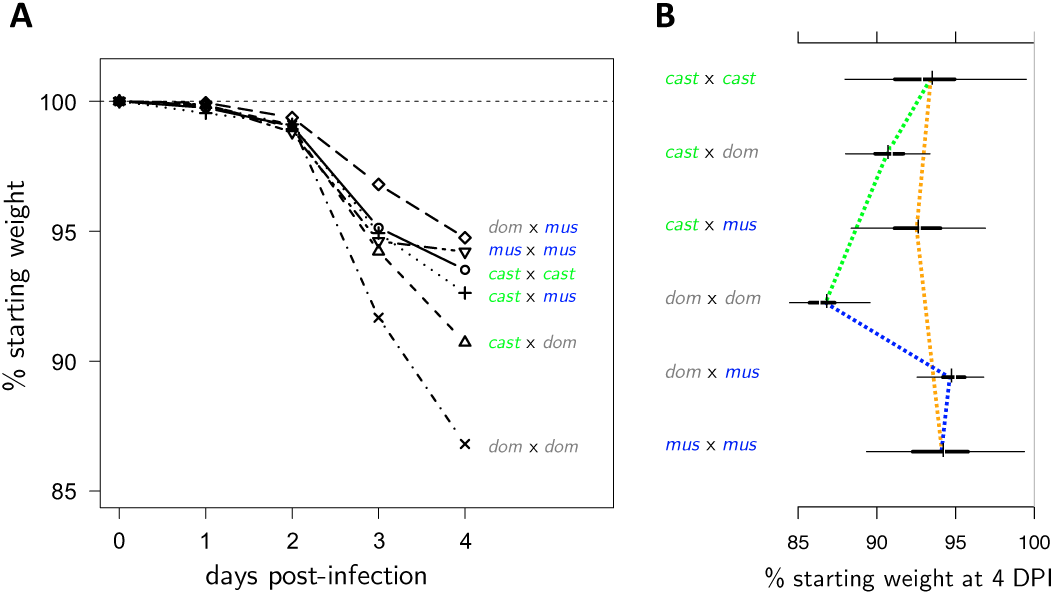
Time course of subspecies-specific *Mx1* haplotype effects on IAV-induced weight change in the diallel. A) Predictive means of *Mx1* diplotype effects across four days post-infection, modeled simultaneously with other diallel effects and covariates. B) HPD intervals of *Mx1* diplotype effects on weight change on day 4 post-infection. Increased resistance is indicated by values further to the right. Dashed lines highlight the mode of interaction between *Mx1* haplotypes: green 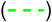 shows the additive effect of crossing *cast* with *dom*, blue 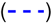 the dominant effect of crossing *mus* with *dom*, and orange 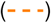 the neglible effect of *cast* crossed with *mus*.

#### D.2. Evidence for distinct additive and non-additive effects of Mx1 functional groups

After controlling for other diallel effects, the predicted weight loss over the course of 4 days varies in a manner consistent with *Mx1* allele combination (Figure 5A). We observed that, as expected, *dom × dom* crosses were predicted to have much more overall post IAV-infection weight loss at D3 and D4 compared with all other crosses. Notably, the most protected group appeared to be the *dom × mus* haplotype, at both D3 and D4 post-infection, although the HPD intervals overlap with other *Mx1*-functional groups. The rank order of effects changes from D3 to D4 due to the dramatic slowing of weight loss in the *mus × mus* crosses from D3 to D4 compared with D2 to D3.

Although we did not observe any strain- or pairwise-specific non-additive effects in the diallel prior to inclusion of the *Mx1* random effect, we did observe a pattern of dominance in crosses between *mus* and *dom*, even as there was a pattern of additivity in the crosses between *cast* and *dom* (Figure 5B). Whereas it might be expected that host alleles from *Mx1*-null strains should act in a recessive manner, this appears not to be the case for this phenotype and time point in crosses of *cast* with *dom*, such that the functional *Mx1* allele from CAST appears to operate in an additive manner. This further supports the previous observation that the CAST *Mx1* alleles differ from the *mus Mx1* alleles in their protective host response to IAV (Ferris *et al.* 2013).

#### D.3. Dominance and additivity of Mx1 alleles against the functional null: mus is dominant, cast acts additively

To better characterize how the *Mx1* effects on infection response exhibit aspects of genetic dominance vs. genetic additivity, we estimated for each functional *Mx1* allele a “dominance index”, after Kacser and Burns (1981). This measures the distance between the expected phenotype of a homozygous functional allele, in our case *mus* or *cast*, and the heterozygote formed with a null allele, in our case *dom*. On this scale, 0 denotes the functional allele being dominant to the null, 1 denotes it being recessive and 0.5 indicates pure additivity (see x-axis scale in Figure 6A, B, and more details in **Methods**).

**Figure 6.**
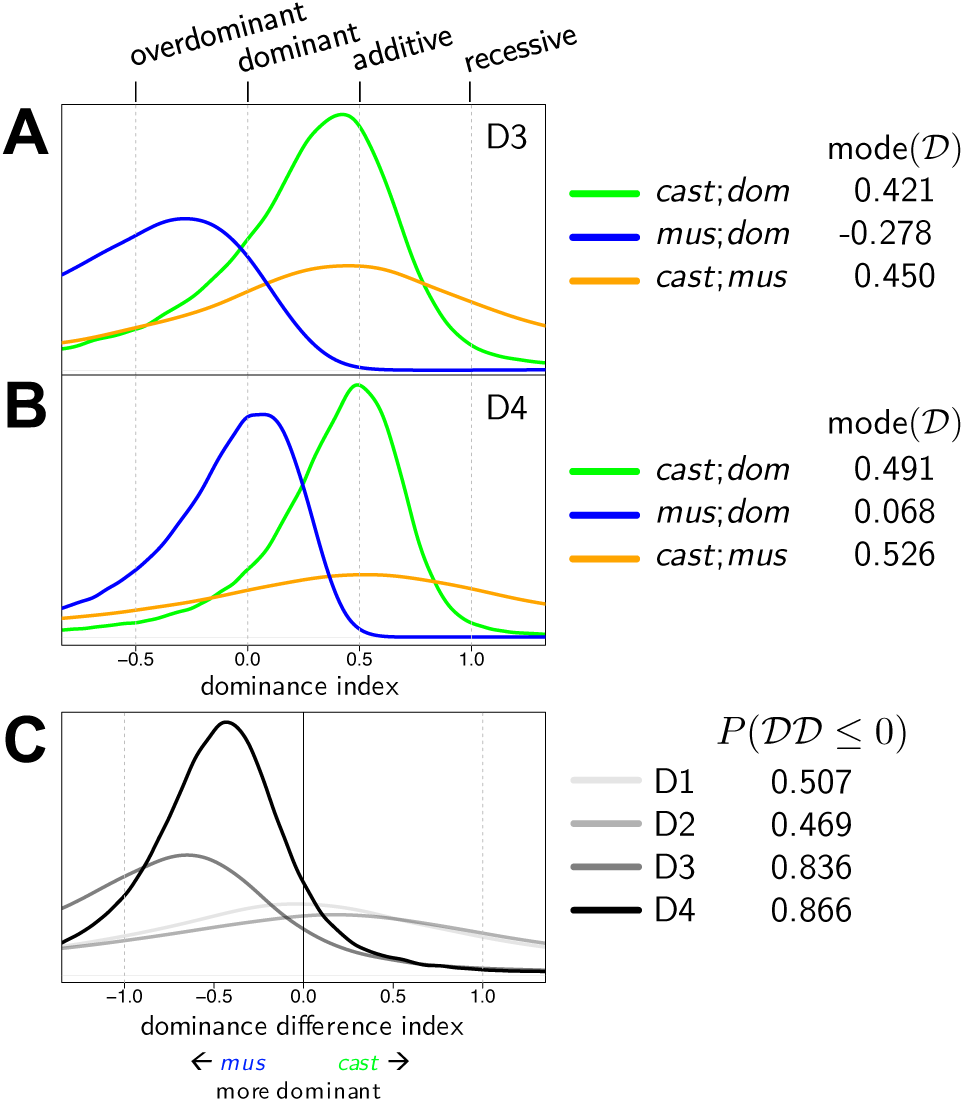
Posterior density of the dominance index on (A) D3 and (B) D4. C) Posterior density of the dominance difference index, *i.e.*, the difference between the dominance indices of *cast* and *mus*, across all 4 days.

The dominance indices of the two functional *Mx1* alleles, *mus* and *cast*, were sharply different (Figure 6A,B; Table S8). We found that *mus* against *dom* was −0.278 (=posterior mode of 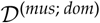; 80% HPD interval −2.547, 0.329) at D3 and 0.068 at D4 (−0.568, 0.380), a clear signal of *mus* exerting classical dominance over the functional null. In contrast, the dominance index of *cast* against *dom* was 0.421 (−0.534 to 0.907) and 0.491 (−0.028,0.836) for D3 and D4, consistent with *cast* and the functional null being codominant (*i.e.*, having an additive relationship). The difference of the two dominance indices, whose posterior distribution is shown in Figure 6 for each timepoint, quantifies the distinction between *mus* and *cast* more directly, putting the probability that *mus* is more dominant than *cast* (*i.e.*, 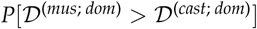) at 83.6% for D3 and 86.6% for D4.

**Figure 7.**
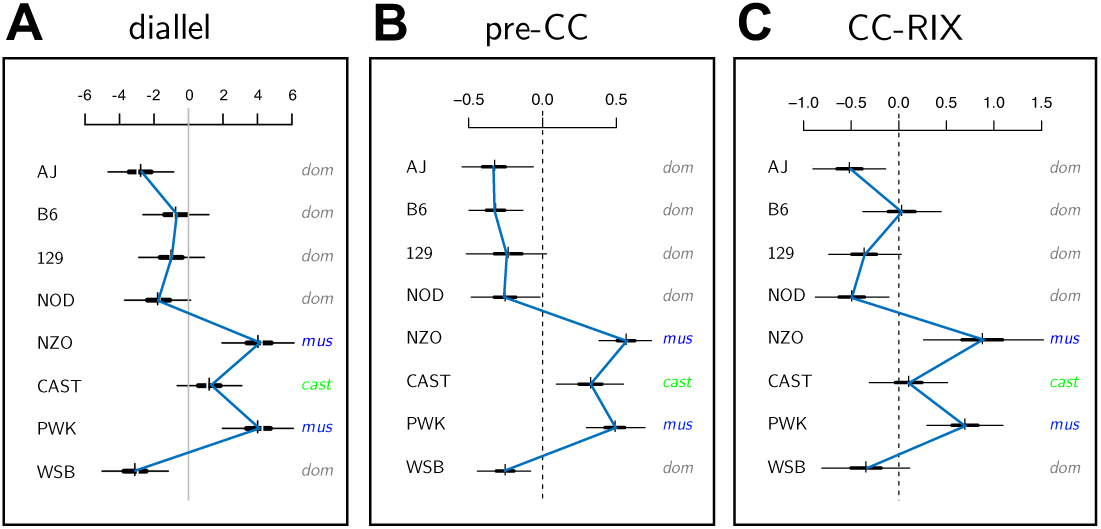
Additive CC-strain haplotype effects on IAV-induced weight loss across three CC-related populations. A) Additive effects from the CC founder diallel of mice infected with IAV (PR8) or mock virus (n_flu_=393, n_mock_=131) at day 4 p.i. (from Figure 1). B) Additive strain haplotype effects at the *Mx1* locus for female pre-CC mice (n=155) infected with IAV (PR8) at day 4 p.i. C) Additive strain haplotype effects at *Mx1* for female CC-RIX mice (n=1,402) infected with IAV (CA04) at day 7 p.i. Estimates are shown as HPD intervals as described in Figure 1, with blue lines connecting posterior means. Parameter scales are given as additional IAV-induced weight loss per dose of strain in % of (A) D0, and (B, C) normalized effect size.

### E. *Mx1* effects show consistent pattern in related multiparent populations, pre-CC and CC-RIX

We examined effects associated with the *Mx1* locus in two related recombinant CC populations, the pre-CC of Ferris *et al.* (2013) and a set of CC-RIX lines first described here, and observed that the pattern of locus-specific strain haplotype effects was strikingly similar to that observed in our diallel Figure 7). This suggests that the pattern of genome-wide additive effects in the diallel is largely driven by the effect of *Mx1* haplotypes in the founder strains. This similarity in pattern is consistent, even though the virus isolate and the peak weight loss timepoint differed in the CC-RIX population (CA04 human pandemic strain, D7 p.i.) compared with the diallel and pre-CC (PR8 mouse-adapted strain, D4 p.i.) (Table S4). In all three populations, NZO and PWK alleles provide the most resistance to IAV-induced weight loss, and CAST alleles are slightly less protective. In the pre-CC, effects of AJ, B6, 129, NOD, and WSB haplotypes are all approximately the same, and clearly separated from the additive effects of strains with functional *Mx1*. In the diallel and in the CC-RIX (at *Mx1*), however, AJ and WSB haplotypes are on average more susceptible than the B6 haplotype, and there is less separation between additive effects of CAST and those from from *Mx1*-null strains. The proportion of variance in weight loss explained by *Mx1* was estimated as 0.5 (95 % HPD interval: 0.43, 0.54) and 0.54 (0.42, 0.63) for pre-CC and CC-RIX mice, respectively (Figure S16 and Figure S17). Note that an in-depth analysis of dominance indices for the *Mx1* locus was not possible in these populations owing to the relatively sparse coverage of heterozygote diplotype states in the pre-CC and homozygous functional diplotype states in the CC.

## 7. DISCUSSION

We describe a general approach for investigating heritable effects on host susceptibility to virus-induced disease—in our case pathogenesis induced by IAV—using a diallel cross of the eight CC founder strains. The results from this diallel are informative not only in more clearly defining genetic architecture of the host influenza response, but also prospectively: they anticipate sources of heritable variation likely to be present in the CC, the DO and other derived experimental populations, and therefore provide a ready basis for the rational design of future studies. As an illustration of this, we demonstrate concordant effects of viral resistance locus *Mx1* across the CC founder diallel, pre-CC and a set of CC-RIX lines.

With regard specifically to IAV pathogenesis, our study sought to better understand host genetic effects on this outcome in terms of their (1) time-dependence, (2) consistency across related populations, and (3) conditionality; for example, dependence on interactions between alleles at the same locus (dominance, at *Mx1*) or at different loci (epistasis). Regarding time-dependence (1), we found that, whereas the effect of being female rather than male is evident from day 1, the effects of genetics appear later, becoming evident only on day 3 and then increasing through day 4 post-infection. Regarding consistency (2), we found that the effects of the *Mx1* alleles seen previously in the CC founders remain stable across inbred, F1, and recombinant populations. Regarding conditionality (3), we found something unexpected: evidence that the two *Mx1* functional classes, *casteneus* (CAST) and *musculus* (NZO and PWK), which were previously characterized as being functional alleles, in fact behave differently when present in the heterozygous state with susceptible *Mx1* alleles from *domesticus* (AJ, B6, 129, NOD, WSB). Specifically, the protection conferred by the presence of a *musculus Mx1* allele is the same regardless of whether it is in the homozygote state or paired as a heterozygote with the null *domesticus* allele; the *musculus* allele is therefore dominant to *domesticus*. But for the CAST allele, when paired in the same way with *domesticus*, its protection is weakened, to an extent consistent with CAST and *domesticus* being codominant, that is, having an additive relationship.

### A. The level of resistance to IAV among different inbred mice is conditional on IAV subtype and strain

Differences in *Mx1* function have been identified between a variety of inbred mouse strains, including the CC founders (Ferris *et al.* 2013; Xiong *et al.* 2014; Leist *et al.* 2016). Our results were largely consistent with those studies.

Notably, in their examination of the CC founders with H3N2 infection, Leist *et al.* (2016) identified AJ and WSB strains as being most susceptible, and NZO and PWK as being most resistant, which agrees with our diallel additive effects. However, in contrast with our results showing partial protection against H1N1 IAV with CAST *Mx1*, which is consistent with our prior findings in the pre-CC (Ferris *et al.* 2013), they found CAST mice, grouping with AJ and WSB, to be highly susceptible. This difference could arise for at least two reasons. First, across the influenza field, even in identical RI panels (Boon *et al.* 2009; Nedelko *et al.* 2012), host genetic effects appear to be IAV subtype-specific. Second, the effectiveness of *Mx1*’s antiviral activities can vary depending on IAV subtypes (Riegger *et al.* 2015; Dittmann *et al.* 2008; Zimmermann *et al.* 2011; Mänz *et al.* 2013; Verhelst *et al.* 2012). Differentiating these two possibilities, however, is beyond the scope of this work.

Although the molecular differences in CAST *Mx1* that produce a deficient response in comparison with *mus Mx1* have not been defined, some work has been done in inbred mice to better understand CAST/EiJ-(strain)-specific antiviral responses. In order to interpret what they saw as a unique antiviral deficiency of CAST mice, transcriptomic experiments by Leist *et al.* (2016) suggested enhanced susceptibility is due to leucocyte recruitment deficiency (relative to NZO and PWK) in the lung. In the CC founder study of Xiong *et al.* (2014), several transcriptomic differences separated the CAST response to PR8 from the that of the other strains, including differential splicing of *Irak1* and lack of *Ifng* expression at D4 p.i., which was consistent with *Ifng* deficiency observed by Earl *et al.* (2012) leading to lethal monkeypox infection of CAST mice. Because these studies were completed in inbred CAST mice, the role of CAST *Mx1* is confounded with the genome-wide differences between CAST and the other CC founders.

Thus, there are several challenges to understanding the unique IAV resistance profile of CAST *Mx1* based on existing studies: (1) studies in inbred lines are unable to probe the overall or *Mx1*-specific dominance architecture due to a lack of heterozygosity; and (2) studies in non-recombinant lines that identify a unique phenotype in CAST compared with other founders are unable to separate the effect of CAST *Mx1* from effects arising from the rest of the CAST genome. Our study in part circumvents these shortcomings by: (1) additionally examining F1 hybrids, and (2) exploring the emerging phenotypes from an ongoing IAV infection screen using CC-RIX, themselves F1s of RI strains.

### B. Complex additive effects patterns mask strong signals of dominance

In our initial analysis, we found that most of the phenotypic variation explained in infection response is driven by additive genetics with no particular signal of dominance. However, when we explicitly modeled *Mx1* status, using a term that competes with a subset of the additive and dominance diallel effects, we found that the *Mx1* functional classes act in a manner consistent with a strong dominance pattern for *musculus Mx1* (Figure 5). It seems striking that such a pattern of dominance could be underlying an apparently heavily additive effect signal.

Identifying dominance requires a good basis for comparing inbreds with hybrids. However, since the diallel is mainly composed of F1 hybrids with relatively few (8 vs 54) inbreds, this basis for comparison is often weak. The BayesDiallel model handles this by considering the hybrid state as the baseline and treats the inbred state as the exception (a deviation) relevant to a minority of categories, as discussed further in Lenarcic *et al.* (2012). Inferred dominance effects are therefore vague because the data that informs it is sparse, and low estimates of dominance variance comes from absence of information rather than from information about the absence of an effect. Nonetheless, greater precision was available when considering dominance of substrain-specific *Mx1* because dominance information was pooled across multiple strains and strainpairs.

The fact that the proportion of estimated additive vs. non-additive variance is influenced by model parameterization motivates careful consideration of both study design and analysis. As Huang and Mackay (2016) have recently described, model parameterizations can have critical effects on the detection of non-additivity, with the same data strongly supporting evidence for mostly additive or mostly non-additive effects, depending on the model. Related issues have been described at the locus level by Sabourin *et al.* (2015), who showed that when applying penalized regression to multi-SNP fine-mapping in GWAS, genotype parameterization interacts with how priors/penalties are assigned and can make biallelic dominance hard to identify in some cases. Yet even when dominance is not of interest per se, failure to accommodate it can disrupt estimation of additivity: in the pre-CC QTL mapping study of Phillippi *et al.* (2014), dominance signals arising from residual heterozygosity disrupted detection of an additive QTL for basal levels of CD23 (encoded by *Fcerii*); this was resolved by treating heterozygote diplotypes, whose occurence was too sparse to be modeled, as inherently noisier via downweighting.

### C. Antiviral genes are expected to be dominant, but CAST *Mx1* exhibits additivity

The degree of genetic dominance of host resistance factors to viral infection in humans and mice has not been thoroughly explored. In general, in the context of biochemical and immunological studies one might expect, just as with *musculus Mx1* combined with *domesticus Mx1*, that genes encoding strong-acting antivirals when combined with a null mutant would be mostly dominant. In quantitative genetics, however, it is more often expected that genetic contributions will be mostly additive. In this study, at the *Mx1* locus, we observe both.

In genetic crosses of functional and null mice, major host determinants of pathogenesis are might normally be expected to be classified as either recessive or dominant: recessive when null results in loss-of-susceptibility for a host factor required for disease susceptibility; dominant when null results in loss-of-function for a host gene required for virus resistance. The recessive case is especially true of passive immunity gained by knockout of host genes critical to viral entry and life cycle, and has been demonstrated in a variety of studies on crop resistance (Fraser 1990; Kang *et al.* 2005; Truniger and Aranda 2009; Hashimoto *et al.* 2016), and explored in studies of the effects of CCR5 deficiency (CCR5-delta32 deletion) in resistance to HIV infection and pathogenesis in humans (Samson *et al.* 1996; Liu *et al.* 1996; Hütter *et al.* 2009), however the degree of protection in the CCR5-delta32 heterozygous individuals is not fully understood (Marmor *et al.* 2001; Trecarichi *et al.* 2006). The dominant case could be considered for a viral sensor, where a single inherited functional copy still provides sufficient sensitivity for viral detection and control, resembling that of an individual inheriting two copies, one from each parent. This type of dominance is best explained by the model proposed by Kacser and Burns (1981), a metabolic-enzymatic model for the architecture of dominance at specific loci, and has been explored further in studies of viral resistance in plants, such as in Fraser and Loon (1986) and Fraser (1992). The Kascer-Burns model also provides a mechanism that could in some cases give rise to additivity.

Kacser and Burns (1981) predicted that, biochemically, for most enzymes, if there is a 50% reduction in enzyme activity in the heterozygote of a null × functional cross, then in most cases the resulting phenotype will resemble that in the homozygous functional individual and the null allele would likely be characterized as operating in a “recessive” manner. According to their model, the phenotype (or “flux”) resulting from a given enzymatic pathway with multiple enzymes joined by “kinetic linking” is a summation of the change in flux due to each specific enzyme activity (“selectivity coefficient”). This means that even a dramatic change in activity for any one enzyme in a physiological system results in barely discernible changes in the system overall, as long as some functional enzyme from the locus of interest is produced.

However, the authors also describe two cases where systemic flux can be partially reduced in the heterozygote: (1) in pathways where there are exceptionally few enzymes involved in the system (this case is unlikely for an IFN-responsive antiviral pathway such as *Mx1*); and (2) in pathways where the selectivity coefficient (functional activity) of the enzyme is very low, a case termed heterozygote “indeterminacy,” which we henceforth equate to additivity. As further explored by Keightley (1996), dominance may be incomplete when less active allelic members of a series are involved in a cross with null mutants, resulting in a more additive relationship; this seems most likely to explain our observation of CAST *Mx1* effects, and the lower antiviral activity of CAST *Mx1* observed in Nürnberger *et al.* (2016), discussed below, appears to support this.

### D. Recent work exploring CAST *Mx1* antiviral deficiency

Important insights into why CAST *Mx1* might be additive come from recent functional studies. Nürnberger *et al.* (2016) engineered B6 mice expressing either the CAST-derived or A2G-derived MX1 proteins. A2G encodes an MX1 protein sequence similar to the NZO and PWK *musculus* class described in this study. CAST MX1 differs from A2G and *musculus*, with corresponding amino acid changes G83R and A222V in the G domain, which is important for enzymatic and antiviral function. Nürnberger *et al.* (2016) clearly show that CAST provides intermediate protection from IAV, in their case using H7N7 (SC35M) and H5N1 (R65) viruses, and suggest that sequence changes in the CAST *Mx1* allele result in reduced enzyme stability, metabolic instability, and possibly in altered dimerization of MX1 monomers and/or changes in MX1 GTPase antiviral activity. It is unknown whether the differences they observed would lead to changes in the dominance of CAST and A2G *Mx1*, although we might expect this to be the case given our mouse infection results. We have verified that the same variants, G83R and A222V, differentiate CAST coding sequence from NZO and PWK, as in Srivastava *et al.* (2009) and using http://isvdb.unc.edu (Oreper *et al.* 2017), and that these are the only nonsynonymous variants on coding transcripts of *Mx1* that differentiate CAST from NZO and PWK. Although we see substantial protection from weight loss in CAST mice, we see a deficiency in the anti-viral effects (as measured by RNA-seq viral reads in infected lungs) of CAST *Mx1* on D2 and D4 post-infection (data not shown, via RNA-seq reads from Xiong *et al.* 2014, and transcript analysis in Ferris *et al.* 2013). Our work motivates further functional studies of the MX1 protein using *Mx1* transgenic mice.

### E. *Mx1*-independent effects and their follow-up: new studies should leverage CAST *Mx1* additivity

A substantial proportion of heritable variance in the diallel was *Mx1*-independent (VarP[all*|Mx1*]=33.81, Table S2). This was broadly driven additive genetics and both symmetric and asymmetric epistasis (*i.e.*, differing by parent-of-origin) (Figure 4C, Figure 4E). Relatedly, in our analysis of the *Mx1* locus in the CC-RIX, we estimated *Mx1*-independent effects attributable to overall genome-similarity to account for 21% of phenotypic variance. Both observations suggest the presence of additional QTL that could be drawn out given a suitable follow-up design.

Consider the design of a second CC-RIX. Here our knowledge of differences in *Mx1* dominance becomes a valuable guide: prioritizing CC F1s with one copy of *musculus Mx1* would reduce power because it would cause *Mx1*-independent drivers to be masked; however, prioritizing CC F1s with one or fewer copies of *castaneous Mx1* would leave the *Mx1*-independent effects exposed and QTL underlying them more easily detected.

The inclusion of mice with a single functional *Mx1* in a mapping population provides a basis for mapping loci that modify the effect of *Mx1*, as well as mapping *Mx1*-independent loci controlling disease. Shin *et al.* (2015) showed that even the protectiveness of *Mx1* from the A2G inbred strain is conditional and depends on host genetic background. Thus, CC-RIX designs that incorporate heterozygous classes of *domesticus Mx1* crossed with either CAST *Mx1* or *musculus Mx1* can be of substantial benefit for mapping novel loci affecting infection outcomes, and at least 40% of the F1 crosses in our CC-RIX study incorporate lines which have one single copy (CAST or *musculus*) of *Mx1*.

### F. Diallels as pilot data more generally

Diallels have a long history in quantitative genetics (Schmidt 1919; and refs in, eg, Christie and Shattuck 1992; Verhoeven *et al.* 2006; Lenarcic *et al.* 2012), and have most commonly been used as a way to assess the relative potency of different genomes with respect to a studied trait, yielding, for example, estimates of generalized combining ability (GCA) for each strain and estimates of specific combining ability (SCA) for each F1. More ambitiously, they have been used to obtain an overall picture of a trait’s genetic architecture. In many respects, this picture is clearly incomplete: even within the limited genetic space spanned by the founders, the diallel shows only the effects of swapping intact haploid genomes, with no ability to see the effects of recombination. But in other respects it is comprehensive: in considering every F1 combination, one can observe evidence for types of effects — dominance, epistasis, parent-of-origin, epistasis by parent-of-origin, all sex-specific versions thereof — that would be hard or impossible to identify in other settings, *e.g.*, outbreeding populations derived from the same set of founders.

A number of studies have sought to combine the features of a diallel with those of such derived outbred crosses to obtain picture of genetic architecture that is in some way informed by both. These include studies that map QTL across multiple biparental (*e.g.*, F2) crosses derived from a diallel or diallel-like population (*e.g.*, Rebai and Goffinet 1993; Xu 1998; Rebaï and Goffinet 2000; Liu and Zeng 2000; Ogut *et al.* 2015), and at least one theoretical study, that of Verhoeven *et al.* (2006), examining the extent to which such information can be analyzed jointly and reconciled with data from the original diallel itself.

The goals of our study were more prospective: we use the diallel to prioritize follow-up designs in target populations that segregate genetic material from the same set of founders; the diallel provides evidence of heritable features that would be expected to exist in the CC, and that could be examined in more detail in a suitably designed CC-based experiment. A comprehensive view of IAV resistance architecture, even within the genetic space of the CC-founder genomes, would be achievable only asymptotically through countless, diverse studies; in this, the diallel can be seen as a compass, identifying promising initial directions.

### G. Potential for joint analysis

Despite our diallel focus, since we do examine data from two other closely related populations it is nonetheless interesting to consider the potential value of analyzing all three populations jointly. In theory, a joint analysis could, in bringing greater numbers, sharpen estimates of key parameters representing effects that are shared across the studies while also serving to more precisely characterize key differences between them — the idea being that such formal consideration of similarities and differences would lead to a more nuanced picture of the examined trait’s genetic architecture, in particular the relationship between QTL and overall effects (see for example related discussions in Holland 2007).

A related hope would be that the effects estimated in the diallel could increase power to map QTL in the pre-CC and CC-RIX, for example, by serving as priors in a Bayesian model of the QTL allelic series (such as that proposed by Jannink and Wu 2003).

Our preference against a joint analysis in this case centers on the fact that the three studies differ not only in their genetic configuration, which if alone might make such an analysis justified, but also in other important respects that would make formally-estimated between-study discrepancies hard to interpret and potentially misleading.

First, the studies differed in their phenotype definitions. Infection response in the diallel was defined rigorously, in terms of mock and infected mice, but in the pre-CC and CC-RIX its definition was looser, lacking mock controls; although the effects for *Mx1* across populations turned out to be consistent, any discrepancies identified in a joint model would have been hard to attribute to differences in genetics vs. differences in protocol.

Second, the studies differed in the IAV variant used. In the pre-CC and diallel we used PR8 but for the newer CC-RIX experiment we switched to CA04. Our reason for this change is that CA04, a mouse-adapted virus from a 2009 H1N1 clinical isolate, infects mice with a lower rate of mortality, making it a better candidate for QTL-mapping studies of IAV pathogenesis, and, being a human virus, CA04 is more amenable to translational studies.

Nonetheless, absent these conflationary factors, a joint analysis could be valuable, and we expect that joint analyses of heritable effects across populations are likely to be particularly apt when comparable experiments are repeated across subsets of the CC-founder diallel, the DO, the CC, and CC-RIX, or more generally across different components of a multiparent super-population.

### H. Summary

Our study demonstrates the use of diallel crosses for identifying different types of heritable effects that can affect host responses to IAV infection. As such, we find reproducible effects of *Mx1* alleles across first order crosses and recombined populations (despite inexact coordination between protocols), confirming our previous findings that the CAST *Mx1* allele exhibits an intermediate resistance phenotype against H1N1 strains of influenza virus (Ferris *et al.* 2013), and also identifying novel attributes of the CAST and *musculus Mx1* alleles with respect to additivity and dominance. Despite a body of literature on the effects of null mutations in *Mx1*, the importance of allelic variation at this antiviral gene is just beginning to be understood. A GWAS study published in 2011 found that *Mx1* allelic variation likely plays a role in viral disease manifestation in humans, specifically with regards to West Nile virus infection (Bigham *et al.* 2011), highlighting a need for further study of the role of natural allelic variation in *Mx1* on virus infections in future research.

## 9. ACKNOWLEDGMENTS

We thank Dr. Peter Palese, at the Icahn School of Medicine at Mount Sinai, NYC, NY, for provision of mouse-adapted IAV H1N1 PR8 viral stocks (A/Puerto Rico/8/1934) for the diallel and pre-CC infections. We thank Dr. Yoshihiro Kawaoka, at the University of Wisconsin Department of Pathobiological Sciences, for providing the IAV H1N1 CA04 (A/California/04/2009) infectious clone plasmids that were used to generate viral stocks for the CC-RIX infections. We thank Timothy A. Bell for breeding the diallel and CC-RIX mice. We also thank Alan B. Lenarcic for assistance with statistical analysis software, and Sarah D. Turner for assistance with data visualization. Many thanks to the referees and editors who contributed to this manuscript by providing the authors with detailed questions and comments.

Author contributions: PLM, MTF, GRK, MTH, and WV wrote the manuscript. PLM, MTF, DWT, FPMV, MTH, RSB, and WV designed experiments. PLM, MTF, DRM, ACW, AW, CRM, CRM, KEN, KSP, ASC, FPMV performed experiments. GDS bred the mice. PLM, GRK, and WV performed the statistical analysis. The authors declare no conflicts of interest.

## 10. APPENDIX A: DEFINING A POTENTIAL OUTCOMES MODEL OF TREATMENT RESPONSE WITH QUARTETS

In the potential outcomes framework of Neyman (1923) and Rubin (1974) the causal effect of an applied treatment on a measured outcome in an individual *i* is defined as the difference between the outcome under treatment and the outcome that would have been observed if *i* were instead to have received the control. In our case, for some outcome measure *y*, we defined the causal effect as the infection response

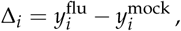

where 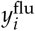 and 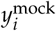 are “potential outcomes”, one of which is observed (the factual) and other of which is unobserved (the counterfactual). Since it is impossible to observe both simultaneously, the causal effect Δ_*i*_ can never be measured directly (Holland 1986). It can however be estimated as

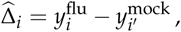

with the accuracy of this estimate depending on how closely *i′* matches *i*. Our desire for lack of bias in this measure motivates our treatment assignment being randomly ascribed within a group of matched individuals.

In the treatment-response diallel, we are primarily interested not in infection response for a particular mouse but rather the expectation of this quantity for mice within a given diallel category, or more generally within a group of matched individuals *q*,

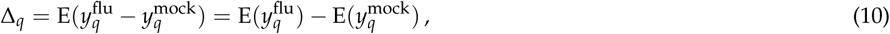

where in our case *q* is defined as mice specific to a given diallel category and experimental batch. In practice it is natural to estimate this quantity as

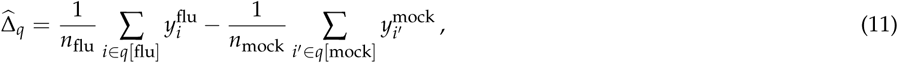

where *q*[flu] and *q*[mock] are, respectively, the set of mice in group *q* assigned to flu and mock treatment groups. The variance of this estimate is

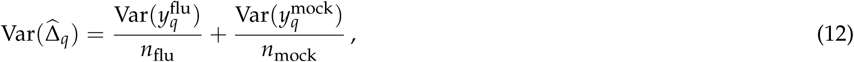

and if it is considered likely, as in this study, that the infected phenotypes will be more variable than the mock, Var(*y*^flu^) *>* Var(*y*^mock^), then it is most efficient experimentally to devote more individuals to the the infected arm than the mock arm, *i.e.*, *n*_flu_ *> n*_mock_.

In our experimental design, we have flu:mock in the ratio 3:1 for each group *q*. It is therefore natural to define the unit of observation *q* as a quartet, that is, Eq 11 with *n*_flu_ = 3 and *n*_mock_ = 1. This means that each diallel category can be represented by multiple quartets, corresponding to multiple observations of 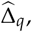, denoted in the **Methods** as delta_*q*_.

We now note several assumptions and connections. Eq 10 equates unit-level with marginal causal effects and thereby assumes no interference between units, specifically, that mice in the same quartet do not affect each other’s outcomes; this is approximately true based on the well-established evidence that mice do not transmit H1N1 influenza virus (Lowen *et al.* 2006; Edenborough *et al.* 2012), a finding we have also verified by weight loss profiles and RNA-seq of CC founder strains co-housed with H1N1(PR8)-infected mice (Xiong *et al.* 2014). Last, the definition of 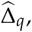 in Eq 11 is analogous to an inverse probability weighted causal effect estimate.

## 11. APPENDIX B: STOCHASTIC REGRESSION IMPUTATION OF MISSING QUARTET PARTNERS

The equation for the variance of 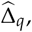, namely Eq 12, implies that modeling the residual in Eq 5 as homoskedastic would require *n*_flu_ and *n*_mock_ to be constant throughout — in other words, in order to ensure comparable precision of infection responses, all quartets should be complete. However, in the diallel experiment some combinations of batch and diallel category had one or more flu mice missing. In these cases, quartets were defined to have missing values that would be filled in by imputation. The imputation scheme used here corresponds to stochastic regression imputation (*e.g.*, Gelman and Hill 2006) whereby the incomplete dataset is repeatedly augmented to a completed data set using sampled variates from a prediction model, each completed dataset is subject to the BayesDiallel analysis described in **Methods**, and then results across the completed datasets are aggregated.

Each imputation required two steps: since the target phenotype of a missing mouse, namely its p.i. weight loss, was considered potentially dependent on its day 0 weight, we first imputed missing values for D0 and then imputed missing p.i. weight loss conditional on D0. In addition, at each day p.i., there was one batch/diallel category combination with 1 mock and 4 infecteds; for this case only, in each round of imputation, we created a completed quartet by randomly deleting one of the four infecteds.

The two-step stochastic regression imputation was performed as follows. Define the observed diallel data for D0 and pct^day[flu]^ as D0_obs_ and 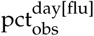 respectively, and lett 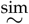 represent regression with BayesDiallel followed by stochastic regression imputation, that is, sampling from the posterior predictive. For each *t* = 1, …, 1000 round of imputation, we first impute missing D0 values as

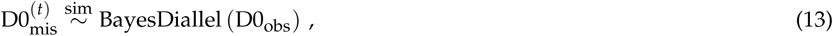

where BayesDiallel is fitted as Model 1 in Table 2. These imputed values are then combined with observed D0 values to give the completed set,

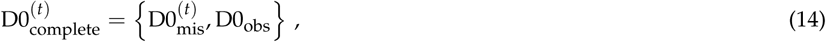

which are then used to impute p.i. weight loss at all timepoints (D1 to D4) for the missing mice as

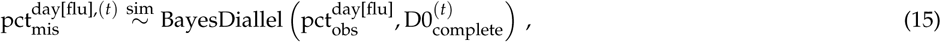

leading in each case to the completed flu p.i. weight loss data,

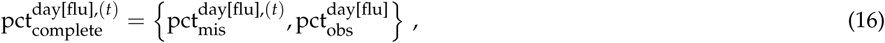

and subsequent calculation of quartet-based infection response values as

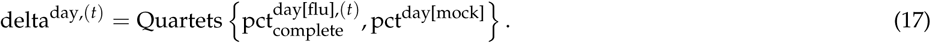

Each of the 1000 infection response data sets is analyzed separately using BayesDiallel (Model 3 in Table 2). The MCMC chains from each replicate are aggregated and thinned (sampled across even intervals) and the aggregate results are reported, according to the procedure outlined in Algorithm 1 in the Supplement.

The number of animals imputed were 33 at D1, 33 at D2, 15 at D3, and 16 at D4, in each case corresponding to a small proportion (2.8%-3.1%) of the total data set. Phenotypes were not impputed non-productive diallel genotypes (which contain no mock or infected mice at all), *i.e. jk* ∈ {NZO×CAST, NZO×PWK} (Chesler *et al.* 2008).

## 12. DATA UPLOAD DETAILS

Data and analysis files in this supplement include:

- File_S1_Readme.pdf - read me file providing overview of data upload.
- File_S2_Packages.tar.gz - packages used for analysis.
- File_S3_FluDiData.csv - diallel data file.
- File_S4_diallel.tar.gz - diallel analysis files.
- File_S5_preCC.tar.gz - pre-CC data and analysis files.
- File_S6_CC-RIX.tar.gz - CC-RIX data and analysis files.

After unzipping supplemental files S2, S4, S5, and S6 in a common folder, the directory structure will be as follows:

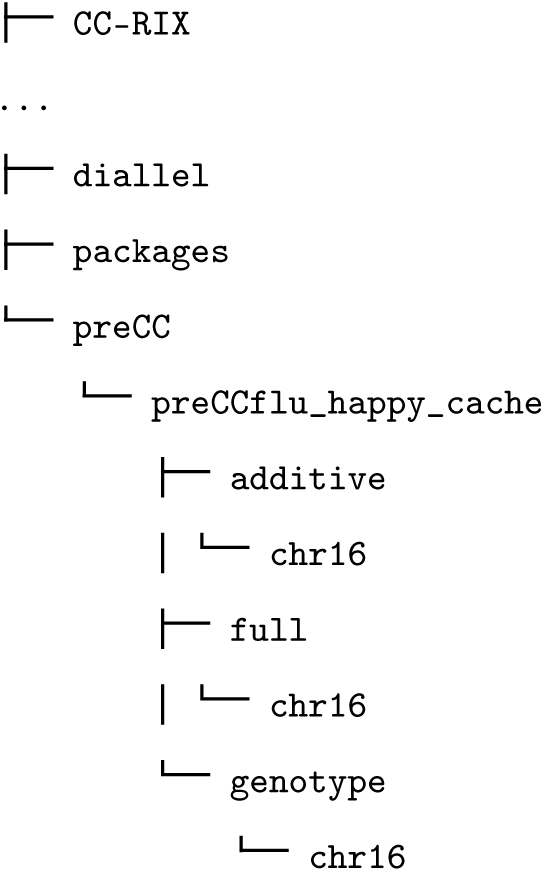

### A. Diallel Analysis

The files that are needed to reproduce the diallel analysis are given in the “diallel” folder, which includes the raw phenotype file, `FluDiData.csv,` and the following settings files, `CategoriesToImputeD4.csv, and settings.config.D4.` Additional *.R, *.sh, and *.pl scripts necessary to run the analysis are included.

To run the analysis for D4 treatment response on a macOS machine with both R and Perl installed, first install prerequisites, as necessary, and then install BayesDiallel, BayesSpike, and treatmentResponseDiallel. On a bash shell, from the “diallel” directory, use the following command (example using D4):

~~~
bash MIMQ_Run_D4.sh &
~~~

When the jobs on all cores are complete, then run:

~~~
bash MIMQ_PostRun_D4.sh &
~~~

Again, when the jobs on all cores are complete, run the following to clean up temporary files:

~~~
bash MIMQ_cleanup.sh &
~~~

### B. pre-CC Analysis

The files that are needed to reproduce the pre-CC analysis are given in the “preCC” folder. The raw phenotype file is `Flu-preCC-data.csv,` and the analysis script is `main_analysis_preCC.R`. To run the analysis, install the required prerequisites, and then INLA and Diploffect; proceed to run the analysis script from R.

### C. CC-RIX Analysis

The files that are needed to reproduce the CC-RIX analysis are given in the “CC-RIX” folder. The raw phenotype file is `Flu-CC-RIX-data.csv,` and the analysis script is `main_analysis_CC_RIX.R.` To run the analysis, install the required prerequisites, and then INLA and Diploffect; proceed to run the analysis script from R.

### D. Packages

The specific versions of the main analysis software packages used in this paper are provided in the “packages” folder. The packages, data and code required to reproduce the analysis in this manuscript are available as a data repository at http://dx.doi.org/10.5281/zenodo.293015, which is a static version of the GitHub repository maintained here: https://github.com/mauriziopaul/flu-diallel. The BayesDiallel and Diploffect software packages have been previously published (Lenarcic et al., 2012, *Genetics*; Zhang et al., 2014, *Genetics*), and are available at the Valdar lab website software page (http://valdarlab.unc.edu/software.html). The R package that facilitates the treatment response analysis in this paper is provided in the file `treatmentResponseDiallel_0.0.0.9000.tar.gz,` and is available at the following GitHub site: https://github.com/mauriziopaul/treatmentResponseDiallel. It can be installed according to instructions on the website. It can also be installed using the command ‘R CMD INSTALL treatmentResponseDiallel_0.0.0.9000.tar.gz’ in a terminal. The following prerequisite packages for treatmentResponseDiallel can be installed in a similar fashion: `configfile_0.11.tar.gz` and `cmdline_1.1.tar.gz`.

### E. File Types

- *.R - These are R scripts used for analysis of the respective populations.
- *.pl - This is a Perl script used in the diallel analysis.
- *.sh - These are bash scripts used for the diallel analysis.
- *.RData - These files in the happy_format and happy_cache directories contain data that can be loaded in R that is necessary for the Diploffect models to run.
- *.csv - These are comma-separated files containing raw data and/or analysis settings.
- *.alleles - This file can be used to generate a full model in the happy cache.

### SUPPLEMENTAL FIGURES AND TABLES

**Figure S1.**
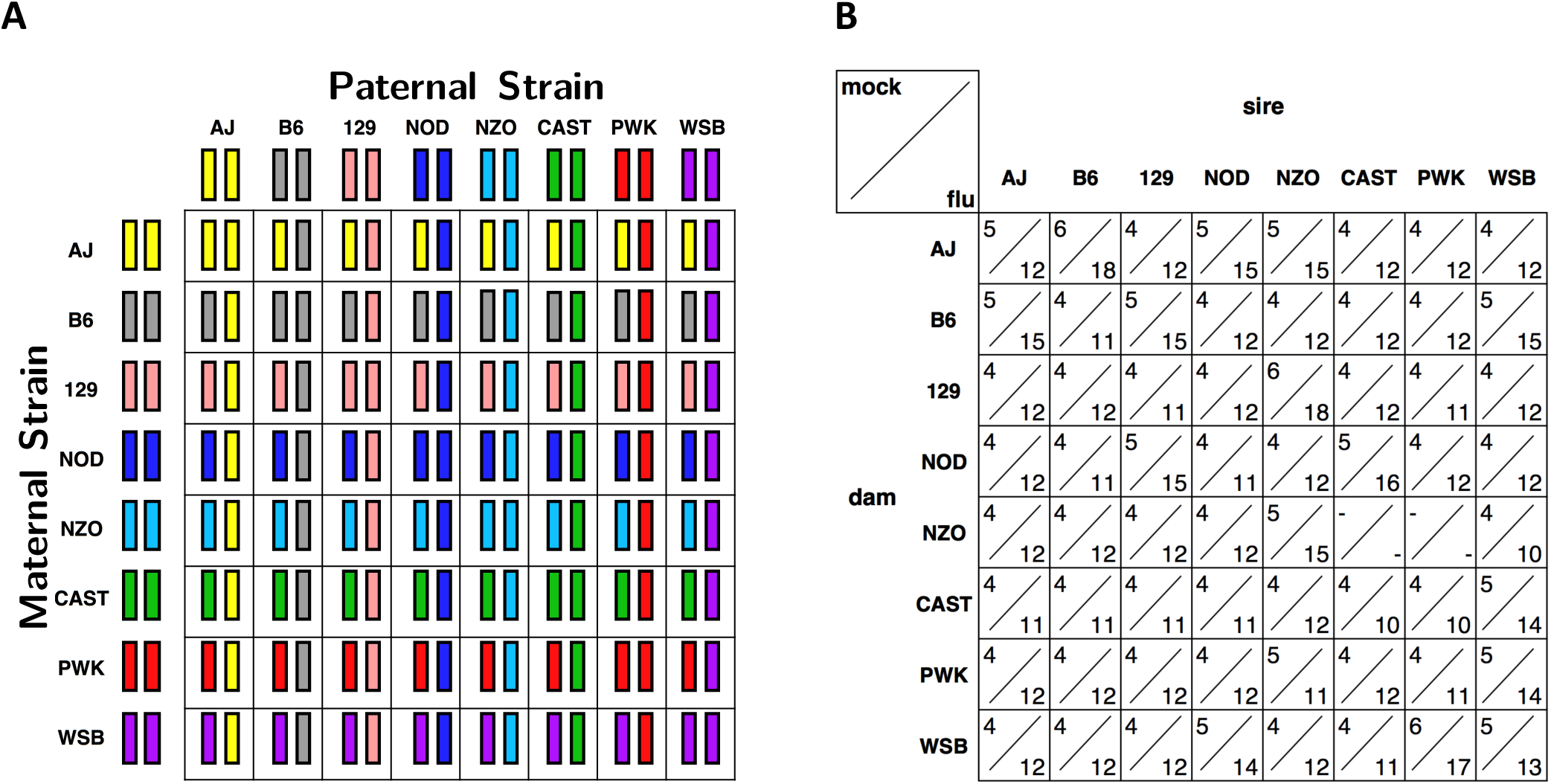
Diallel crossing design for the influenza diallel experiment. (A) Eight inbred *Mus musculus* strains were crossed in both directions to generate animals used in this experiment. All of the animals were followed from D0 through D2. Approximately half of the animals were followed through D4, and the remaining half of the mice were sacrificed on D2 (for analysis not included in this manuscript). (B) Of the 1,043 mice in this experiment, 268 received mock treatment, and 775 received influenza (flu) inoculation, or approximately 1 mock for every 2.9 infected mice within each category. There were one hundred twenty-nine inbred mice and 914 heterozygous (hybrid F1) mice used in this study. Crosses that were not observed (marked with a "-") include NZO×CAST and NZO×PWK, for which viable offspring cannot be produced.

**Figure S2.**
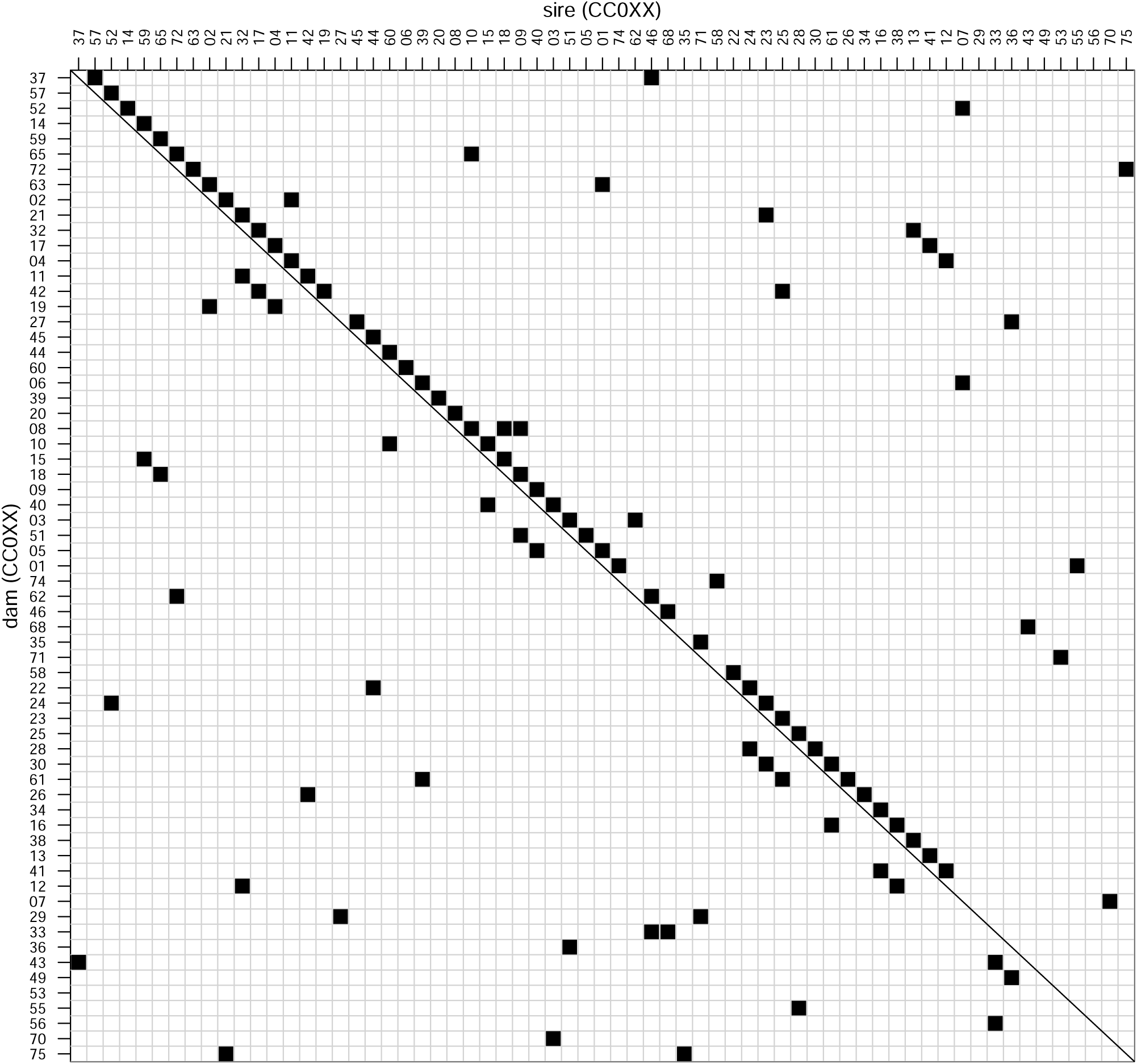
CC-RIX crossing design for the influenza infection experiment. As part of a larger CC-RIX QTL mapping experiment, 65 Collaborative Cross (CC) RI strains were crossed, approximately once as dam and once as sire in a round robin breeding scheme, to generate female animals in 105 CC-RIX lines, aged 8-12 weeks, that were sacrificed on day 7 (D7) post-infection, and used in this experiment. Weights for all of the animals were measured from D0 through D7. There were 1,402 heterozygous (hybrid F1) mice used in this study.

**Figure S3.**
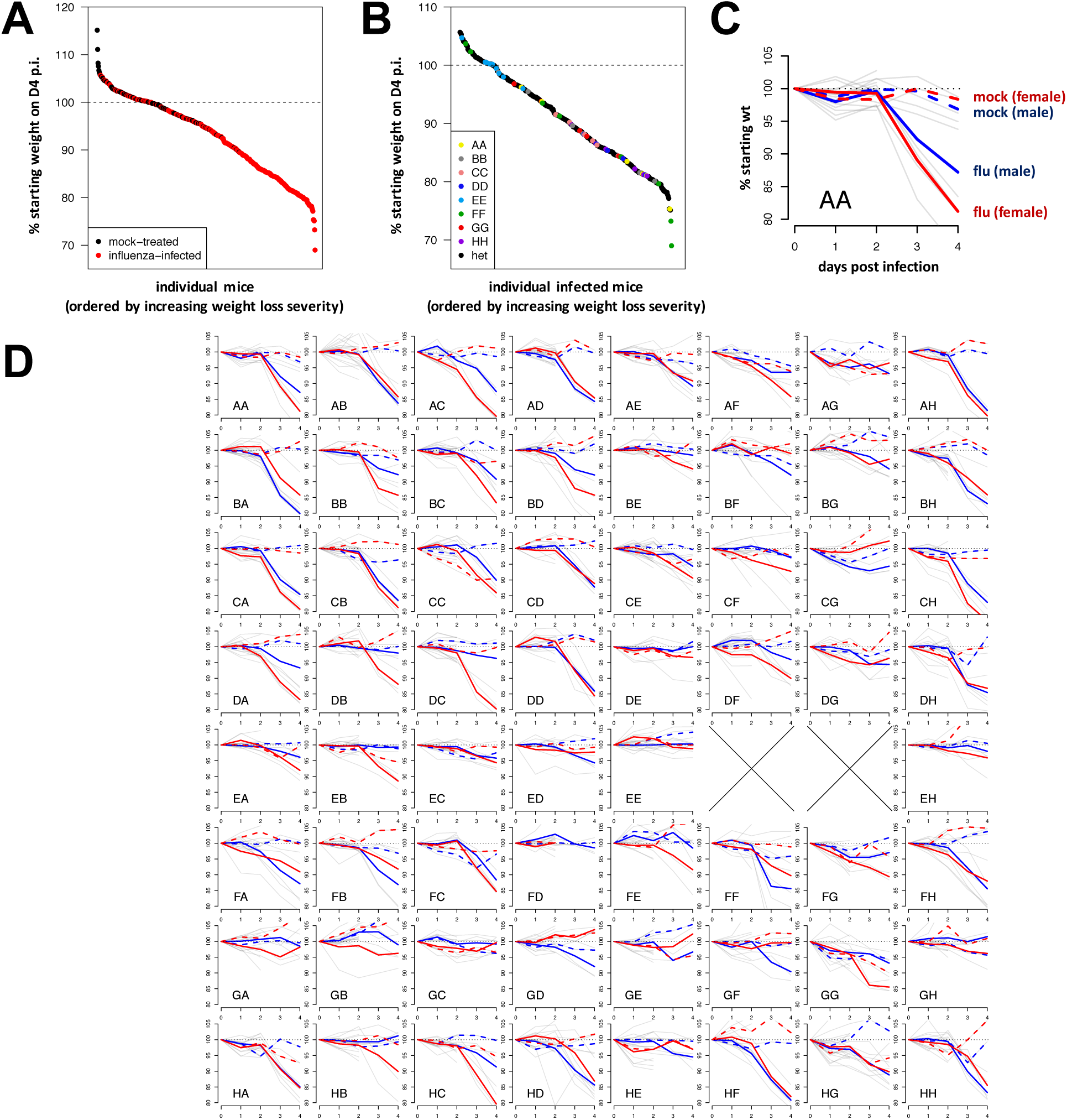
The raw distribution and timecourse of mock- and flu-induced weight change in the diallel. (A) Rank-ordered distribution of individual mock- and flu-induced percent weight change of inbred and F1 animals at D4 p.i. (B) Rank-ordered distribution of inbred and F1 flu-infected percent weight change at D4 p.i. (C) An example of raw weight change curves for AA (AJ×AJ) male and female mice. (D) The panel of raw weights for the 62 possible diallel categories. In (C) and (D), y-axes indicate percent of D0 weight, and x-axes indicate day(s) post-infection, while grey lines indicate individual weight change trajectories; colored lines indicate mean male (blue) and mean female (red) weight trajectories, and among these means, colored line styles indicate mock (dashed) and influenza (solid) treatment means within each group. In (D), the large X’s in EF and EG indicate non-productive F1 matings. Inbred mouse lines AJ, B6, 129, NOD, NZO, CAST, PWK, and WSB are indicated by letters A through H, respectively, with the first letter indicating maternal parentage and the second letter indicating paternal parentage.

**Figure S4.**
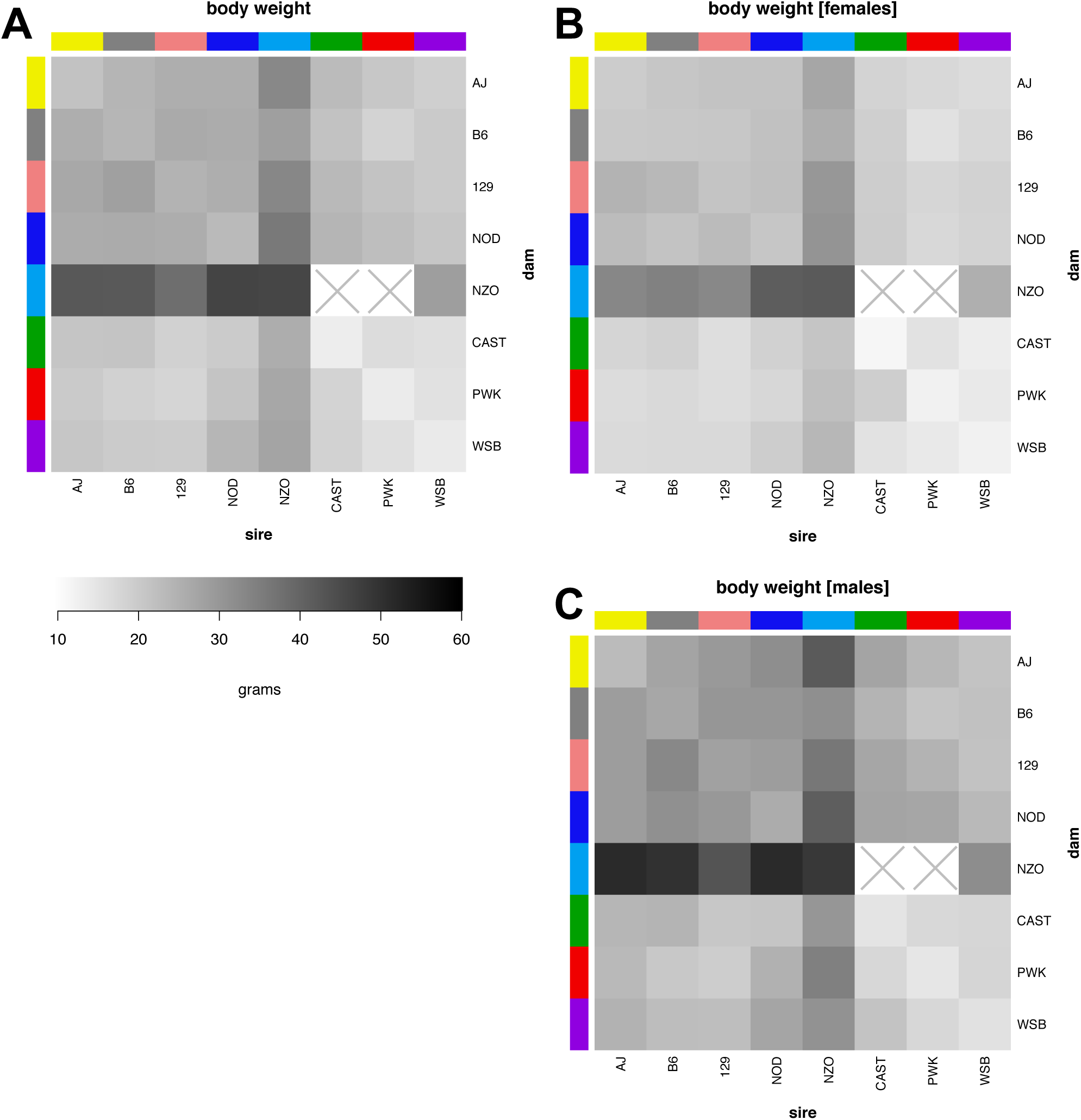
Body mass in a diallel cross of inbred mice. (A) Baseline body weight (in grams) for (B) male and (C) female 8-12 week old mice (n=1,043), presented as mean values across 62 inbred and F1 diallel categories.

**Figure S5.**
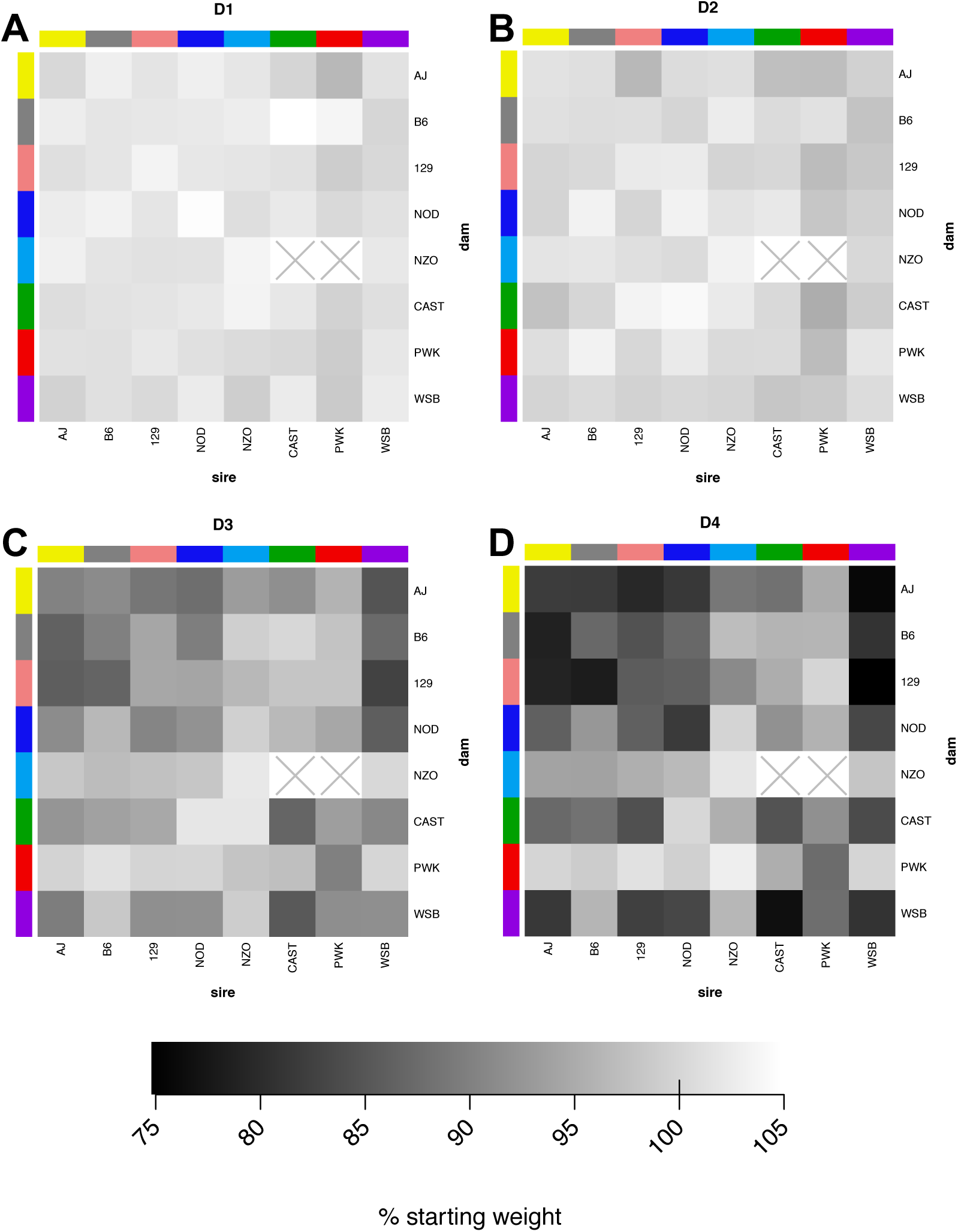
Weight loss in influenza-infected mice from a diallel cross of inbred mice. (A) Change in body weight on (A) day 1 (D1), (B) D2, (C) D3, and (D) D4 post-infection, presented as mean values of percent starting weight across 62 inbred and F1 diallel categories.

**Figure S6.**
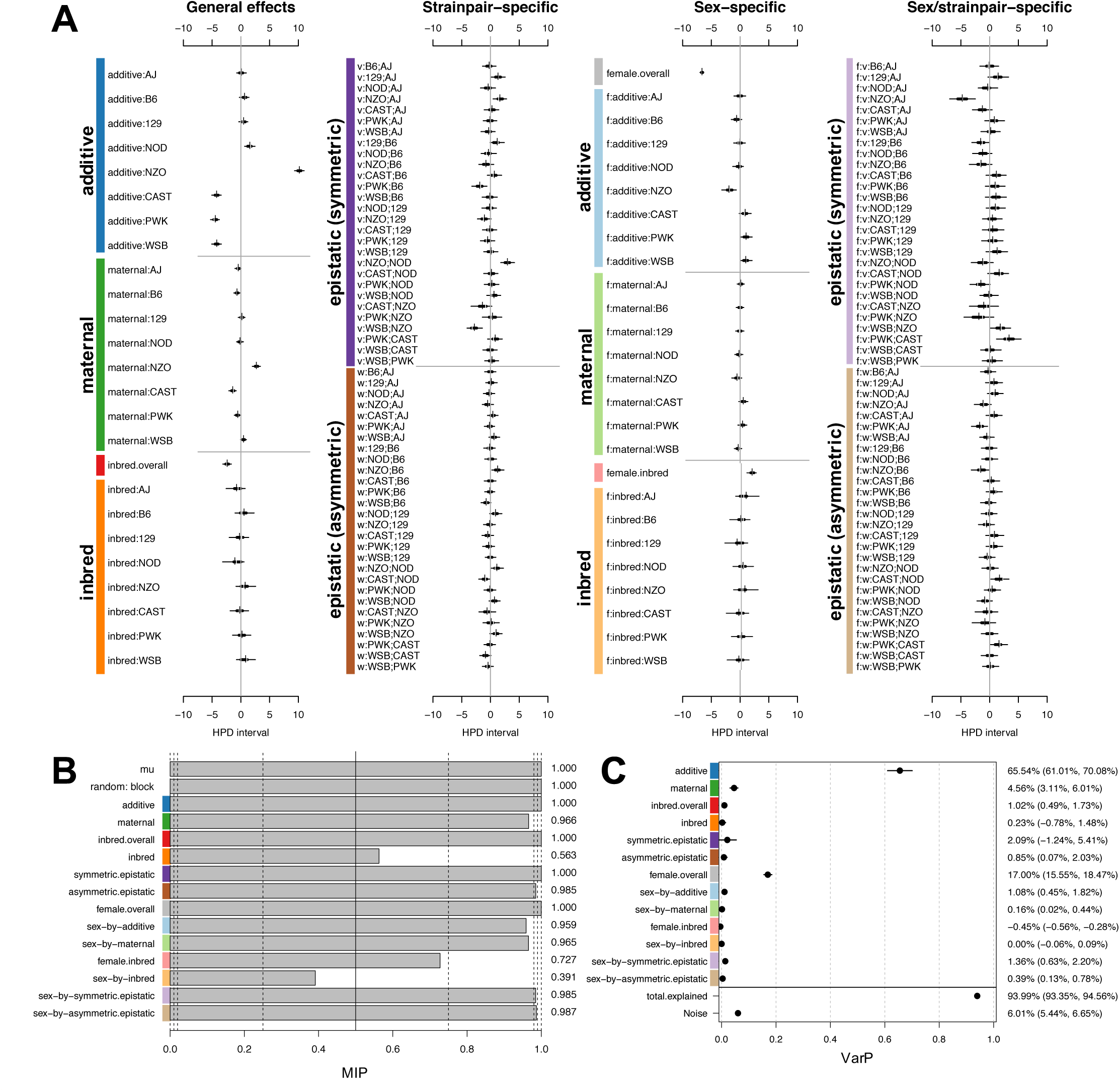
Diallel effects on D0 body weight in a diallel cross of inbred mice. (A) Effect estimates for additive, maternal, inbred, and epistatic effects, including sex-specific effects, are presented as highest posterior density intervals (in grams) for adult baseline body weight in 8-12 week old mice (n=1,043). Parameters are labeled according to the methods, and intervals are presented as in Figure 4A. Symmetric epistatic, asymmetric epistatic, and sex-specific parameters are indicated by “v:”, “w:”, and “f:”, respectively. The overall mean, *µ*, (not shown) is 23.97 (23.56, 24.41) grams. (B) Posterior mean model inclusion probabilities (MIPs) are given for effect parameter classes. (C) Variance projection (VarPs), a generalization of heritability to the diallel effects classes, are shown for three overall effects, five random effects classes, and five corresponding sex-specific random effects classes (with posterior median and 95% HPDs).

**Figure S7.**
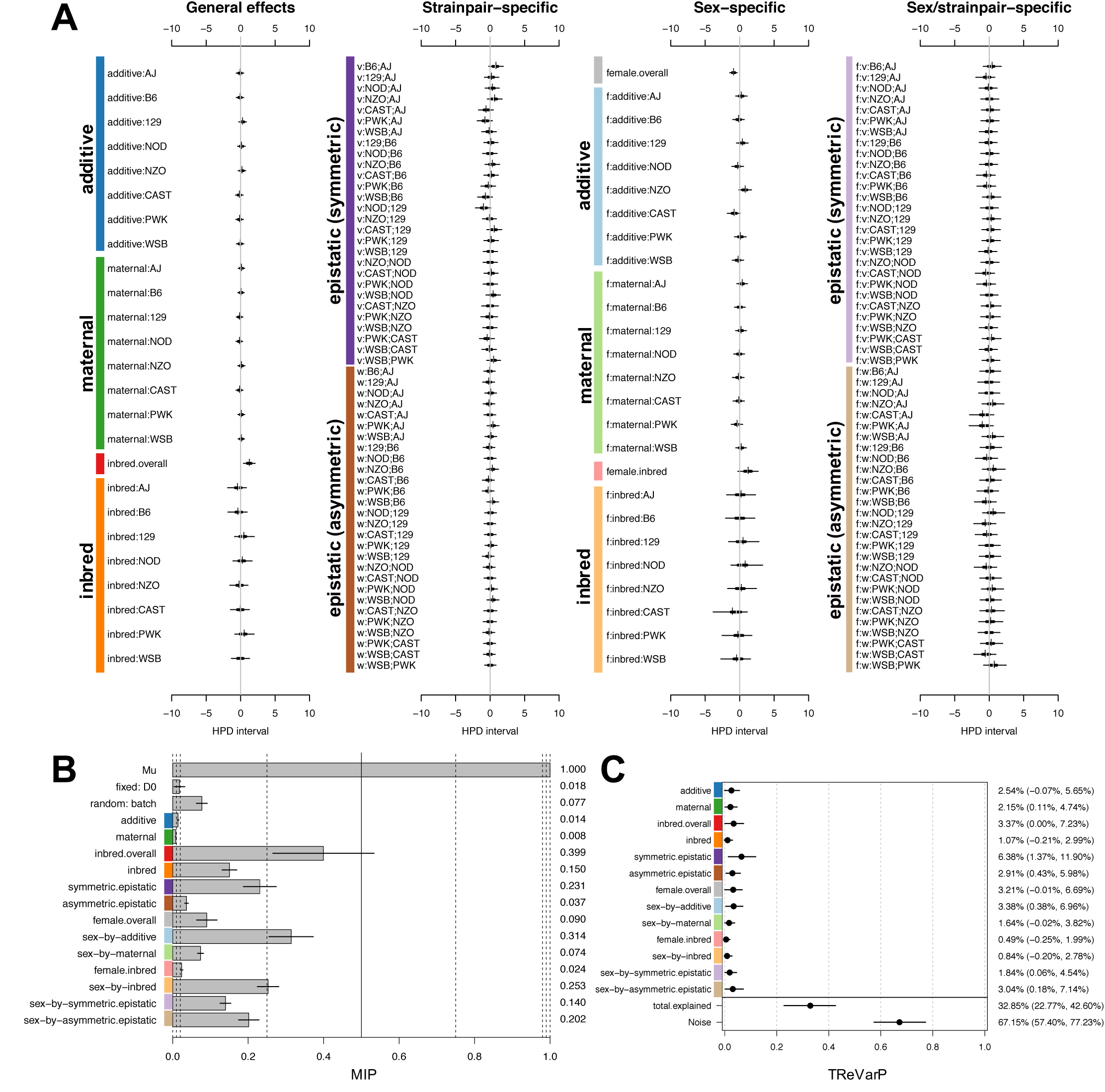
Diallel effects on host infection response (weight change) at D1 post-infection, using multiple imputation matched quartets. Effect estimates for additive, maternal, inbred, and epistatic effects, including sex-specific effects, are presented as highest posterior density intervals (in percent starting weight) in 8-12 week old mice (n=1,042). Parameters are labeled according to the methods, and intervals are presented as in Figure 4A. Symmetric epistatic, asymmetric epistatic, and sex-specific parameters are indicated by “v:”, “w:”, and “f:”, respectively. The overall treatment effect *θ* is −0.131% (−0.484%,0.223%). (B) Model inclusion probabilities (MIPs) are given (posterior mean ± 1 s.d.) for effect parameter classes. (C) Treatment response variance projections (TReVarPs), a generalization of heritability to the diallel effects classes, are shown for three overall effects, five random effects classes, and five corresponding sex-specific random effects classes (with posterior median and 95% HPDs).

**Figure S8.**
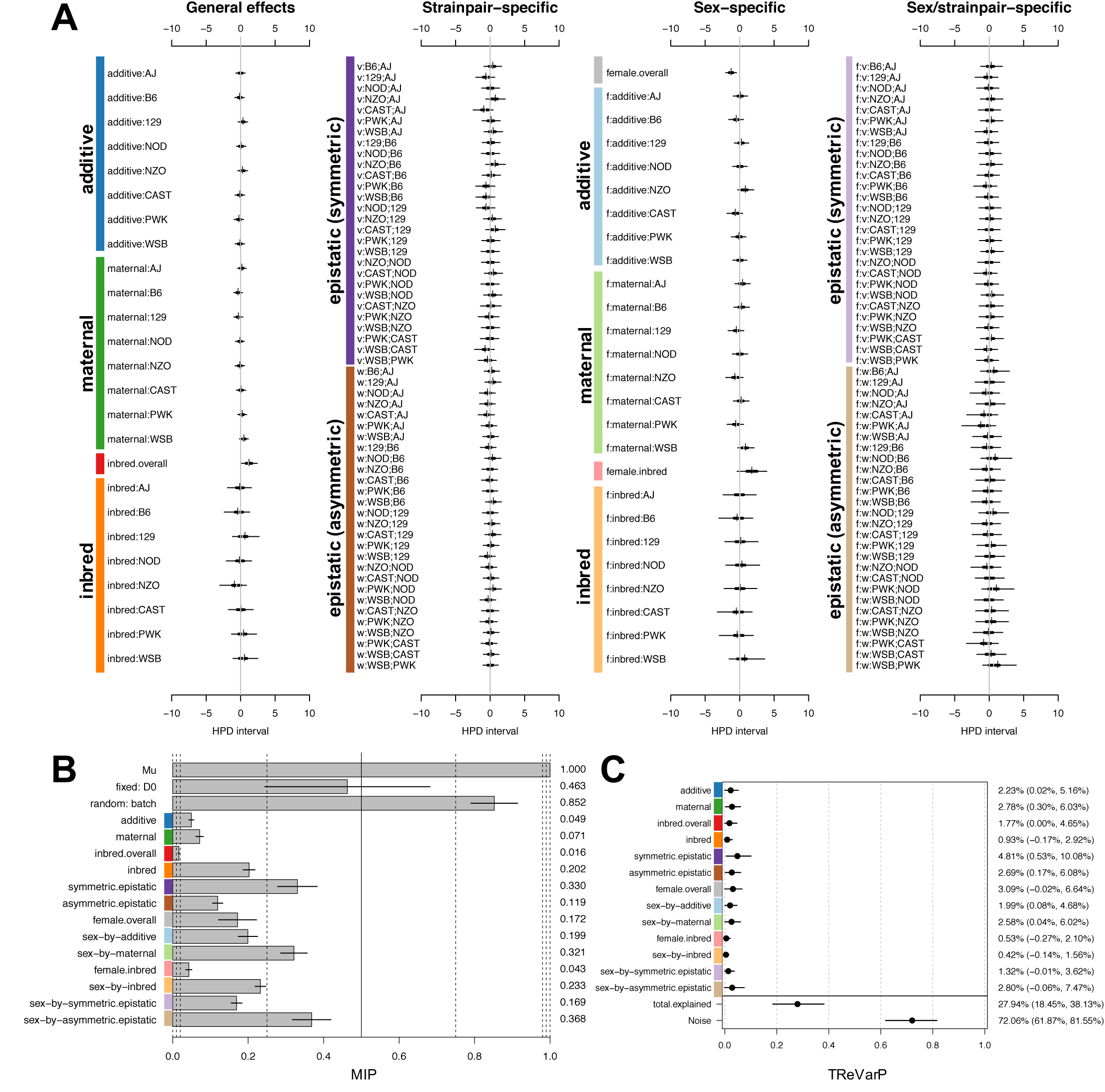
Diallel effects on host infection response (weight change) at D2 post-infection, using multiple imputation matched quartets. Effect estimates for additive, maternal, inbred, and epistatic effects, including sex-specific effects, are presented as highest posterior density intervals (in percent starting weight) in 8-12 week old mice (n=1,042). Parameters are labeled according to the methods, and intervals are presented as in Figure 4A. Symmetric epistatic, asymmetric epistatic, and sex-specific parameters are indicated by “v:”, “w:”, and “f:”, respectively. The overall treatment effect *θ* is −0.833% (−1.328%,−0.318%). (B) Model inclusion probabilities (MIPs) are given (posterior mean ± 1 s.d.) for effect parameter classes. (C) Treatment response variance projections (TReVarPs), a generalization of heritability to the diallel effects classes, are shown for three overall effects, five random effects classes, and five corresponding sex-specific random effects classes (with posterior median and 95% HPDs).

**Figure S9.**
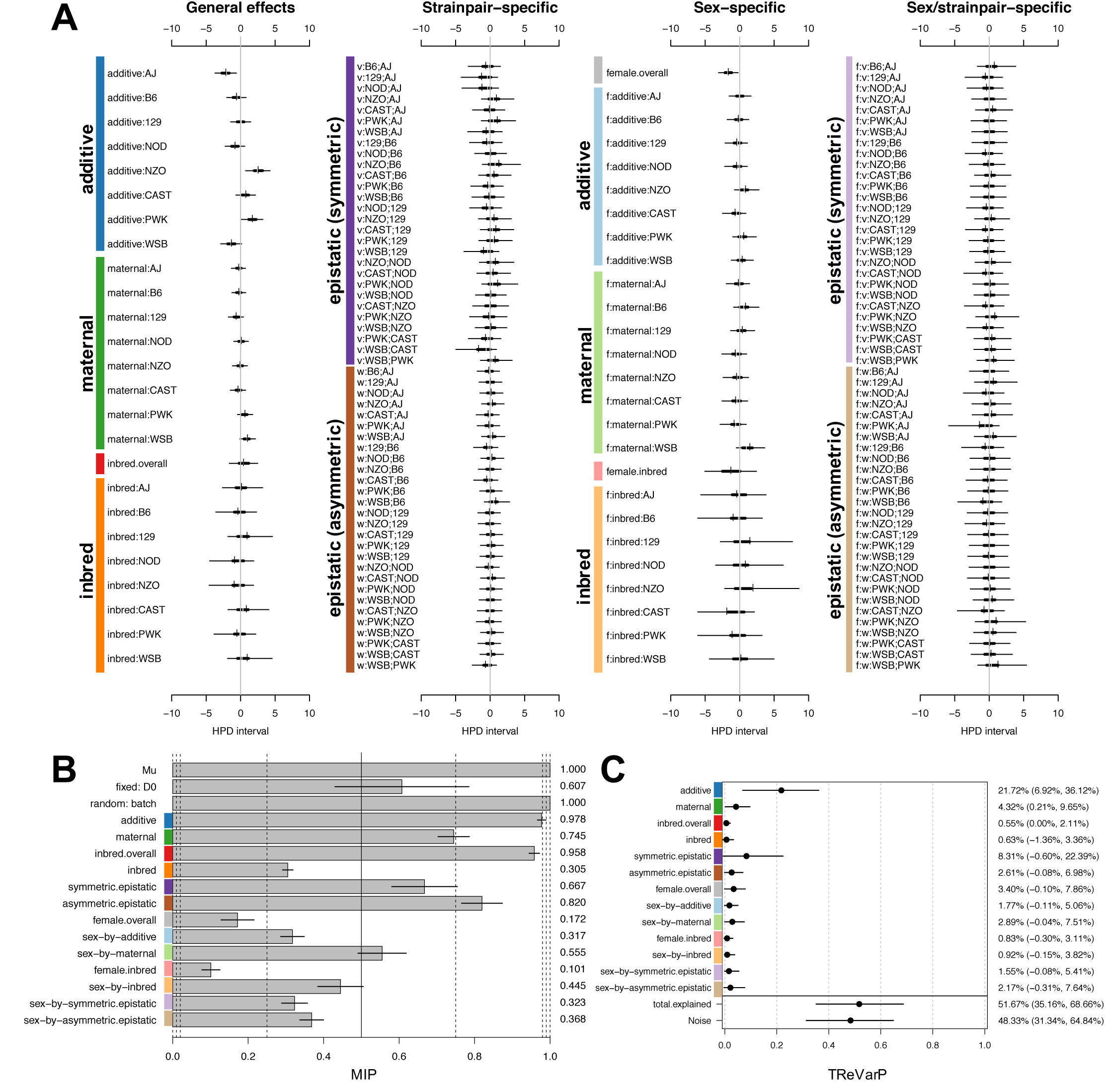
Diallel effects on host infection response (weight change) at D3 post-infection, using multiple imputation matched quartets. Effect estimates for additive, maternal, inbred, and epistatic effects, including sex-specific effects, are presented as highest posterior density intervals (in percent starting weight) in 8-12 week old mice (n=514). Parameters are labeled according to the methods, and intervals are presented as in Figure 4A. Symmetric epistatic, asymmetric epistatic, and sex-specific parameters are indicated by “v:”, “w:”, and “f:”, respectively. The overall treatment effect *θ* is −5.594% (−6.470%, −4.729%). (B) Model inclusion probabilities (MIPs) are given (posterior mean ± 1 s.d.) for effect parameter classes. (C) Treatment response variance projections (TReVarPs), a generalization of heritability to the diallel effects classes, are shown for three overall effects, five random effects classes, and five corresponding sex-specific random effects classes (with posterior median and 95% HPDs).

**Figure S10.**
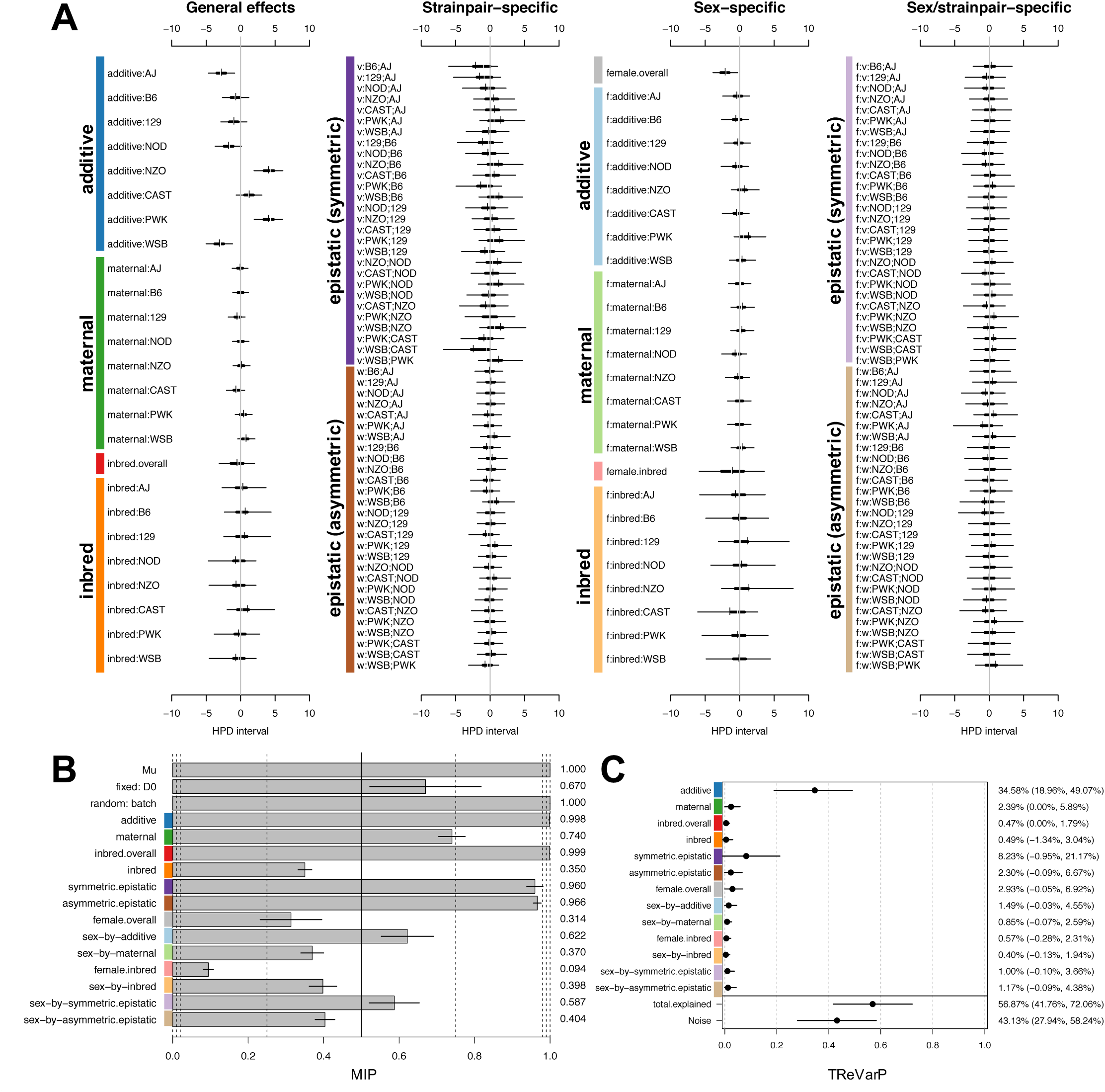
Diallel effects on host infection response (weight change) at D4 post-infection, using multiple imputation matched quartets. Effect estimates for additive, maternal, inbred, and epistatic effects, including sex-specific effects, are presented as highest posterior density intervals (in percent starting weight) in 8-12 week old mice (n=513). Parameters are labeled according to the methods, and intervals are presented as in Figure 4A. Symmetric epistatic, asymmetric epistatic, and sex-specific parameters are indicated by “v:”, “w:”, and “f:”, respectively. The overall treatment effect *θ* is −8.849% (−9.920%, −7.779%). (B) Model inclusion probabilities (MIPs) are given (posterior mean ± 1 s.d.) for effect parameter classes. (C) Treatment response variance projections (TReVarPs), a generalization of heritability to the diallel effects classes, are shown for three overall effects, five random effects classes, and five corresponding sex-specific random effects classes (with posterior median and 95% HPDs).

**Figure S11.**
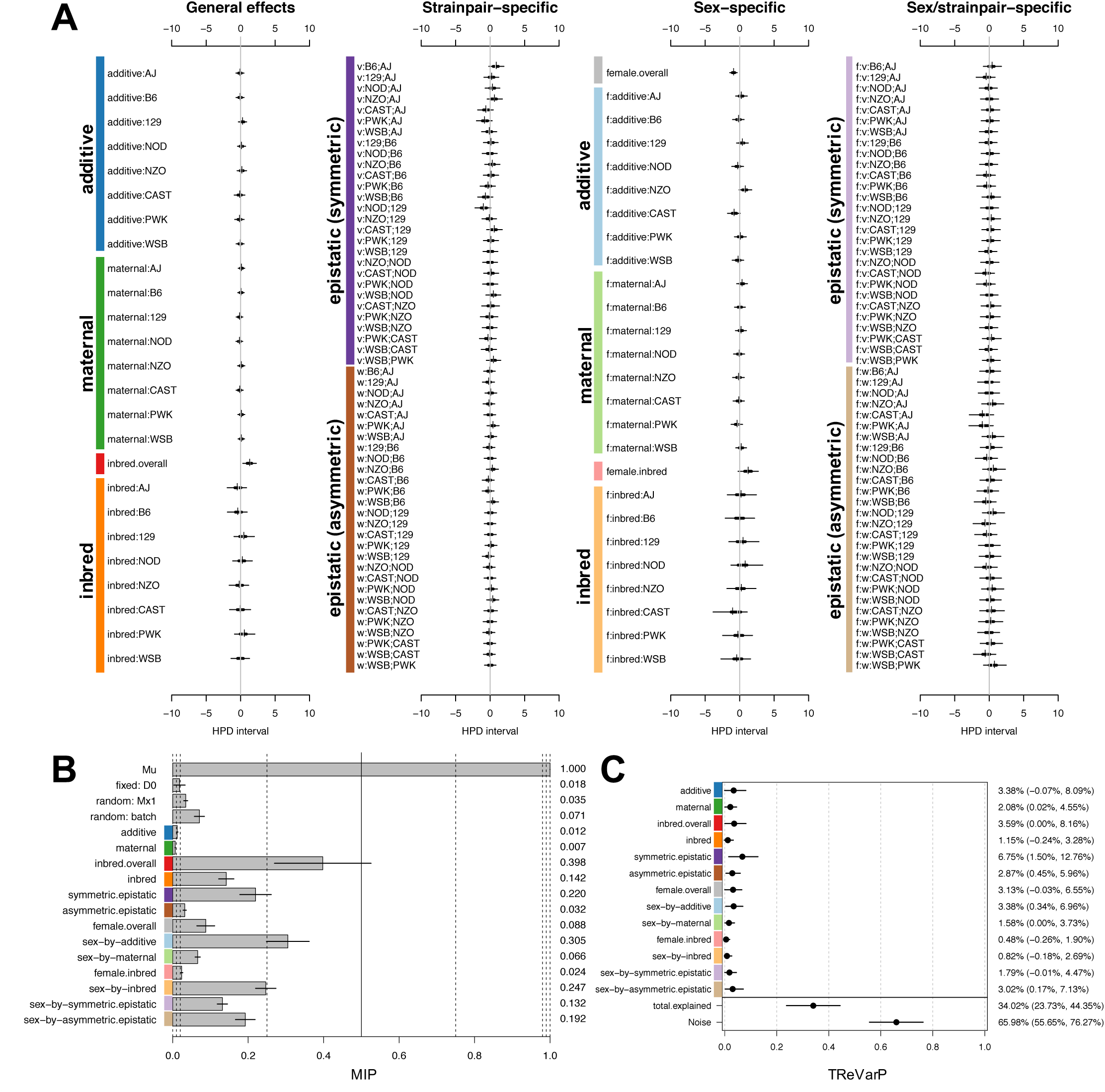
Diallel effects on host infection response (weight change) at D1 post-infection, using multiple imputation matched quartets and accounting for *Mx1*. Effect estimates for additive, maternal, inbred, and epistatic effects, including sex-specific effects, are presented as highest posterior density intervals (in percent starting weight) in 8-12 week old mice (n=1,042) after including random effect *u*(*Mx1* diplo). Parameters are labeled according to the methods, and intervals are presented as in Figure 4A. Symmetric epistatic, asymmetric epistatic, and sex-specific parameters are indicated by “v:”, “w:”, and “f:”, respectively. The overall treatment effect *q* is −0.193% (−1.128%, 0.707%). (B) Model inclusion probabilities (MIPs) are given (posterior mean ± 1 s.d.) for effect parameter classes. (C) Treatment response variance projections (TReVarPs), a generalization of heritability to the diallel effects classes, are shown for three overall effects, five random effects classes, and five corresponding sex-specific random effects classes (with posterior median and 95% HPDs).

**Figure S12.**
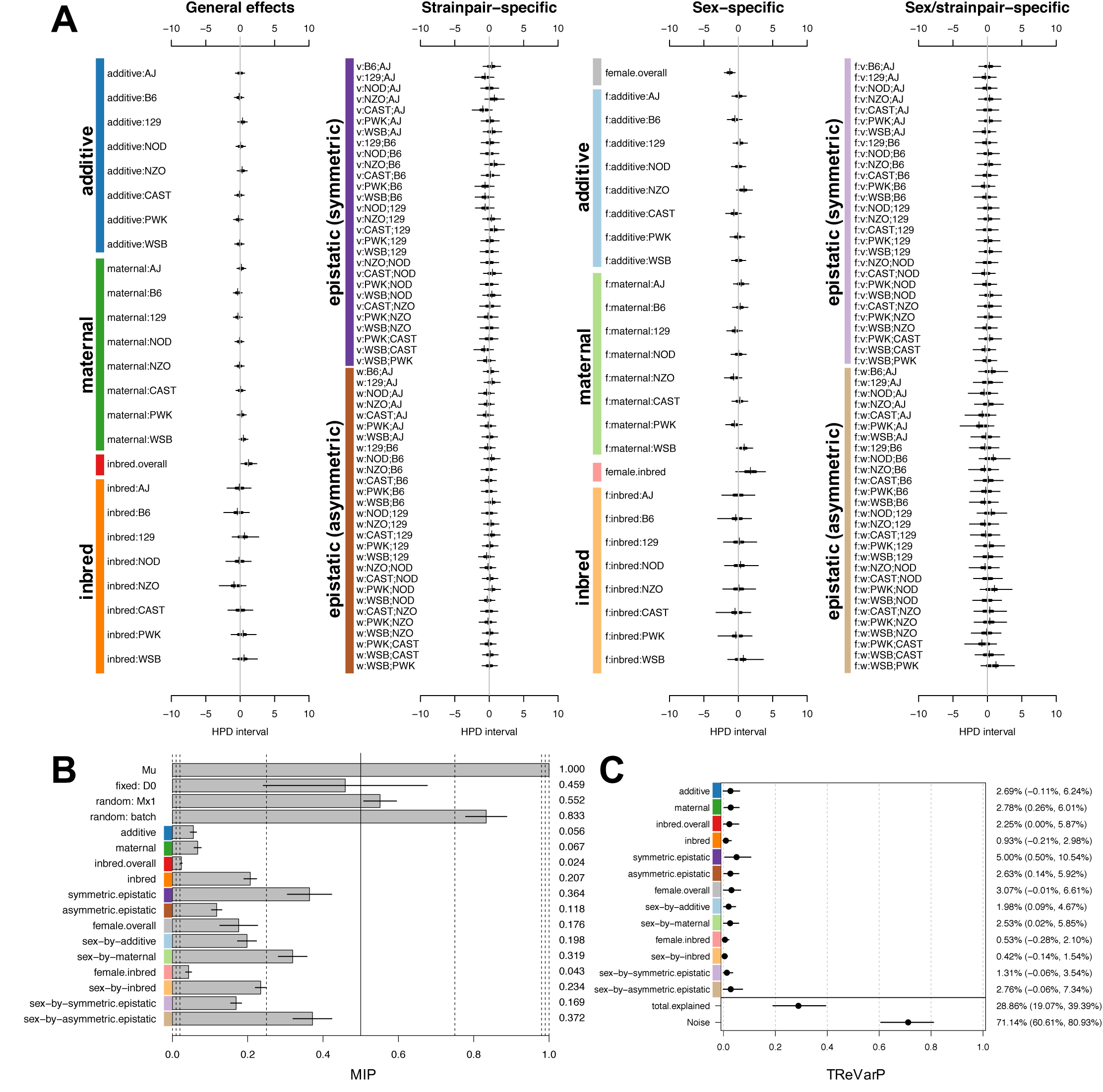
Diallel effects on host infection response (weight change) at D2 post-infection, using multiple imputation matched quartets and accounting for *Mx1*. Effect estimates for additive, maternal, inbred, and epistatic effects, including sex-specific effects, are presented as highest posterior density intervals (in percent starting weight) in 8-12 week old mice (n=1,042) after including random effect *u*^(*Mx1* diplo)^. Parameters are labeled according to the methods, and intervals are presented as in Figure 4A. Symmetric epistatic, asymmetric epistatic, and sex-specific parameters are indicated by “v:”, “w:”, and “f:”, respectively. The overall treatment effect *θ* is −0.925% (−1.991%, 0.113%). (B) Model inclusion probabilities (MIPs) are given (posterior mean ± 1 s.d.) for effect parameter classes. (C) Treatment response variance projections (TReVarPs), a generalization of heritability to the diallel effects classes, are shown for three overall effects, five random effects classes, and five corresponding sex-specific random effects classes (with posterior median and 95% HPDs).

**Figure S13.**
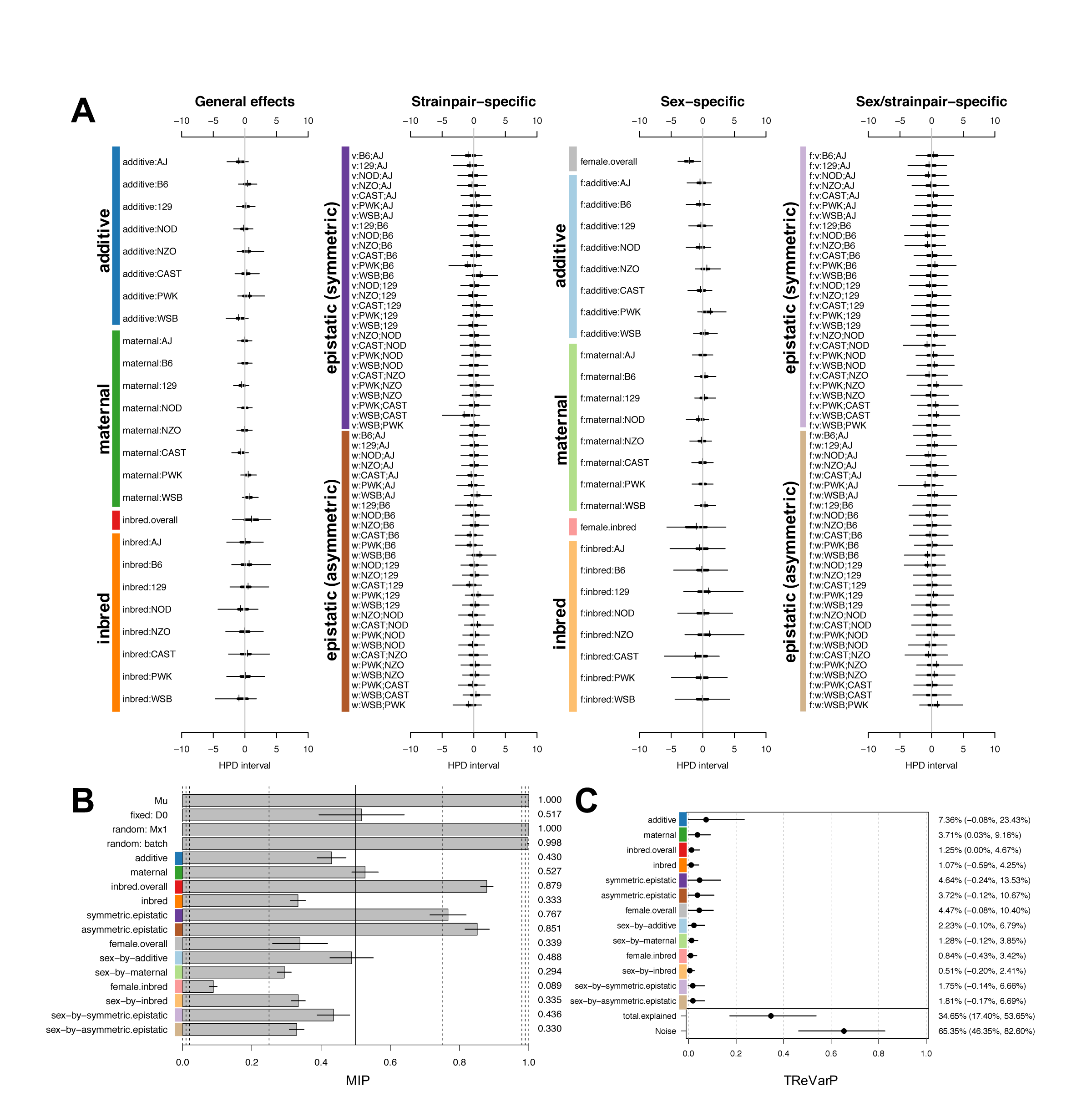
Diallel effects on host infection response (weight change) at D3 post-infection, using multiple imputation matched quartets and accounting for *Mx1*. Effect estimates for additive, maternal, inbred, and epistatic effects, including sex-specific effects, are presented as highest posterior density intervals (in percent starting weight) in 8-12 week old mice (n=514) after including random effect *u*^(*Mx1* diplo)^. Parameters are labeled according to the methods, and intervals are presented as in Figure 4A. Symmetric epistatic, asymmetric epistatic, and sex-specific parameters are indicated by “v:”, “w:”, and “f:”, respectively. The overall treatment effect *θ* is −5.429% (−7.675%, −3.102%). (B) Model inclusion probabilities (MIPs) are given (posterior mean ± 1 s.d.) for effect parameter classes. (C) Treatment response variance projections (TReVarPs), a generalization of heritability to the diallel effects classes, are shown for three overall effects, five random effects classes, and five corresponding sex-specific random effects classes (with posterior median and 95% HPDs).

**Figure S14.**
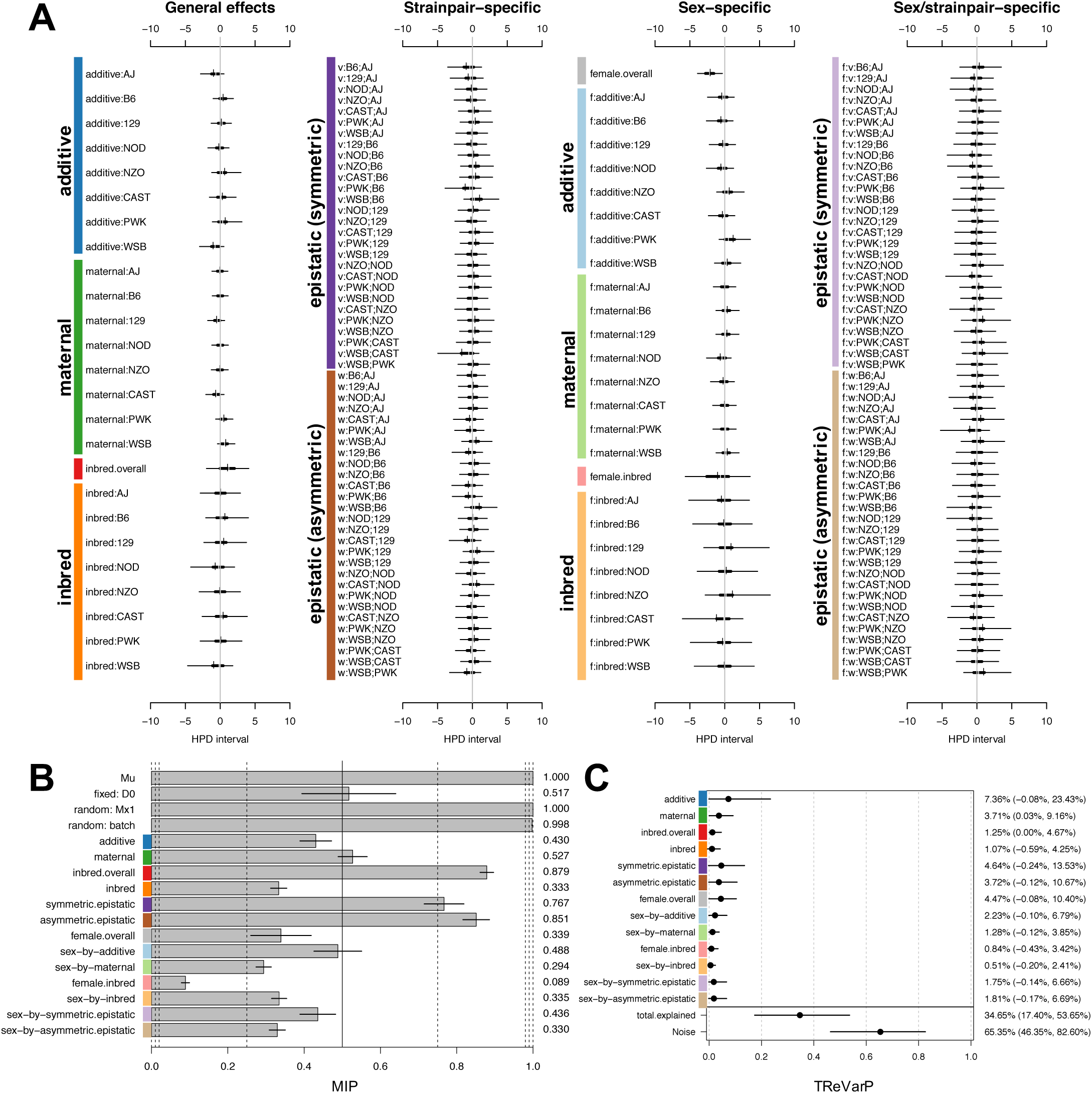
Diallel effects on host infection response (weight change) at D4 post-infection, using multiple imputation matched quartets and accounting for *Mx1*. Effect estimates for additive, maternal, inbred, and epistatic effects, including sex-specific effects, are presented as highest posterior density intervals (in percent starting weight) in 8-12 week old mice (n=513) after including random effect *u*^(*Mx1* diplo)^. Parameters are labeled according to the methods, and intervals are presented as in Figure 4A. Symmetric epistatic, asymmetric epistatic, and sex-specific parameters are indicated by “v:”, “w:”, and “f:”, respectively. The overall treatment effect *θ* is −7.986% (−11.202%, −4.147%). (B) Model inclusion probabilities (MIPs) are given (posterior mean ± 1 s.d.) for effect parameter classes. (C) Treatment response variance projections (TReVarPs), a generalization of heritability to the diallel effects classes, are shown for three overall effects, five random effects classes, and five corresponding sex-specific random effects classes (with posterior median and 95% HPDs).

**Table S1.** Proportion of variance in IAV-induced weight loss attributable to diallel effect classes across 4 days post-infection. Part C of Figure S7 through Figure S10 show Bayesian confidence interval plots of treatment response variance projection (TReVarP) values presented here.

| Effect Class <sup>a</sup> | D1 | 0.025 | 0.975 | D2 | 0.025 | 0.975 | D3 | 0.025 | 0.975 | D4 | 0.025 | 0.975 |
| --- | --- | --- | --- | --- | --- | --- | --- | --- | --- | --- | --- | --- |
| additive | 0.0229 | -0.0007 | 0.0565 | 0.0197 | 0.0002 | 0.0516 | 0.2163 | 0.0692 | 0.3612 | 0.3477 | 0.1896 | 0.4907 |
| maternal | 0.0192 | 0.0011 | 0.0474 | 0.0253 | 0.0030 | 0.0603 | 0.0381 | 0.0021 | 0.0965 | 0.0196 | 0.0000 | 0.0589 |
| inbred (overall) | 0.0306 | 0.0000 | 0.0723 | 0.0142 | 0.0000 | 0.0465 | 0.0025 | 0.0000 | 0.0211 | 0.0021 | 0.0000 | 0.0179 |
| inbred | 0.0082 | -0.0021 | 0.0299 | 0.0064 | -0.0017 | 0.0292 | 0.0031 | -0.0136 | 0.0336 | 0.0023 | -0.0134 | 0.0304 |
| symmetric epistatic | 0.0605 | 0.0137 | 0.1190 | 0.0433 | 0.0053 | 0.1008 | 0.0655 | -0.0060 | 0.2239 | 0.0692 | -0.0095 | 0.2117 |
| asymmetric epistatic | 0.0264 | 0.0043 | 0.0598 | 0.0233 | 0.0017 | 0.0608 | 0.0203 | -0.0008 | 0.0698 | 0.0163 | -0.0009 | 0.0667 |
| female (overall) | 0.0296 | -0.0001 | 0.0669 | 0.0282 | -0.0002 | 0.0664 | 0.0299 | -0.0010 | 0.0786 | 0.0256 | -0.0005 | 0.0692 |
| sex-by-additive | 0.0309 | 0.0038 | 0.0696 | 0.0169 | 0.0008 | 0.0468 | 0.0131 | -0.0011 | 0.0506 | 0.0103 | -0.0003 | 0.0455 |
| sex-by-maternal | 0.0143 | -0.0002 | 0.0382 | 0.0224 | 0.0004 | 0.0602 | 0.0230 | -0.0004 | 0.0751 | 0.0059 | -0.0007 | 0.0259 |
| female inbred (overall) | 0.0024 | -0.0025 | 0.0199 | 0.0027 | -0.0027 | 0.0210 | 0.0047 | -0.0030 | 0.0311 | 0.0027 | -0.0028 | 0.0231 |
| sex-by-inbred | 0.0054 | -0.0020 | 0.0278 | 0.0024 | -0.0014 | 0.0156 | 0.0034 | -0.0015 | 0.0382 | 0.0012 | -0.0013 | 0.0194 |
| sex-by-symm. epis. | 0.0152 | 0.0006 | 0.0454 | 0.0100 | -0.0001 | 0.0362 | 0.0089 | -0.0008 | 0.0541 | 0.0052 | -0.0010 | 0.0366 |
| sex-by-asymm. epis. | 0.0259 | 0.0018 | 0.0714 | 0.0218 | -0.0006 | 0.0747 | 0.0119 | -0.0031 | 0.0764 | 0.0059 | -0.0009 | 0.0438 |
| total explained | 0.3276 | 0.2277 | 0.4260 | 0.2775 | 0.1845 | 0.3813 | 0.5164 | 0.3516 | 0.6866 | 0.5705 | 0.4176 | 0.7206 |
| noise | 0.6724 | 0.5740 | 0.7723 | 0.7225 | 0.6187 | 0.8155 | 0.4836 | 0.3134 | 0.6484 | 0.4295 | 0.2794 | 0.5824 |

<sup>a</sup> To indicate the relative contribution of different diallel effects on host response to IAV, TreVarPs are given for overall effects (female, inbred, female inbred) and strain/strainpair-specific effects (additive, maternal, inbred, symmetric epistatic, asymmetric epistatic) on D1delta through D4delta (median and 95% HPD intervals).

**Table S2.**
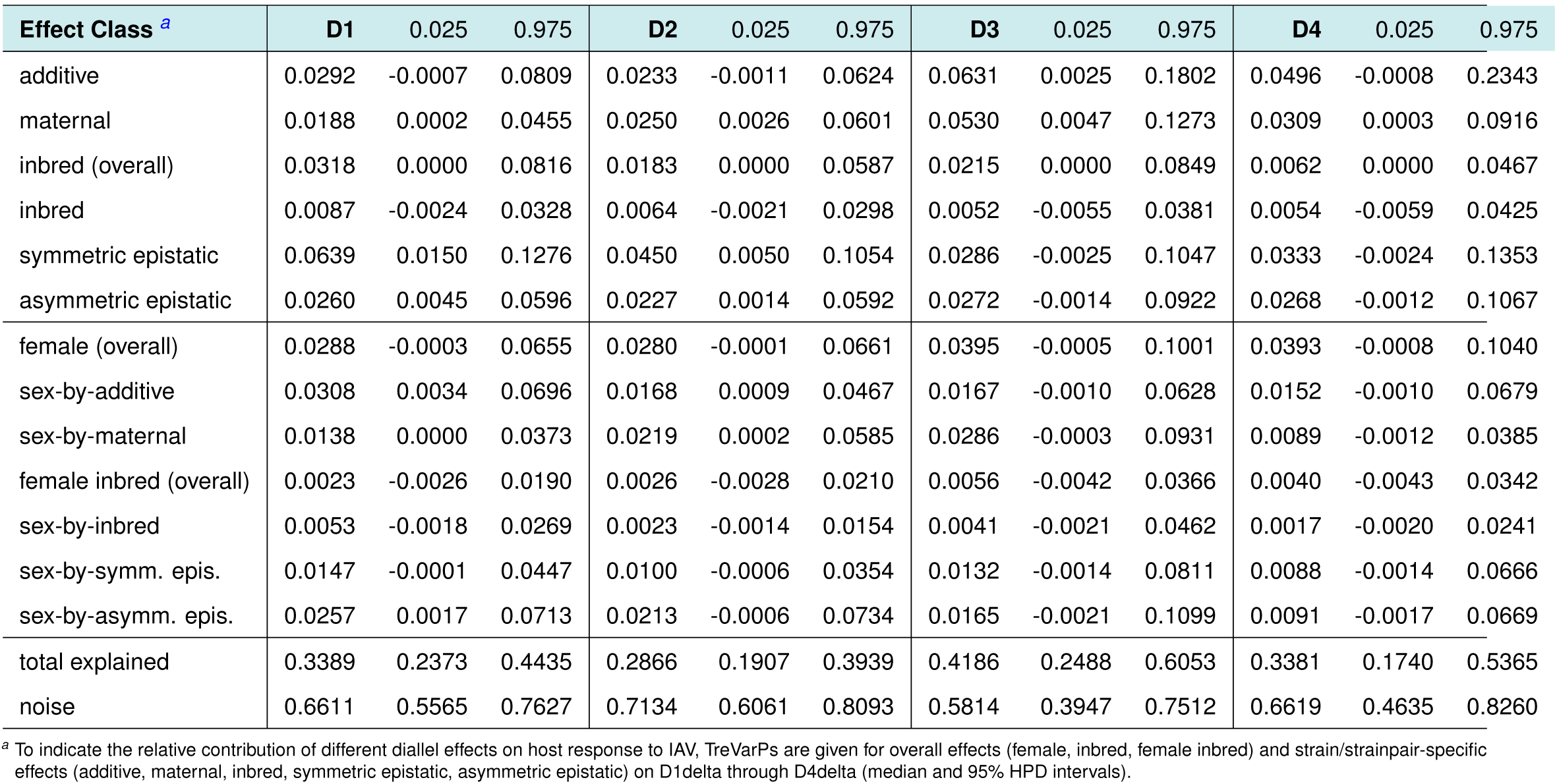
Proportion of variance in IAV-induced weight loss attributable to diallel effect classes across 4 days post-infection, after accounting for *Mx1* diplotype. Part C of Figure S11 through Figure S14 show Bayesian confidence interval plots of treatment response variance projection (TReVarP) values presented here.

| Effect Class <sup>a</sup> | D1 | 0.025 | 0.975 | D2 | 0.025 | 0.975 | D3 | 0.025 | 0.975 | D4 | 0.025 | 0.975 |
| --- | --- | --- | --- | --- | --- | --- | --- | --- | --- | --- | --- | --- |
| additive | 0.0292 | -0.0007 | 0.0809 | 0.0233 | -0.0011 | 0.0624 | 0.0631 | 0.0025 | 0.1802 | 0.0496 | -0.0008 | 0.2343 |
| maternal | 0.0188 | 0.0002 | 0.0455 | 0.0250 | 0.0026 | 0.0601 | 0.0530 | 0.0047 | 0.1273 | 0.0309 | 0.0003 | 0.0916 |
| inbred (overall) | 0.0318 | 0.0000 | 0.0816 | 0.0183 | 0.0000 | 0.0587 | 0.0215 | 0.0000 | 0.0849 | 0.0062 | 0.0000 | 0.0467 |
| inbred | 0.0087 | -0.0024 | 0.0328 | 0.0064 | -0.0021 | 0.0298 | 0.0052 | -0.0055 | 0.0381 | 0.0054 | -0.0059 | 0.0425 |
| symmetric epistatic | 0.0639 | 0.0150 | 0.1276 | 0.0450 | 0.0050 | 0.1054 | 0.0286 | -0.0025 | 0.1047 | 0.0333 | -0.0024 | 0.1353 |
| asymmetric epistatic | 0.0260 | 0.0045 | 0.0596 | 0.0227 | 0.0014 | 0.0592 | 0.0272 | -0.0014 | 0.0922 | 0.0268 | -0.0012 | 0.1067 |
| female (overall) | 0.0288 | -0.0003 | 0.0655 | 0.0280 | -0.0001 | 0.0661 | 0.0395 | -0.0005 | 0.1001 | 0.0393 | -0.0008 | 0.1040 |
| sex-by-additive | 0.0308 | 0.0034 | 0.0696 | 0.0168 | 0.0009 | 0.0467 | 0.0167 | -0.0010 | 0.0628 | 0.0152 | -0.0010 | 0.0679 |
| sex-by-maternal | 0.0138 | 0.0000 | 0.0373 | 0.0219 | 0.0002 | 0.0585 | 0.0286 | -0.0003 | 0.0931 | 0.0089 | -0.0012 | 0.0385 |
| female inbred (overall) | 0.0023 | -0.0026 | 0.0190 | 0.0026 | -0.0028 | 0.0210 | 0.0056 | -0.0042 | 0.0366 | 0.0040 | -0.0043 | 0.0342 |
| sex-by-inbred | 0.0053 | -0.0018 | 0.0269 | 0.0023 | -0.0014 | 0.0154 | 0.0041 | -0.0021 | 0.0462 | 0.0017 | -0.0020 | 0.0241 |
| sex-by-symm. epis. | 0.0147 | -0.0001 | 0.0447 | 0.0100 | -0.0006 | 0.0354 | 0.0132 | -0.0014 | 0.0811 | 0.0088 | -0.0014 | 0.0666 |
| sex-by-asymm. epis. | 0.0257 | 0.0017 | 0.0713 | 0.0213 | -0.0006 | 0.0734 | 0.0165 | -0.0021 | 0.1099 | 0.0091 | -0.0017 | 0.0669 |
| total explained | 0.3389 | 0.2373 | 0.4435 | 0.2866 | 0.1907 | 0.3939 | 0.4186 | 0.2488 | 0.6053 | 0.3381 | 0.1740 | 0.5365 |
| noise | 0.6611 | 0.5565 | 0.7627 | 0.7134 | 0.6061 | 0.8093 | 0.5814 | 0.3947 | 0.7512 | 0.6619 | 0.4635 | 0.8260 |

<sup>a</sup> To indicate the relative contribution of different diallel effects on host response to IAV, TreVarPs are given for overall effects (female, inbred, female inbred) and strain/strainpair-specific effects (additive, maternal, inbred, symmetric epistatic, asymmetric epistatic) on D1delta through D4delta (median and 95% HPD intervals).

**Table S3.** Model inclusion probabilities for diallel effects classes, (A) before and (B) after accounting for *Mx1* diplotype. We consider the following levels for strength of evidence for inclusion: positive {(0.05, 0.25] or [0.75, 0.95)}, strong {(0.01, 0.05] or [0.95, 0.99)} and very strong {[0, 0.01] or [0.99, 1]}, as described in the Statistical Methods. Part B of Figures S7 through S14 show bar plots of posterior MIP values presented here.

| (A) Effect Class | D1 | D2 | D3 | D4 | (B) Effect Class | D1 | D2 | D3 | D4 |
| --- | --- | --- | --- | --- | --- | --- | --- | --- | --- |
| theta | 1.0000 | 1.0000 | 1.0000 | 1.0000 | theta | 1.0000 | 1.0000 | 1.0000 | 1.0000 |
| fixed: D0 | 0.0179 | 0.4630 | 0.6074 | 0.6699 | fixed: D0 | 0.0180 | 0.4590 | 0.6319 | 0.5172 |
| random: batch | 0.0772 | 0.8523 | 1.0000 | 0.9999 | random: batch | 0.0708 | 0.8332 | 0.9969 | 0.9979 |
|  |  |  |  |  | random: *Mx1* diplo | 0.0345 | 0.5515 | 1.0000 | 1.0000 |
| additive | 0.0141 | 0.0494 | 0.9779 | 0.9980 | additive | 0.0120 | 0.0558 | 0.3835 | 0.4305 |
| maternal | 0.0082 | 0.0713 | 0.7446 | 0.7400 | maternal | 0.0068 | 0.0671 | 0.6138 | 0.5272 |
| inbred (overall) | 0.3995 | 0.0164 | 0.9584 | 0.9992 | inbred (overall) | 0.3977 | 0.0238 | 0.6696 | 0.8789 |
| inbred | 0.1504 | 0.2024 | 0.3050 | 0.3504 | inbred | 0.1420 | 0.2068 | 0.2737 | 0.3334 |
| symmetric epistatic | 0.2309 | 0.3305 | 0.6674 | 0.9598 | symmetric epistatic | 0.2195 | 0.3642 | 0.5561 | 0.7668 |
| asymmetric epistatic | 0.0366 | 0.1193 | 0.8197 | 0.9662 | asymmetric epistatic | 0.0318 | 0.1176 | 0.6825 | 0.8511 |
| female (overall) | 0.0904 | 0.1718 | 0.1719 | 0.3135 | female (overall) | 0.0876 | 0.1762 | 0.2294 | 0.3394 |
| sex-by-additive | 0.3140 | 0.1993 | 0.3174 | 0.6220 | sex-by-additive | 0.3051 | 0.1981 | 0.2859 | 0.4884 |
| sex-by-maternal | 0.0738 | 0.3212 | 0.5550 | 0.3697 | sex-by-maternal | 0.0662 | 0.3194 | 0.4769 | 0.2939 |
| female inbred (overall) | 0.0237 | 0.0428 | 0.1012 | 0.0943 | female inbred (overall) | 0.0237 | 0.0432 | 0.0913 | 0.0889 |
| sex-by-inbred | 0.2527 | 0.2325 | 0.4448 | 0.3981 | sex-by-inbred | 0.2466 | 0.2345 | 0.3658 | 0.3345 |
| sex-by-symmetric epistatic | 0.1399 | 0.1694 | 0.3228 | 0.5873 | sex-by-symmetric epistatic | 0.1317 | 0.1692 | 0.3028 | 0.4359 |
| sex-by-asymmetric epistatic | 0.2016 | 0.3683 | 0.3682 | 0.4037 | sex-by-asymmetric epistatic | 0.1924 | 0.3719 | 0.3387 | 0.3296 |

**Figure S15.**
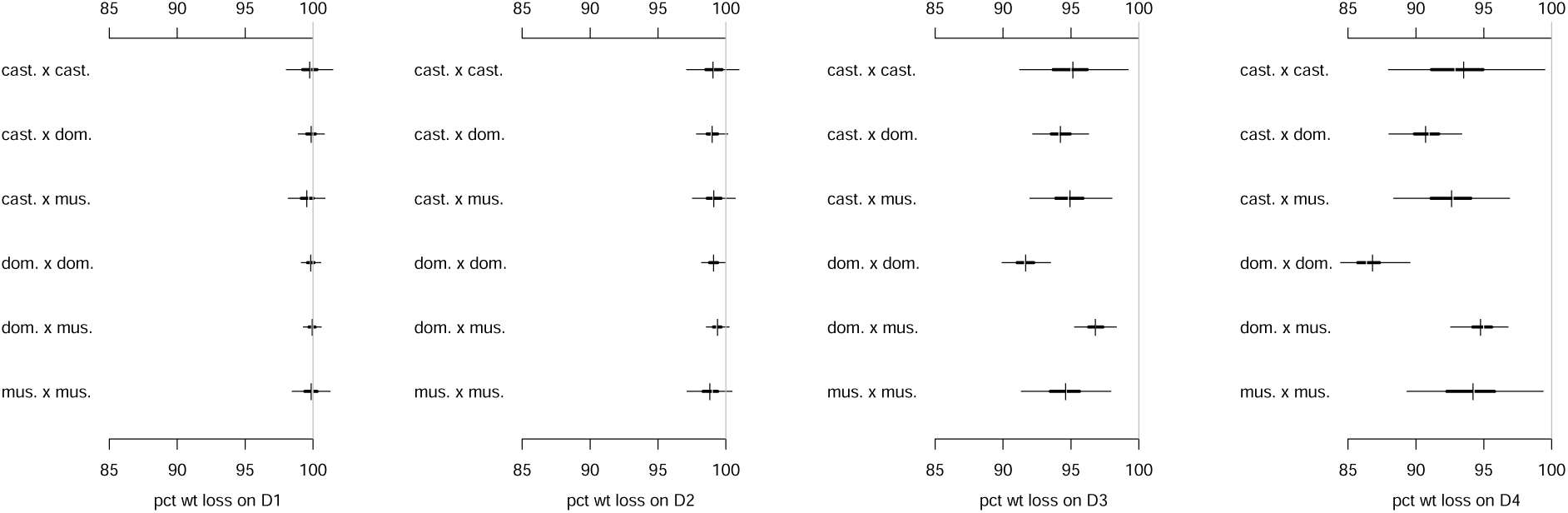
Timecourse of posterior predictive estimates of for *Mx1* diplotype mean effects across four days post-infection. From left to right, predictive estimates of IAV-induced weight change are provided for D1, D2, D3 and D4 p.i.

**Figure S16.**
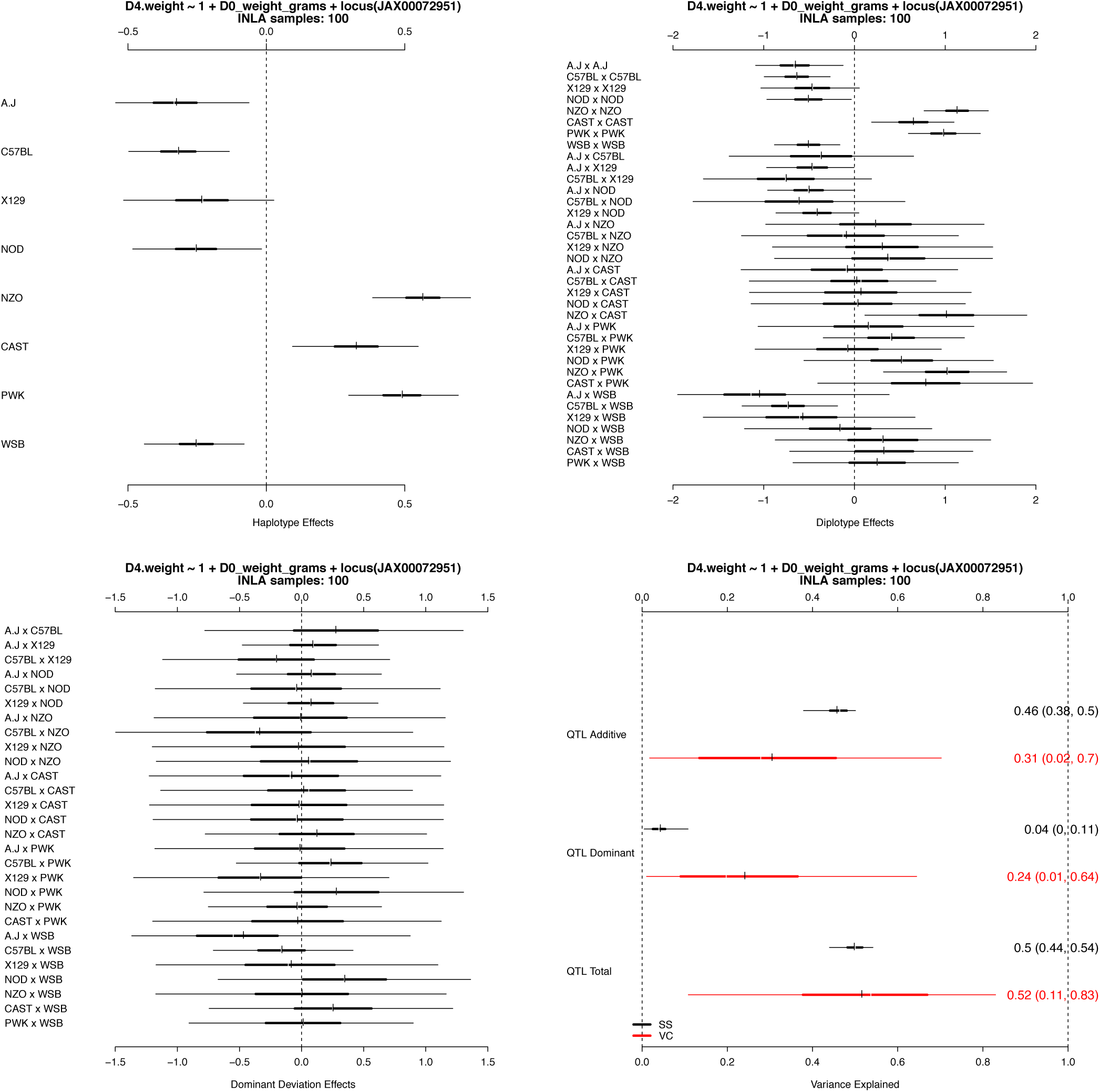
*Mx1* effects and proportion of variance explained on weight loss in the pre-CC. [Top Left] Additive *Mx1* haplotype effects in pre-CC mice (n=155) infected with IAV (PR8) at day 4 post-infection. [Top Right] Bayesian credible intervals of the posterior predictive distribution of mice with *Mx1* diplotype. [Bottom Left] Dominance deviation *Mx1* effects in the pre-CC mice. Wide intervals are indicative of the low levels of heterozygosity observed in the mostly inbred pre-CC mice. [Bottom Right] Bayesian credible intervals of the posterior predictive distribution of the proportion of the phenotypic variance explained by the *Mx1* additive effects, dominant effects and the combination of both effects. Black intervals (VC) are based on the posterior samples of the variance components. Red intervals (SS) are based on the posterior samples of sums of squares estimators. SS estimators are (1) not dependent on variance component estimates, which tend to be disperse, and (2) are based on the observed strains and crosses, rather than all possible strains/crosses, with (1) and (2) resulting in much narrower confidence intervals.

**Figure S17.**
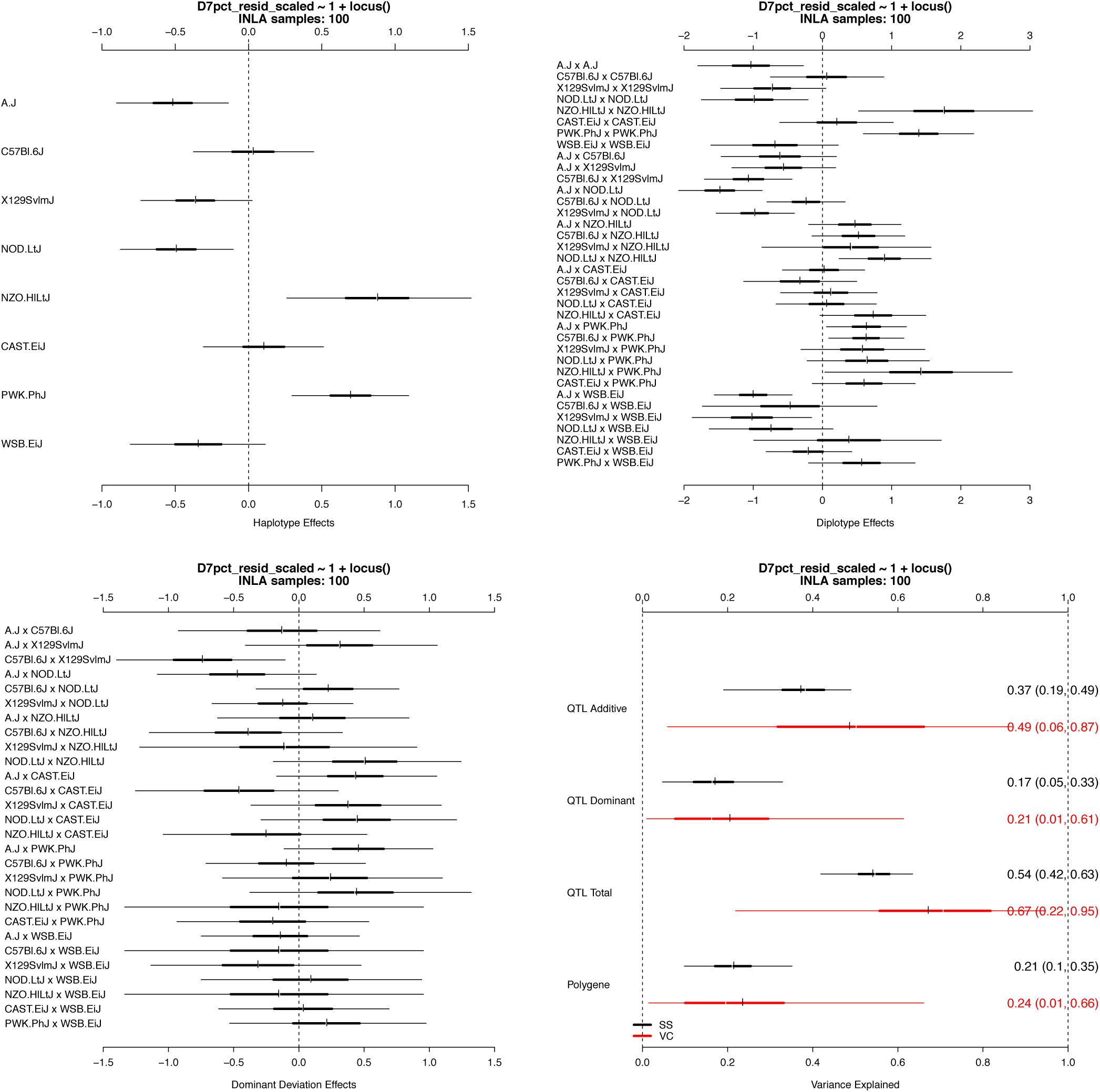
*Mx1* allele effects and proportion of variance explained on weight loss in the CC-RIX. [Top Left] Additive *Mx1* haplotype effects in CC-RIX mice (n=1,402) infected with IAV (CA04) at day 7 post-infection. [Top Right] Bayesian credible intervals of the posterior predictive distribution of mice with *Mx1* diplotype. [Bottom Left] Dominance deviation *Mx1* effects in the CC-RIX mice. Narrower intervals compared to observed in pre-CC is indicative of the high levels of heterozygosity CC-RIX mice. [Bottom Right] Bayesian credible intervals of the posterior predictive distribution of the proportion of the phenotypic variance explained by the *Mx1* additive effects, dominant effects, the combination of both *Mx1* effects, and cumulative effects of other loci captured in the relationship matrix. In the CC-RIX, the relationship matrix models the expected increased phenotypic correlation between CC-RIX mice that share a single CC parent. Black intervals (VC) are based on the posterior samples of the variance components. Red intervals (SS) are based on the posterior samples of sums of squares estimators. SS estimators are (1) not dependent on variance component estimates, which tend to be disperse, and (2) are based on the observed strains and crosses, rather than all possible strains/crosses, with (1) and (2) resulting in much narrower confidence intervals.

**Table S4.** Details of populations analyzed for additive strain-specific effects in Figure 7.

| POPULATION | diallel | pre-CC | CC-RIX |
| --- | --- | --- | --- |
| Recombinant | no | yes | yes |
| Inbreds | yes (some) | yes | no |
| Heterozygotes | yes (most) | no | yes |
| Sexes | males & females | females | females |
| IAV used | H1N1 (PR8) | H1N1 (PR8) | H1N1 (CA04) |
| Post-infection day analyzed | day 4 | day 4 | day 7 |
| Model | Bayes Diallel | Diploffect | Diploffect |

**Table S5.** Table of CC-RIX lines used in this experiment. A total of 1,402 mice from 105 CC-RIX strains (about 13 mice per strain), originating from a sparse diallel cross of 65 CC lines, were assayed for weight loss on day 7 post-infection.

| CC-RIX ID | count | CC-RIX ID | count | CC-RIX ID | count |
| --- | --- | --- | --- | --- | --- |
| CC001xCC055 | 16 | CC020xCC008 | 17 | CC042xCC019 | 13 |
| CC001xCC074 | 11 | CC021xCC023 | 15 | CC042xCC025 | 5 |
| CC002xCC011 | 14 | CC021xCC032 | 3 | CC043xCC033 | 21 |
| CC002xCC021 | 15 | CC022xCC024 | 11 | CC043xCC037 | 27 |
| CC003xCC051 | 12 | CC022xCC044 | 3 | CC044xCC060 | 18 |
| CC003xCC062 | 18 | CC023xCC025 | 18 | CC045xCC044 | 11 |
| CC004xCC011 | 15 | CC024xCC023 | 15 | CC046xCC068 | 21 |
| CC004xCC012 | 12 | CC024xCC052 | 11 | CC049xCC036 | 8 |
| CC005xCC001 | 15 | CC025xCC028 | 20 | CC051xCC005 | 14 |
| CC005xCC040 | 15 | CC026xCC034 | 17 | CC051xCC009 | 14 |
| CC006xCC007 | 20 | CC026xCC042 | 7 | CC052xCC007 | 3 |
| CC006xCC039 | 3 | CC027xCC036 | 3 | CC052xCC014 | 15 |
| CC007xCC070 | 12 | CC027xCC045 | 14 | CC055xCC028 | 21 |
| CC008xCC009 | 3 | CC028xCC024 | 21 | CC056xCC033 | 20 |
| CC008xCC010 | 16 | CC028xCC030 | 18 | CC057xCC052 | 16 |
| CC008xCC018 | 10 | CC029xCC027 | 16 | CC058xCC022 | 17 |
| CC009xCC040 | 12 | CC029xCC071 | 6 | CC059xCC065 | 4 |
| CC010xCC015 | 21 | CC030xCC023 | 23 | CC060xCC006 | 15 |
| CC010xCC060 | 9 | CC030xCC061 | 17 | CC061xCC025 | 16 |
| CC011xCC032 | 13 | CC032xCC013 | 18 | CC061xCC026 | 16 |
| CC011xCC042 | 15 | CC032xCC017 | 15 | CC061xCC039 | 15 |
| CC012xCC032 | 12 | CC033xCC046 | 18 | CC062xCC046 | 3 |
| CC012xCC038 | 20 | CC033xCC068 | 21 | CC062xCC072 | 3 |
| CC013xCC041 | 28 | CC034xCC016 | 10 | CC063xCC001 | 3 |
| CC014xCC059 | 4 | CC035xCC071 | 14 | CC063xCC002 | 7 |
| CC015xCC018 | 12 | CC036xCC051 | 16 | CC065xCC010 | 10 |
| CC015xCC059 | 15 | CC037xCC046 | 12 | CC065xCC072 | 1 |
| CC016xCC038 | 18 | CC037xCC057 | 19 | CC068xCC043 | 9 |
| CC016xCC061 | 13 | CC038xCC013 | 14 | CC070xCC003 | 15 |
| CC017xCC004 | 13 | CC039xCC020 | 14 | CC071xCC053 | 12 |
| CC017xCC041 | 16 | CC040xCC003 | 16 | CC072xCC063 | 4 |
| CC018xCC009 | 19 | CC040xCC015 | 13 | CC072xCC075 | 9 |
| CC018xCC065 | 16 | CC041xCC012 | 12 | CC074xCC058 | 7 |
| CC019xCC002 | 19 | CC041xCC016 | 18 | CC075xCC021 | 3 |
| CC019xCC004 | 16 | CC042xCC017 | 14 | CC075xCC035 | 14 |

**Table S6.**
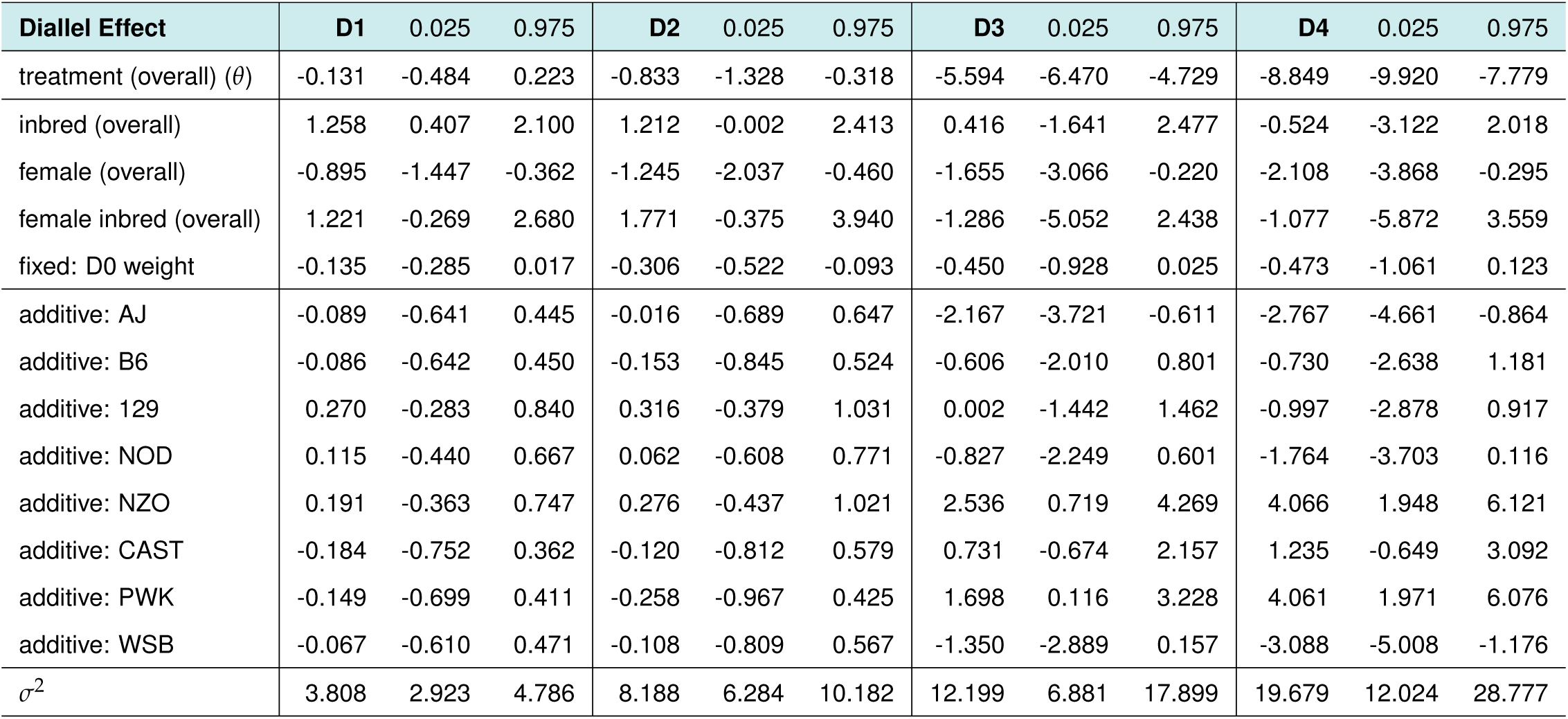
Summary of overall and additive effects from Figures 4A, S7A, S8A, S9A, and S10A. Posterior median and 95% HPD intervals are provided for D1 through D4 p.i.

| Diallel Effect | D1 | 0.025 | 0.975 | D2 | 0.025 | 0.975 | D3 | 0.025 | 0.975 | D4 | 0.025 | 0.975 |
| --- | --- | --- | --- | --- | --- | --- | --- | --- | --- | --- | --- | --- |
| treatment (overall) ($\theta$) | -0.131 | -0.484 | 0.223 | -0.833 | -1.328 | -0.318 | -5.594 | -6.470 | -4.729 | -8.849 | -9.920 | -7.779 |
| inbred (overall) | 1.258 | 0.407 | 2.100 | 1.212 | -0.002 | 2.413 | 0.416 | -1.641 | 2.477 | -0.524 | -3.122 | 2.018 |
| female (overall) | -0.895 | -1.447 | -0.362 | -1.245 | -2.037 | -0.460 | -1.655 | -3.066 | -0.220 | -2.108 | -3.868 | -0.295 |
| female inbred (overall) | 1.221 | -0.269 | 2.680 | 1.771 | -0.375 | 3.940 | -1.286 | -5.052 | 2.438 | -1.077 | -5.872 | 3.559 |
| fixed: D0 weight | -0.135 | -0.285 | 0.017 | -0.306 | -0.522 | -0.093 | -0.450 | -0.928 | 0.025 | -0.473 | -1.061 | 0.123 |
| additive: AJ | -0.089 | -0.641 | 0.445 | -0.016 | -0.689 | 0.647 | -2.167 | -3.721 | -0.611 | -2.767 | -4.661 | -0.864 |
| additive: B6 | -0.086 | -0.642 | 0.450 | -0.153 | -0.845 | 0.524 | -0.606 | -2.010 | 0.801 | -0.730 | -2.638 | 1.181 |
| additive: 129 | 0.270 | -0.283 | 0.840 | 0.316 | -0.379 | 1.031 | 0.002 | -1.442 | 1.462 | -0.997 | -2.878 | 0.917 |
| additive: NOD | 0.115 | -0.440 | 0.667 | 0.062 | -0.608 | 0.771 | -0.827 | -2.249 | 0.601 | -1.764 | -3.703 | 0.116 |
| additive: NZO | 0.191 | -0.363 | 0.747 | 0.276 | -0.437 | 1.021 | 2.536 | 0.719 | 4.269 | 4.066 | 1.948 | 6.121 |
| additive: CAST | -0.184 | -0.752 | 0.362 | -0.120 | -0.812 | 0.579 | 0.731 | -0.674 | 2.157 | 1.235 | -0.649 | 3.092 |
| additive: PWK | -0.149 | -0.699 | 0.411 | -0.258 | -0.967 | 0.425 | 1.698 | 0.116 | 3.228 | 4.061 | 1.971 | 6.076 |
| additive: WSB | -0.067 | -0.610 | 0.471 | -0.108 | -0.809 | 0.567 | -1.350 | -2.889 | 0.157 | -3.088 | -5.008 | -1.176 |
| $\sigma^2$ | 3.808 | 2.923 | 4.786 | 8.188 | 6.284 | 10.182 | 12.199 | 6.881 | 17.899 | 19.679 | 12.024 | 28.777 |

**Table S7.** Summary of overall, additive, and *Mx1* effects from Figures S11A, S12A, S13A, and S14A. Posterior median and 95% HPD intervals are provided for D1 through D4 p.i.

| Diallel Effect | D1 | 0.025 | 0.975 | D2 | 0.025 | 0.975 | D3 | 0.025 | 0.975 | D4 | 0.025 | 0.975 |
| --- | --- | --- | --- | --- | --- | --- | --- | --- | --- | --- | --- | --- |
| treatment (overall) ($\theta$) | -0.193 | -1.128 | 0.707 | -0.925 | -1.991 | 0.113 | -5.464 | -7.675 | -3.102 | -7.986 | -11.202 | -4.147 |
| inbred (overall) | 1.296 | 0.313 | 2.276 | 1.385 | 0.017 | 2.738 | 1.945 | -0.417 | 4.282 | 1.046 | -1.971 | 4.101 |
| female (overall) | -0.894 | -1.435 | -0.356 | -1.251 | -2.039 | -0.468 | -1.681 | -3.061 | -0.333 | -2.105 | -3.905 | -0.339 |
| female inbred (overall) | 1.224 | -0.231 | 2.705 | 1.765 | -0.364 | 3.931 | -1.221 | -4.945 | 2.312 | -1.018 | -5.663 | 3.654 |
| fixed: D0 weight | -0.137 | -0.289 | 0.012 | -0.313 | -0.528 | -0.099 | -0.548 | -0.996 | -0.096 | -0.576 | -1.155 | -0.016 |
| additive: AJ | -0.094 | -0.679 | 0.489 | -0.015 | -0.718 | 0.691 | -1.147 | -2.589 | 0.112 | -0.854 | -2.819 | 0.549 |
| additive: B6 | -0.086 | -0.664 | 0.501 | -0.145 | -0.867 | 0.567 | 0.190 | -0.980 | 1.352 | 0.401 | -1.017 | 1.891 |
| additive: 129 | 0.272 | -0.327 | 0.873 | 0.323 | -0.404 | 1.076 | 0.457 | -0.695 | 1.663 | 0.174 | -1.299 | 1.600 |
| additive: NOD | 0.113 | -0.453 | 0.722 | 0.067 | -0.639 | 0.799 | -0.046 | -1.221 | 1.130 | -0.148 | -1.769 | 1.266 |
| additive: NZO | 0.167 | -0.510 | 0.859 | 0.232 | -0.576 | 1.068 | 0.635 | -0.737 | 2.279 | 0.447 | -1.221 | 2.979 |
| additive: CAST | -0.137 | -0.935 | 0.619 | -0.048 | -0.942 | 0.831 | 0.267 | -1.225 | 1.866 | 0.230 | -1.545 | 2.277 |
| additive: PWK | -0.162 | -0.857 | 0.506 | -0.303 | -1.146 | 0.493 | 0.105 | -1.273 | 1.578 | 0.523 | -1.124 | 3.116 |
| additive: WSB | -0.072 | -0.653 | 0.501 | -0.111 | -0.836 | 0.605 | -0.514 | -1.811 | 0.650 | -0.909 | -2.962 | 0.533 |
| *Mx1*: *cast* × *cast* | -0.043 | -1.555 | 1.470 | -0.016 | -1.712 | 1.652 | 0.511 | -2.990 | 4.202 | 1.257 | -3.773 | 6.904 |
| *Mx1*: *cast* × *dom* | 0.057 | -1.077 | 1.193 | -0.089 | -1.397 | 1.183 | -0.311 | -3.008 | 2.256 | -1.287 | -5.596 | 2.401 |
| *Mx1*: *cast* × *mus* | -0.238 | -1.572 | 1.028 | 0.036 | -1.409 | 1.521 | 0.317 | -2.619 | 3.433 | 0.451 | -3.933 | 5.062 |
| *Mx1*: *dom* × *dom* | 0.028 | -1.076 | 1.217 | 0.015 | -1.253 | 1.290 | -2.831 | -5.872 | -0.131 | -5.274 | -9.857 | -0.779 |
| *Mx1*: *dom* × *mus* | 0.135 | -0.897 | 1.213 | 0.290 | -0.870 | 1.522 | 2.198 | -0.207 | 4.743 | 2.624 | -1.118 | 6.299 |
| *Mx1*: *mus* × *mus* | 0.061 | -1.251 | 1.445 | -0.234 | -1.817 | 1.213 | 0.016 | -3.171 | 3.295 | 1.948 | -2.574 | 7.167 |
| $\sigma^2$ | 3.797 | 2.913 | 4.769 | 8.190 | 6.328 | 10.268 | 11.270 | 6.644 | 16.627 | 19.296 | 12.210 | 27.397 |

**Table S8.**
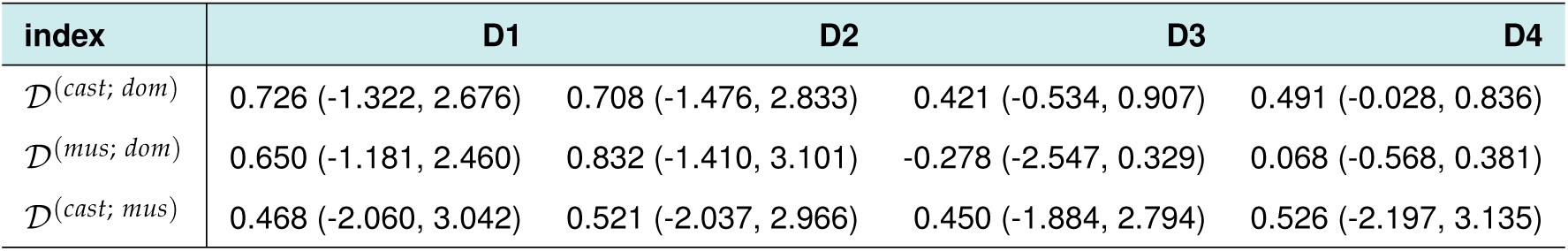
Summary of dominance index for *Mx1* from *cast* or *mus*, combined with *Mx1* from *dom* in our diallel study. Posterior mode and and 80% HPD intervals are provided from D1 through D4 p.i. The posterior mode is obtained from the value of 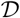 that has the maximum posterior density within the range [−1, 2], via the `density` function from the `stats` package in R.

| index | D1 | D2 | D3 | D4 |
| --- | --- | --- | --- | --- |
| $\mathcal{D}^{(cast; dom)}$ | 0.726 (-1.322, 2.676) | 0.708 (-1.476, 2.833) | 0.421 (-0.534, 0.907) | 0.491 (-0.028, 0.836) |
| $\mathcal{D}^{(mus; dom)}$ | 0.650 (-1.181, 2.460) | 0.832 (-1.410, 3.101) | -0.278 (-2.547, 0.329) | 0.068 (-0.568, 0.381) |
| $\mathcal{D}^{(cast; mus)}$ | 0.468 (-2.060, 3.042) | 0.521 (-2.037, 2.966) | 0.450 (-1.884, 2.794) | 0.526 (-2.197, 3.135) |

**Algorithm 1:**
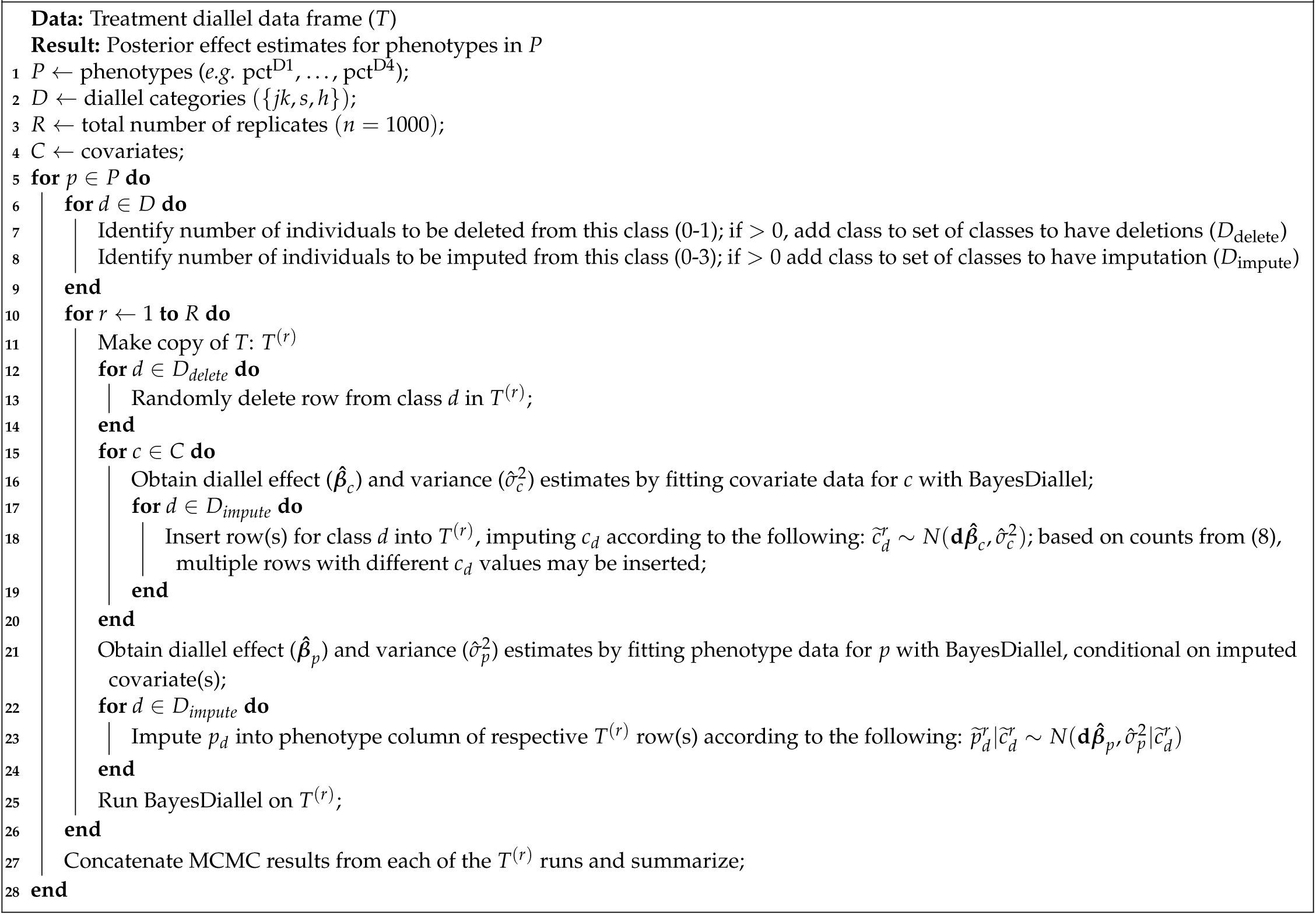
Multiple Imputation Matched Quartets (MIMQ)

